# Collective protection drives human gut microbiota response to amoxicillin treatment

**DOI:** 10.64898/2026.04.24.720353

**Authors:** Paul Lubrano, Mélanie Magnan, Caroline Steiner, André Birgy, Benoît Chassaing, Mélanie Deschasaux-Tanguy, Arnaud Gutierrez, Hugo C. Barreto, Claire Hobson, Sophie Magréault, Vincent Jullien, Mathilde Lescat, Olivier Tenaillon

## Abstract

The gut microbiota is central to human health, contributing to nutrient processing, metabolite production and protection against pathogens. Yet it is also an unintended target of antibiotic treatments, particularly after oral administration. Antibiotic exposure can therefore disrupt community structure, leading to dysbiosis and promoting the selection of resistant bacteria. Although studies in patients and animal models have shown that these effects vary markedly between individuals, host-related factors have made it difficult to isolate the specific contribution of the microbiota itself. Here, we used a controlled *in vitro* gut model (MBRA) to examine how 16 human gut microbiotas from the NutriNet-Santé cohort respond to the widely prescribed β-lactam amoxicillin (AMX). By combining dense temporal sampling, 16S amplicon sequencing and mass spectrometry, we observed highly heterogeneous response trajectories, ranging from near-stable communities to strong but reversible disruptions. These differences were not only reflected in the final magnitude of perturbation, but also in the timing, pace and recovery of microbiota change during treatment. Initial community composition partly structured these responses, as *Lachnospiraceae*/*Bacteroidaceae* ratio strongly correlated with perturbation. Dynamic quantification of AMX further showed that microbiotas differed in their capacity to deplete the drug over time, thereby altering the duration of exposure above critical concentration thresholds. Supplementation with clavulanic acid that inhibits β-lactamases confirmed that this process was largely mediated by β-lactamase activity. Finally, microbiotas displaying rapid AMX depletion showed reduced selection of resistant *Enterobacteriaceae*. Together, our results indicate that both initial community composition and the temporal dynamics of antibiotic inactivation jointly determine microbiota resilience and resistance selection.

## Introduction

The remarkable gains in public health brought by antibiotics use are now under strain from the rise and spread of resistant bacteria. Once largely confined to hospitals, resistance has become a community-wide concern^1,2^. This is based on the fact that, for instance in France, around 90% of antibiotic use occurs outside hospitals^3–5^. Moreover, most community prescriptions are taken orally, a route that amplifies the collateral impact on the body’s largest microbial ecosystem—the intestinal microbiota^6–8^. Each course of treatment perturbs community members, regardless of their roles in the underlying infection. For example, *Escherichia coli,* a pathobiont, is estimated to be exposed to antibiotics as a commensal in more than 90% of treatments^9^.

Repeated antibiotic exposures have two broad consequences. First, they disrupt the composition of the gut microbiota. This disturbance, commonly termed “dysbiosis”, can persist for weeks or even months after treatment^10–13^, with a range of downstream detrimental consequences. Indeed, the gut microbiota provides essential services to its host, ranging from immune system modulation^14^, nutrition^15^ or pathogen colonisation resistance^16^. Second, antibiotic exposure may favor the emergence and spread of antibiotic resistance. The diversity of gut microbiotas suits *de novo* acquisition of resistance or horizontal transfer of resistance genes between species^17–19^. The growing prominence of *E. coli* as a public-health threat illustrates this dynamic: once mostly antibiotic-sensitive and easy to treat, it has been associated with over one million deaths worldwide in 2019^20^, a substantial share linked to antibiotic resistance.

Yet, while the overall impact of antibiotics on the gut microbiota is clear, the dynamics within and between individuals are more intricate. Studies examining identical treatments across different individuals revealed considerable variability in the magnitude of microbiota disruption^6,21–23^. Likewise, although simple models predict that antibiotics should select for resistant strains and eliminate sensitive ones, sensitive strains have been observed to persist—and in some cases even outcompete resistant ones^24–26^. Understanding the sources of this heterogeneity could offer new avenues to limit the spread of resistance and reduce antibiotic-mediated gut microbiota dysbiosis.

Several factors may be at play. Some relate to the host, including immune function^14,27^ and the metabolization of antibiotics^28^. The composition of the microbiota may also influence how a community responds to treatment^22,29–31^ and, in turn, how resistance is selected. Indeed, members of the microbiota vary in their intrinsic or acquired resistance to antibiotics, so community composition alone may drive differential responses^32^. Furthermore, some bacteria can actively degrade antibiotics through specific enzymes, reducing effective concentrations and potentially limiting both disruption and selection for resistance^7^. Since intestinal microbial communities can be shaped by various factors, including diet^33,34^ and probiotics^35,36^, they are amenable to targeted intervention. Investigating the extent to which gut microbiotas modulates the effects of antibiotics could therefore open avenues for mitigating these effects.

To this end, *in vitro* modelling systems provide a useful complement to clinical and animal studies. While human trials and murine models remain essential, they make it difficult to disentangle host effects versus microbiota-driven processes, lack reproducibility across individuals with distinct microbiotas, and offer only indirect access to gut dynamics (fecal sampling)^37,38^. These constraints have led to the development of *in vitro* models capable of long-term maintenance of complex microbial communities under controlled anaerobic conditions^39–41^. Although such systems remove host variability and enable direct sampling and high-throughput analyses, their main limitation is their simplified representations of the gut ecosystem. Nonetheless, they provide an informative framework for studying the contribution of gut microbiotas to antibiotic response.

Among these systems, the MiniBioReactor Array (MBRA) offers a useful balance between throughput and ecological stability^42,43^. Built around a 48-chamber anaerobic chemostat, it allows microbial communities to be stably maintained for up to two weeks. This duration accommodates both a stabilisation phase, a treatment phase that mimics repeated antibiotic exposure (typically with two doses per day), and a final recovery phase. Continuous flow conditions reproduce aspects of gut dynamics, including concentration gradients experienced during oral treatment In this study, we used this MBRA system to examine how 16 distinct microbiotas isolated from healthy donors respond to the β-lactam amoxicillin. Our choice was motivated by the fact that β-lactams are among the most widely prescribed antibiotics in France^44^. Among them, amoxicillin and the amoxicillin–clavulanic acid combination are first-line therapies for common respiratory and otorhinolaryngological infections across age groups^45^. Their favorable safety profile and frequent administration, particularly during early childhood, make these aminopenicillins one of the most prevalent antibiotic exposures experienced by the human microbiota. With our study, we observed a wide range of perturbation profiles amongst the 16 microbiotas, confirming their strong influence on AMX-mediated perturbation irrespective of host-specific factors. Amongst heterogeneity determinants, we show that the *Lachnospiraceae*/*Bacteroidaceae* ratio of a microbiota before treatment correlates with AMX perturbation. Using the β-lactamase inhibitor clavulanic acid and dynamic quantification of AMX with mass spectrometry, we further highlight that the key factor determining robustness to perturbation is the capacity of communities to deplete amoxicillin during treatment and protect their most sensitive members as well as preventing selection of AMX-resistant *Enterobacteriaceae*.

## Results

### Description of chosen donors, MBRA system, sequencing and data analysis

We first obtained gut microbiotas from fecal samples of the Etude NutriNet santé cohort donors. This cohort includes participants over 15 years old representing the general French population^46^. A total of 103 participants were selected based on their fiber consumption (see Material and methods). Each donor immediately stored its feces in tube upon production. Feces were then delivered within 2 hours to our laboratories and were aliquoted in an anaerobic chamber. We chose 16 different donors from the cohort, based on the diversity of their diet, especially in term of fiber content (Table S1). Fecal samples were filtered and inoculated in MBRA chambers. A total of six chambers were inoculated per donor.

Our experimental setup was the following (Fig.1a): microbiotas were first cultivated for a total of 5 days to stabilise them in the chambers^33,42^. At Day 5, three chambers per donor were treated with 100 µg/mL of AMX (see Material and methods). The three remaining chambers were treated with the AMX solvent, water. Inoculation of AMX or water was done directly in the chambers with the help of a syringe. To simulate oral treatments in human communities, AMX or water doses were added to each chamber twice a day, first in the morning and then 8 hours later. This led to an 8/16 hours treatment regimen. After five days, we stopped the treatments and pursued the cultivation for five more days. This period allowed us to analyse the capacity of the microbiotas to recover after the treatment.

During the experiment, we sampled the chambers in the morning at defined days (3/5/6/8/10/11/13/15). During treatment (Day 5 - Day10), the sampling was done in the morning right before the addition of AMX or water. We extracted genomic DNA and performed 16S rRNA amplicon sequencing. Using QIIME2^47^ and the Greengenes2 database^48^, each read was attributed to a bacterial family (taxonomic rank 5). We chose this taxonomic rank to reduce noise from the data, as our average read count per sample neighboured 18 000 (Fig. S1). Read counts were converted to frequencies which we used for our analysis (Fig. S2). Further processing of the data is described in the Material and methods section. We also monitored the total number of *Enterobacteriaceae* in each chamber using Drigalski selective agar^49^. Additional selective agar plates were supplemented with AMX (32 µg/mL) to estimate the proportion of AMX-resistant *Enterobacteriaceae* amongst the total population. Next, using this data, we investigated AMX perturbation on each of the 16 donors within our MBRA system.

### Gut microbiotas are variably perturbed by AMX

Treatment perturbation of each microbiota was evaluated using its composition on Day 5 as a reference (Fig. 1b). We relied on α-and β-diversity metrics to track changes (Supplementary note 1 – Fig. 1c, 1d, S3-6). While α-diversity scales with the number of equally represented bacterial families in a microbiota, β-diversity depends on compositional variations. Hence, both metrics are complementary to estimate dysbiosis (change in composition and microbiota equilibrium). Both β-diversity and α-diversity metrics were integrated in “perturbation curves” plotted across the experiment (Fig. 1e and Fig S7). Analysis of the resulting curves showed heterogeneity among donors as well as common perturbation patterns. For most AMX-treated chambers, perturbation was maximal on the first day of treatment (Day 6). We also observed a recovery after AMX removal, reminiscent of *in vivo* observations^37,50,51^.

**Fig 1.**
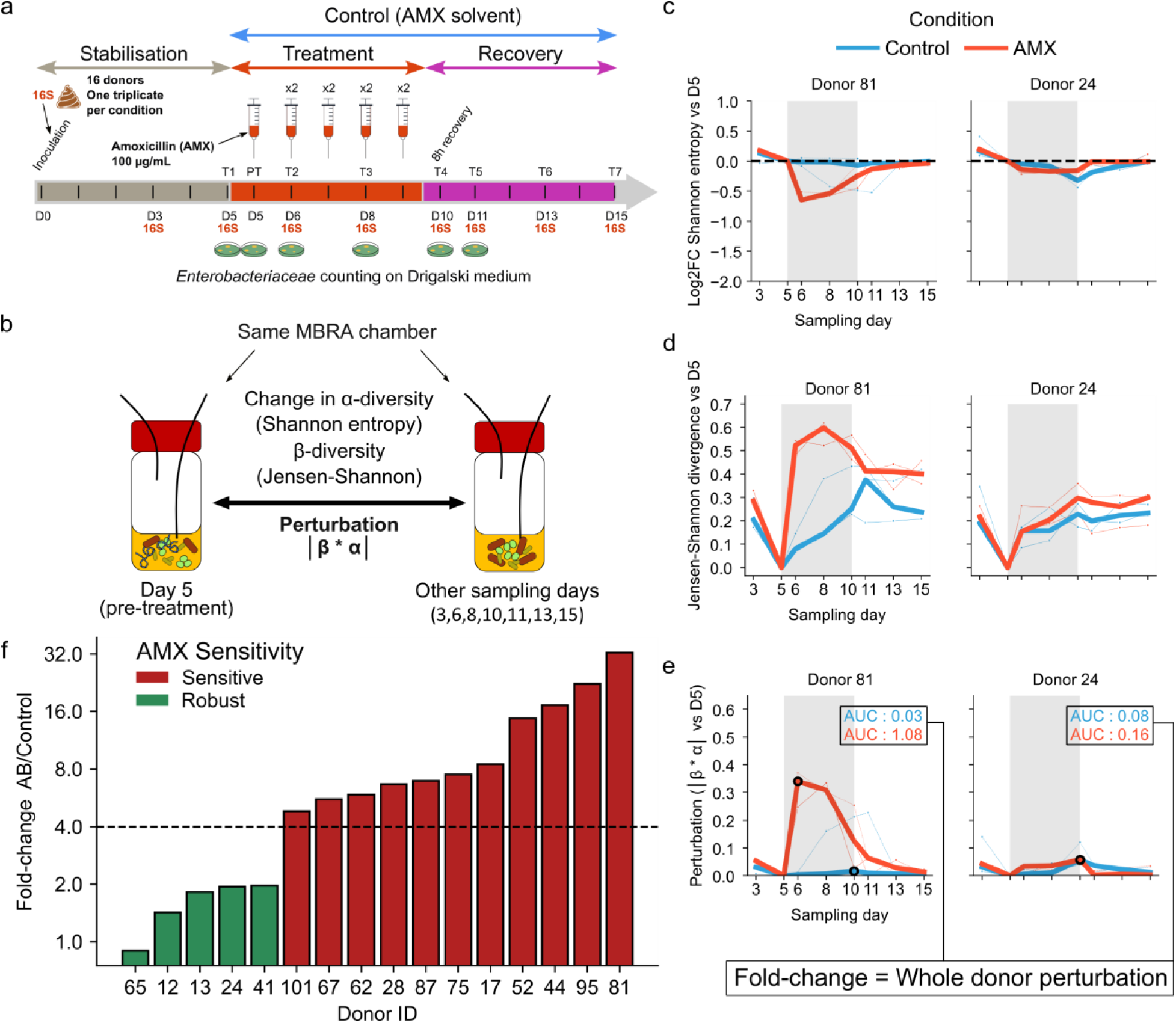
Adding amoxicillin to human gut microbiotas *in vitro* leads to different perturbation patterns. **a.** Representation of the experimental set-up using a Mini BioReactor Array (MBRA). A total of 16 donors microbiotas are extracted from feces and cultivated in 2 x 3 replicates in MBRA chambers. After 5 days of stabilisation, one triplicate is treated two times a day with amoxicillin (AMX), while the other receives only the AMX solvent (water). The treatment period lasts 5 days and is followed with a 5-days recovery period. Bacterial families in the chambers are monitored using 16S rRNA amplicon sequencing at indicated time points (red). *Enterobacteriaceae* are also plated on Drigalski medium with/without AMX to monitor resistance selection in this family. **b.** Representation of the metrics used to estimate microbiotas perturbation. For a given chamber (one replicate, one donor, one condition), perturbation is always estimated in relation to the chamber microbiota at Day 5 (just before treatment) and using both α and β-diversity indices. In the case of α-diversity, the value is obtained through the log2 Fold-change of the α-diversities. A perturbation score is also calculated with the absolute of the product of both α and β-diversities metrics. **c.** An example of α-diversity change (as Shannon entropy) in the chambers of two donors (81 and 24). The grey rectangle corresponds to the treatment window. Blue lines show change in the Control triplicate and red lines show change in the AMX-treated triplicate. Replicates are represented in thin lines, and median of the triplicates are represented in bold lines. **d.** Same as (c) but with β-diversity (Jensen-Shannon divergence) during the experiment. **e.** Same as (c) but with the perturbation score. Here, Area Under Curve (AUC) values are shown for the median curves. The fold-change of the control and AMX-treated median curves AUCs indicates the strength of a donor perturbation caused by the AMX treatment. Black dots represent the day of median maximum perturbation observed within the treatment window. **f.** Bar plot representing the strength of AMX perturbation across all the 16 donors of the panel. Donors are categorised between “AMX-Sensitive” and “AMX-Robust” depending on the value (sensitive: above 4).

We next used the Area Under Curve (AUC) of the perturbation curves within the treatment window to integrate treatment perturbations. This highlighted that some microbiotas were more dynamic than others, showing perturbation even in the Control condition. For example, the median AUC of Donor 12 Control chambers was of 0.15 while it was of 0.03 for Donor 52. Hence, to fully isolate AMX-only perturbation for each donor, AUCs of the AMX-treated chambers were contrasted to the ones of the Control chambers through a fold-change (FC - Fig. 1f). This FC varied from 0.89 (Donor 65) to 32.5 (Donor 81), confirming strong disparities of AMX responses in gut microbiotas of our 16-donor panel. This large repertoire of perturbations is a strong support for our initial hypothesis: AMX perturbation greatly varies between gut microbiotas, and that regardless of host-related factors. We observed a threshold separating Donor 41 and Donor 101 (FC respectively of 1.97 and 4.8). This allowed us to classify donors as « AMX Robust » or « AMX Sensitive ». Changing the taxonomic rank or including rarefaction did not impact our analysis (Supplementary note 2, Fig. S8-9). We next sought to understand what might have influenced AMX response differences amongst gut microbiotas of our panel.

### Pre-treatment microbiota composition does not correlate with AMX perturbation unlike *Lachnospiraceae*/*Bacteroidaceae* ratio

Previous studies have reported that bacterial families may be more vulnerable to antibiotic treatments than others. In various models, families like *Lachnospiraceae* or *Ruminococcaceae* have been shown to be highly sensitive to AMX^52–54^. Others, such as *Enterobacteriaceae* or *Bacteroideacaeae* are generally more robust to that antibiotic^55,56^, likely because these families contain known β-lactamases producers^57–60^. Therefore, we investigated how microbiota composition before treatment (Day 5) may be predictive of perturbation. Here, composition refers to the combined frequencies of bacterial families within each chamber.

First, we ensured that community composition was reproducible between chambers from the same donor at Day 5. We employed hierarchical clustering on a Jensen-Shannon divergence matrix (Fig. 2a), and observed robust clustering of chambers from the same donor. We noted exceptions linked with variations within the MBRA system (e.g *Enterococcaceae* expansion was observed in two of the six replicates of Donor 41 chambers). Excluding these variations, this highlighted that gut microbiotas from the same inoculum remain similar after a 5-days stabilisation period and confirmed the robustness of our system for correlation analyses.

**Fig 2.**
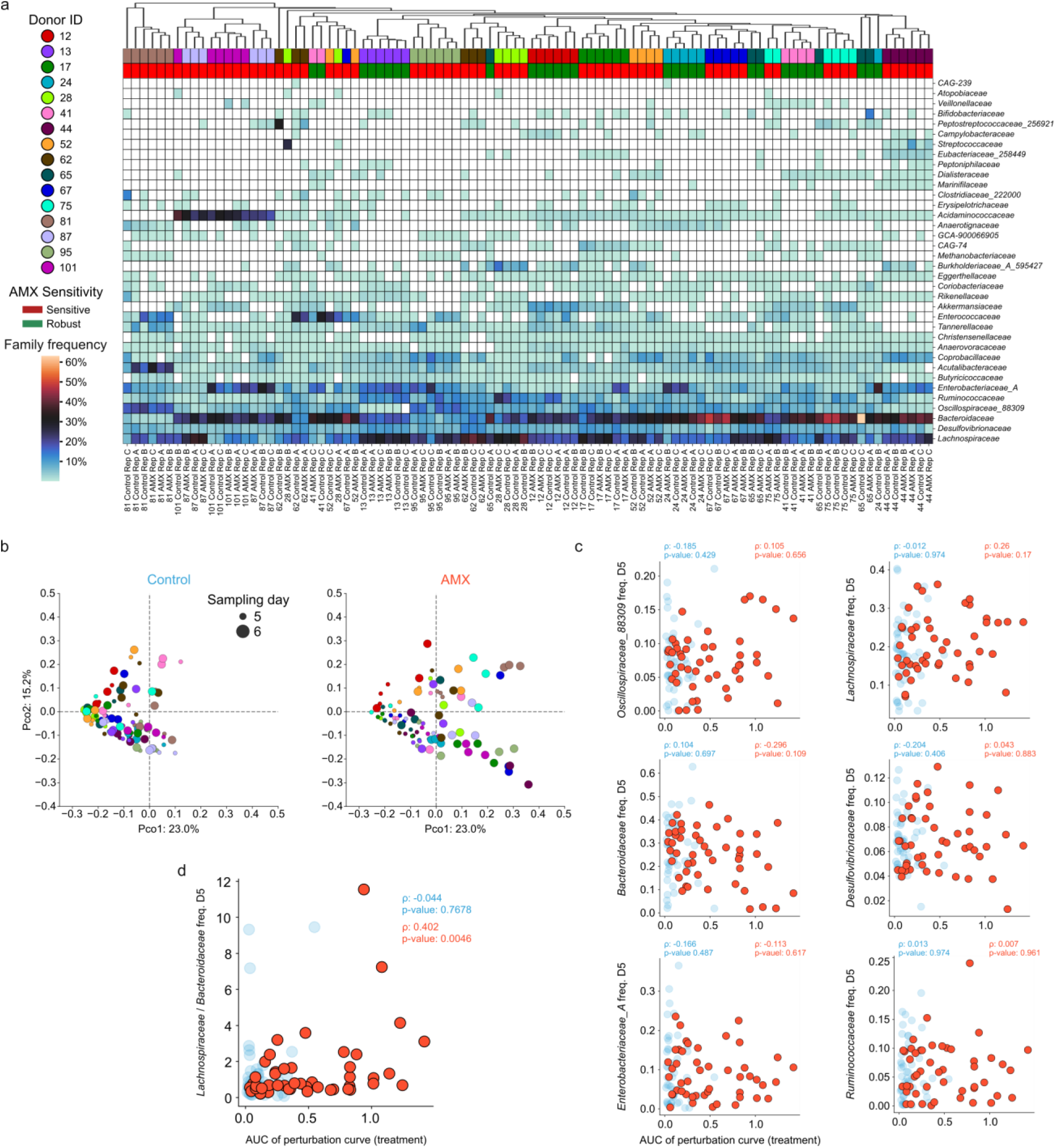
Pre-treatment microbiota composition does not correlate with perturbation unlike *Lachnospiraceae*/*Bacteroidaceae* ratio. **a.** Clustermap representing the microbiota composition of each chamber at Day 5, just before addition of AMX or water (Control condition). Hierarchical clustering of the microbiotas was performed using a Jensen-Shannon divergence matrix. Donor IDs are indicated in colors above the heatmap. Donor AMX sensitivity is also indicated above the heatmap, using the classification of Fig. 1. Heatmap cells are colored according to the frequency (%) of a bacterial family. **b.** Principal coordinate analysis of all microbiotas of this study. Microbiotas of all chambers at all sampling days were used to build a Jensen-Shannon divergence matrix. On the left are shown microbiotas in control chambers at Day 5 and Day 6. On the right, microbiotas treated with AMX are shown at Day 5 and Day 6. Dot size corresponds to the displayed sampling day. Dots are colored according to the Donor ID associated with the chamber. **c.** Frequencies of the six most abundant bacterial families at Day 5 in each chamber, plotted with their respective chambers perturbations during treatment. Each dot represents a chamber, with light blue dots corresponding to control chambers and red dots corresponding to AMX-treated chambers. Spearman (ρ) correlation coefficients and associated Benjamini-Hochberg corrected p-values are provided. **d**. Same plot as in **c.** but this time plotting the ratio of frequencies of *Bacteroidaceae* and *Lachnospiraceae*.

To this end, we used the perturbation value to each MBRA chamber, which is the AUC of its perturbation curve within the treatment window (see previous section). We then tested whether pre-treatment composition of each chamber correlated with perturbation, and found no strong and significant correlations (Supplementary note 3, Fig. S10-11). Data visualisation with a Principal Coordinates Analysis (PCOA) confirmed good clustering of chambers from the same donor based on their compositions. On that representation, close chambers from different donors at Day 5 scattered from each other at Day 6 (e.g Donors 95/24; 75/17 - Fig. 2b and S12). These approaches highlighted that microbiota composition before AMX treatment is not predictive of perturbation.

As we could not find clear patterns with whole microbiotas compositions, we questioned whether the frequency of individual families at Day 5 might be predictive of a chamber perturbation (Fig. 2c). We focused on the six most shared families across the donor panel: *Enterobacteriaceae*, *Bacteroidaceae*, *Lachnospiraceae*, *Desulfovibrionaceae*, *Oscillospiraceae_88309* and *Ruminococcaceae*. The prevalence of these families within the human gut microbiota has been observed across multiple studies^61^. None of these families frequencies showed a strong correlation with perturbation in AMX-treated chambers, including the abundant *Lachnospiraceae* and *Bacteroidaceae* (respective Spearman ρ = 0.26 and −0.296; p-values of 0.114 and 0.074). Nevertheless, we found a strong and significant correlation between the *Bacteroidaceae* to *Lachnospiraceae* frequency ratio and AMX perturbation (ρ =−0.402, p-value<0.05) (Fig 2d). This association was restricted to the AMX-treated chambers and indicates that, to a certain extent, microbiotas with higher frequencies of *Bacteroidaceae* than *Lachnospiraceae* may be more robust to AMX perturbation. This is coherent with regards to the known AMX-sensitivity of these families^52,53,55^.

Overall, it was challenging to predict the AMX response of a microbiota based on its composition before treatment. Therefore, to gain more insight on perturbation heterogeneity between the microbiotas in our MBRA system, we investigated AMX-responses of the 16 donors gut microbiotas during the treatment instead.

### Key AMX-sensitive bacterial families are stable within robust microbiotas

We tracked frequency changes of bacterial families within each chamber, using their Day 5 frequencies as a reference (Fig 3a). The resulting “family perturbation curves” represented how a family varies across the experiment and in each chamber. The AUC of these curves within the treatment window was used as a proxy to quantify the impact of treatment (Control or AMX). As frequencies were noisy for lowly abundant species (see Fig 2a for instance), we focused our analysis on the six most abundant bacterial families (see previously, Fig 3b). In some chambers, we used pseudocounts for families that disappeared during treatment (see Material and methods). For example, this was frequently observed for the AMX-sensitive *Lachnospiraceae* in AMX-treated chambers. Other families AUCs were also plotted (Fig. S13).

**Fig 3:**
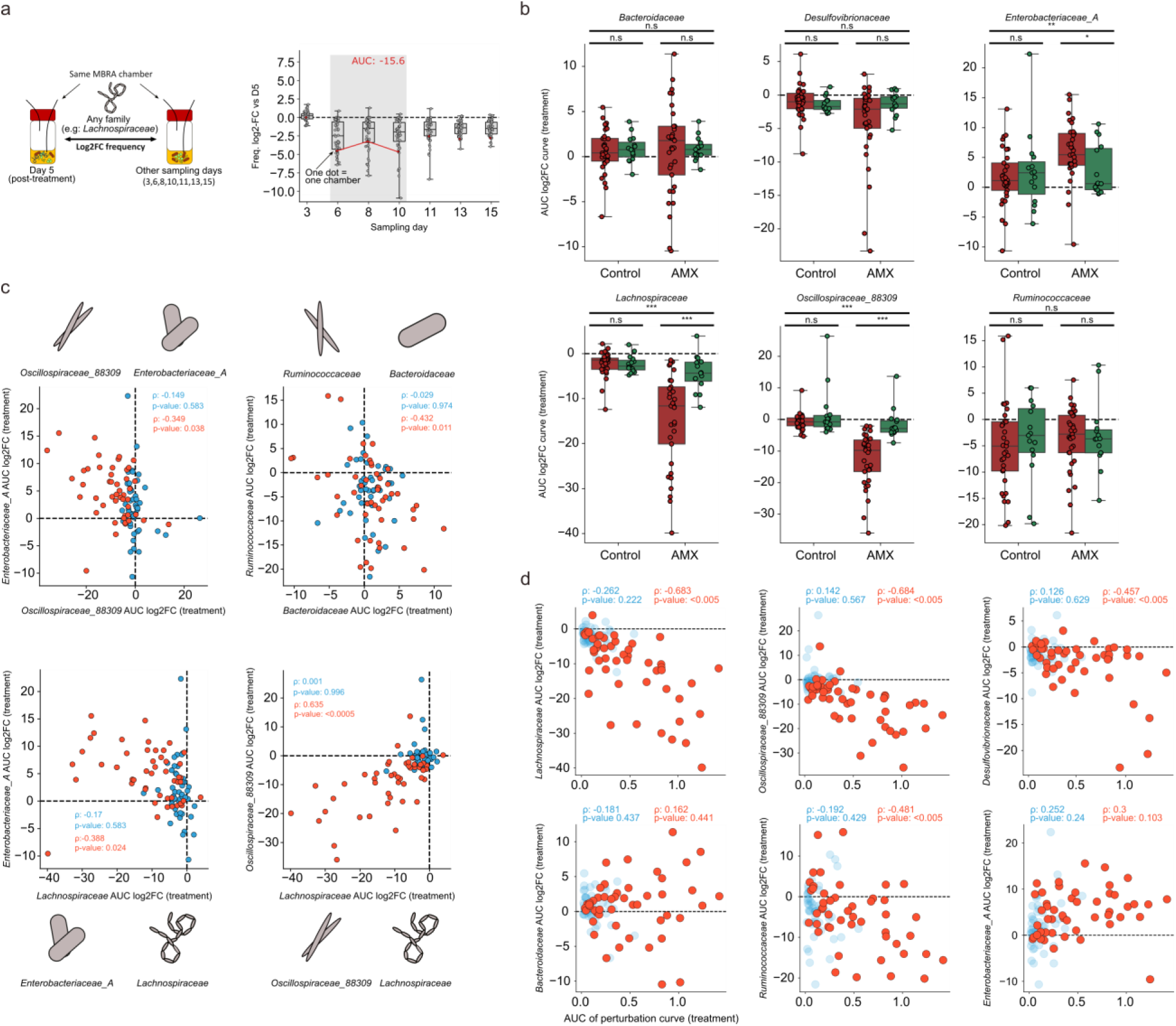
AMX robustness is linked with key bacterial families stabilisation. **a.** Schematisation of the method used to calculate the Area Under Curve (AUC) values displayed on this figure. For each bacterial family, its frequency change between each sampling day and the Day 5 frequency is calculated in each MBRA chamber. The AUC is then computed within the treatment window (in grey). **b.** AUC values of six major families in chambers either treated with AMX or untreated (control condition). In green are chambers of AMX-Robust donors, and in red chambers of AMX-Sensitive donors. Mann-Whitney p-values are used to display significance (n.s= non significant; * = p-value < 0.05; ** = p-value < 0.005; *** = p-value < 0.0005) when comparing the various groups (Robust vs Sensitive or Control vs AMX). Exact p-values and Cliff’s delta effect size are provided in the Data table (S21) **c.** Comparison between AUCs of two major bacterial families in each chamber. In red are chambers treated with AMX and in blue, Control chambers. Spearman (ρ) correlation coefficients are provided with their associated p-values (Benjamini-Hochberg corrected). **d.** AUC values of the six most abundant bacterial families, plotted with associated perturbation in their respective chambers during treatment. Each dot represents a chamber, with light blue dots corresponding to Control chambers and red dots corresponding to AMX-treated chambers. Spearman (ρ) correlation coefficients and associated Benjamini-Hochberg corrected p-values are provided. Exact p-values are provided in the Data table (S24).

To observe systemic trends, we grouped all AUCs of each family according to the sensitivity group of their donors (AMX-Sensitive vs AMX-Robust; see Fig. 1f), and the treatment used for their chambers (AMX vs Control). This revealed interesting patterns. First, when comparing Control and AMX conditions, we could highlight the behavior of each family in MBRA chambers (see Fig.3b – Control vs AMX tests). A notable example is that of the *Enterobacteriaceae*. The tendency of this family to increase in frequency in the Control MBRA chambers (Median AUCs of all Control chambers = 1.32) may reflect a fitness advantage in our system. Oppositely, *Lachnospiraceae* decreased in the Control MBRA chambers (Median AUCs of all Control chambers = −1.93), indicating that this family may have a fitness defect in our system irrespective of addition of AMX.

As expected, we observed that AMX had a large impact on the AUCs of the families. Both *Lachnospiraceae* (Control/AMX p-value<0.05 and Cliff’s delta=0.69) and *Oscillospiraceae_88309* (Control/AMX p-value<0.05, Cliff’s delta=0.75) were strongly affected, while *Enterobacteriaceae* benefited from the treatment (Control/AMX p-value<0.05, Cliff’s delta=−0.37). AMX sensitive families tended to recover from the treatment after its termination, increasing in frequency in most perturbed chambers (Fig S14 – Supplementary note 4).

As the microbiota composition results from a complex ecological equilibrium between its members, the loss of some abundant families during treatment may result in systemic modifications, destabilising the whole ecosystem. We explored these dynamics by comparing AUCs of families against each other within each chamber (Fig. 3c). Many family’s AUCs were significantly correlated in chambers treated with AMX. For example, losses of *Lachnospiraceae* were associated with an increase of *Enterobacterioceae* (Spearman ρ=−0.388, p-value=0.024), and similarly *Ruminococcaceae* and *Bacteroidaceae* abundance changes were negatively correlated (Spearman ρ=−0.432, p-value=0.011). Although these correlations may be coincidental and not reflect causality links, they suggest that the change in abundance of a family could lead to a chain of events that affect other families due to ecological interactions (e.g competition and cross feeding)^62^. Interestingly, we observed strong and significant correlations between AUCs of *Lachnospiraceae*/*Oscillospiraceae_88309* (ρ=−0.683 and −0684, both p-values <0.05) and AMX perturbations in chambers (Fig. 3d), which hint these families may have a key role in maintaining microbiotas equilibrium.

We finally looked at how families were affected according to the classification of their donors (see Fig.3b and S14, AMX-Robust vs AMX-Sensitive tests). Coherently, we overall found a systemic reduction of the perturbation of families from AMX-Robust donors. For instance, *Enterobacteriaceae* increase was strongly reduced (p-value<0.05, Cliff’s delta=−0.42) and *Bacteroidaceae* showed significant reduction of absolute abundance change (p-value<0.05, Cliff’s delta=−0.689 -Fig. S15). Most importantly, the effect was notable in the families *Oscillospiraceae_88309* and *Lachnospiraceae* (both p-value<0.05, Cliff’s delta=0.85 and 0.64). This highlighted that these two major AMX-sensitive families did not respond to AMX in gut microbiotas from robust donors.

The lack of response of a family to an antibiotic may come from two sources: genetic resistance, or protection from other members of the microbiota. AMX resistance can be caused by acquisition of new Penicillin Binding Proteins (PBP) and mutations in PBPs genes^63^, reduction of import^64,65^, AcrAB-TolC efflux^66,67^, or the expression of AMX-hydrolising β-lactamases^68,69^. In gut microbiotas, β-lactamase producers may reduce AMX concentration and provide collective protection to sensitive species.

We reasoned that our *Enterobacteriaceae* counting on Drigalski agar should provide a first indication on whether wide spread resistance or collective protection was at play (Fig. S16). In robust donors (12/13/24/41/65), *Enterobacteriaceae* counts were not affected during AMX treatment despite the median proportion of resistant strains being moderate to low in Donor 13 (3.45 %), Donor 41 (9.6 %) and Donor 65 (50%). No AMX-resistant *Enterobacteriaceae* were counted in microbiotas from Donors 24 and 12. Therefore, sensitive *Enterobacteriaceae* persist during AMX treatment in AMX-Robust gut microbiotas. This supports the hypothesis of a collective protection.

In summary, our analysis suggests that gut microbiotas from AMX-Robust donors can collectively protect AMX-sensitive *Lachnospiraceae*, *Oscillospiraceae_88309* and *Enterobacteriaceae*. We proceeded to demonstrate that this collective protection was the result of β-lactamases.

### Amoxicillin collective protection is reduced by β-lactamase inhibition

If gut microbiotas from AMX-Robust donors are indeed collectively protected by β-lactamases (Fig. 4a), then inhibiting these enzymes shall impact protection. AMX is frequently employed in combination with clavulanic acid, an inhibitor of class A β-lactamases^70,71^. The resulting combination is called amoxicillin-clavulanic acid (amox-clav, AMC). We repeated our MBRA experiment, this time challenging the 16 gut microbiotas with 100 µg/mL of AMX and 20 µg/mL of clavulanic acid (Fig. 4b – see Material and methods). We also repeated our *Enterobacteriaceae* counting on Drigalski agar.

**Fig 4:**
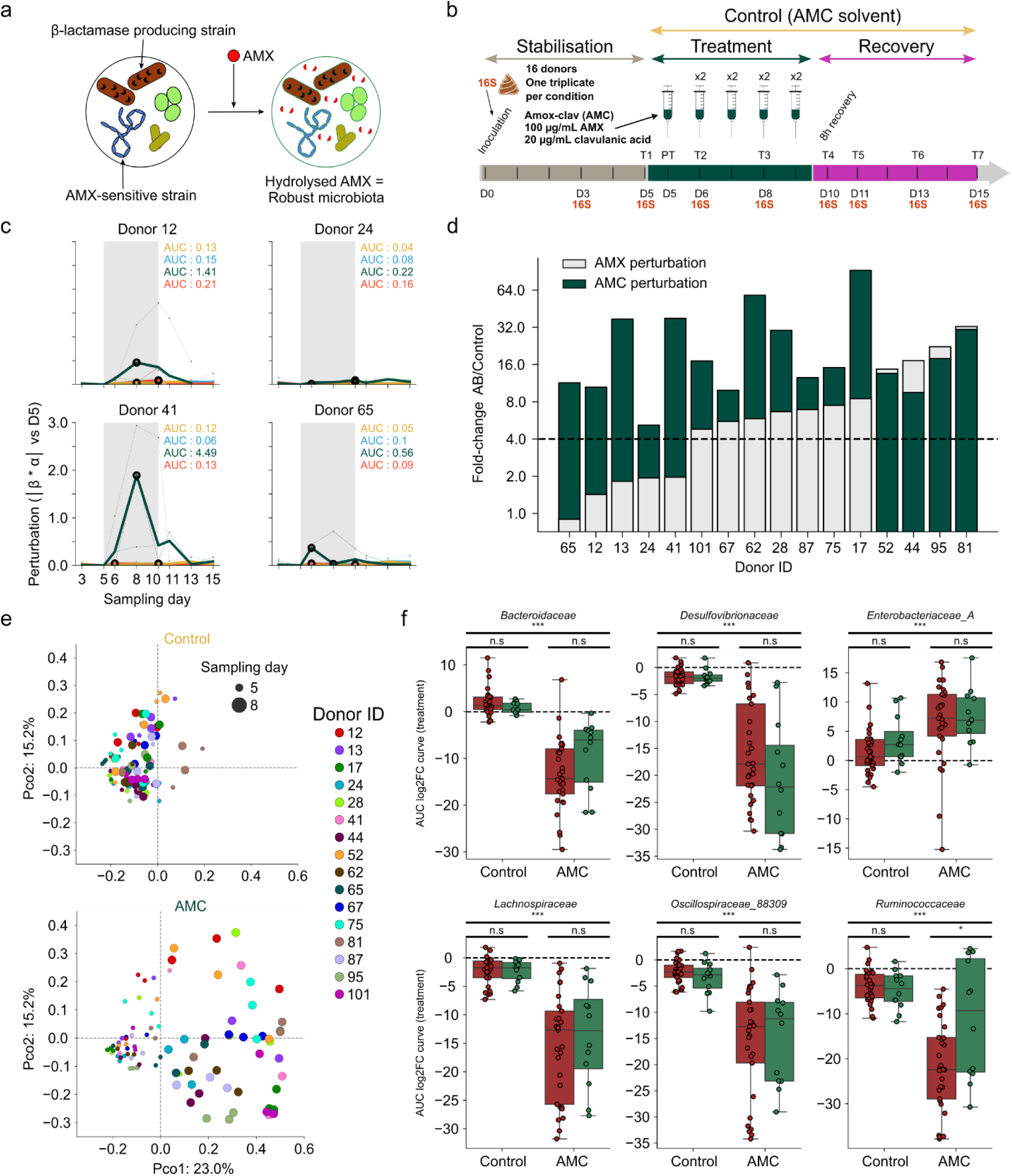
Addition of clavulanic acid greatly increases AMX perturbation even in AMX-Robust microbiotas. **a.** Proposed model tested to explain AMX robustness observed in microbiotas. Bacterial strains express β-lactamases that can protect sensitive families through hydrolysis of AMX. **b.** Representation of the experimental set-up using Mini BioReactor Arrays (MBRA) to test AMX-clav (AMC) perturbation on microbiotas. This experimental set-up is identical to the one displayed and discussed in Fig.1a. **c.** Perturbation curves of four AMX-robust donors with AMC. Red and blue curves show the results obtained with AMX and associated control condition (same data as in Fig. 1, shown for comparison). Dark green curves show chambers treated with AMC and ochre curves are treated with AMC solvent (control condition). The AUC of the median curve of each triplicate is also indicated. **d**. Bar plot representing the strength of AMX and AMC perturbations across all the 16 donors of the panel. The data for AMX is the same as in Fig.1f and used for comparison. **e.** Principal coordinate analysis of all bacterial communities of this study. Communities of all chambers at all sampling days were used to build a Jensen-Shannon divergence matrix. At the top are shown communities in control chambers at Day 5 and Day 8. At the bottom, communities treated with AMC are shown at Day 5 and Day 8. Dot size corresponds to the displayed sampling day. Dots are colored according to the Donor ID associated with the chamber. The plot is the same as Fig. 2b and PcoA coordinates were calculated with the entire dataset used in this article (AMX, AMX control, AMC, AMC control). **f.** AUC values of six major families in chambers either treated with AMC or untreated (Control condition). In green are chambers of AMX-Robust donors, and in red chambers of AMX-Sensitive donors. Mann-Whitney p-values are used to display significance (n.s= non-significant; * = p-value < 0.05; ** = p-value < 0.005; *** = p-value < 0.0005) when comparing the various groups (Robust vs Sensitive or Control vs AMC). Exact p-values and Cliff’s delta effect size are provided in the Data table (S21) Note that the tests and plots were done excluding donors 87 and 13.

Microbiota compositions across donors showed good consistency with the previous experiments, with some variability depending on taxonomic ranks (Supplementary note 5, Fig. S17-19). This validated possibilities of comparisons between both AMX and AMC experiments. Using both α-diversity and β-diversity metrics as previously, we observed that, except for Donor 24, all AMX-Robust donors were strongly affected by AMC (Fig 4c and Fig. S20). Using the perturbation curves, we found diverging perturbation patterns from the AMX experiment. With AMC, many microbiotas reached maximum perturbation at Days 8 or 10 instead of at Day 6. In some cases (e.g Donors 17, 41 and 101), we also observed long-lasting perturbations in AMC-treated microbiotas several days after treatment end.

Seeking to evaluate the impact of clavulanic acid, we compared the FCs calculated from the AMX and AMC experiments (Fig. 4d). Apart from Donor 24, that showed the lowest AMC perturbation of the panel (6.34), all other AMX-Robust donors experienced a strong perturbation in the presence of AMC, sometimes as high as the most AMX-perturbed donors (Donors 52/44/95/81). Additionally, many AMX-Sensitive donors were more perturbed by AMC (Donors 101/67/62/28/87/75/17). Using a PCoA to compare each chamber microbiota composition between Day 5 and Day 8 revealed how microbiotas were heavily impacted by AMC treatment (Fig. 4e).

Finally, to evaluate the impact of clavulanic acid on individual bacterial families, we computed their AUCs within each chamber as previously (Fig. 4f and Fig. S21). We focused on major families. *Bacteroidaceae* frequencies, which rarely decreased during AMX treatment, plummeted during AMC treatment (p-value<0.05, Cliff’s delta=0.948). *Lachnospiracaeae* and *Oscillospiraceae_88309* frequencies unsurprisingly decreased in most chambers (both p-values<0.05, respective Cliff’s delta=0.866 and 0.85), and both families were no longer stable in chambers of AMX-Robust donors (respective p-values=0.483 and 0.785, Cliff’s delta=0.144 and 0.057). Interestingly, both families frequencies tended to recover from Day 13, compared to Day 11 after AMX treatment (Fig. S22). As both families reflect well the intensity of the antibiotic selective pressure, this delay further exposes the strong intensity of the AMC treatment across donors.

The decrease of most families frequencies contrasted with increases of *Enterobacteriaceae* and *Enterococcaceae* frequencies in most chambers (Fig. S22). However, using our Drigalski agar counting data, we observed that frequency increases of *Enterobactericeae* sometimes hid a decrease in absolute counts within chambers (Fig. S23). For example, this was the case for Donors 67 and 81, and indicates a strong selective pressure from AMC even on *Enterobacteriaceae*. Furthermore, we observed the increase of AMX-resistant *Enterobacteriaceae* proportions in AMC-treated chambers from a subset of Donors (17/41/101). This observation highlights the AMC-mediated selection of resistant strains in the gut microbiotas of this donor subset.

In conclusion, the addition of clavulanic acid abolished most microbiotas robustness to AMX. This supports our β-lactamase collective protection hypothesis. Interestingly, there was still nuances between donors, and clavulanic acid increased perturbations even in donors that were already sensitive to AMX alone. This suggested that these microbiotas should still have a lower degree of collective protection. We reasoned that a complete understanding of AMX-response heterogeneity would require a dynamic and quantitative investigation of that collective protection.

### Dynamic donor-dependent depletion of AMX

To test whether donor’s microbiota display various kinetics of AMX degradation, we quantified AMX using mass spectrometry (Fig. 5a). Three donors were chosen from our panel: Donor 24 for its robustness to both AMX and AMC, Donor 41 for its robustness to AMX that is abolished by clavulanic acid addition, and Donor 17 for its sensitivity to AMX that is further enhanced by clavulanic acid. To inform us on the level of AMX loss irrespective of potential β-lactamase mediated degradation, we used a negative control (MBRA chambers with no microbiotas). A positive control involved the use of chambers inoculated with mono-cultures of a CTX-M (clavulanic acid sensitive) producing *E. coli* strain.

**Fig 5:**
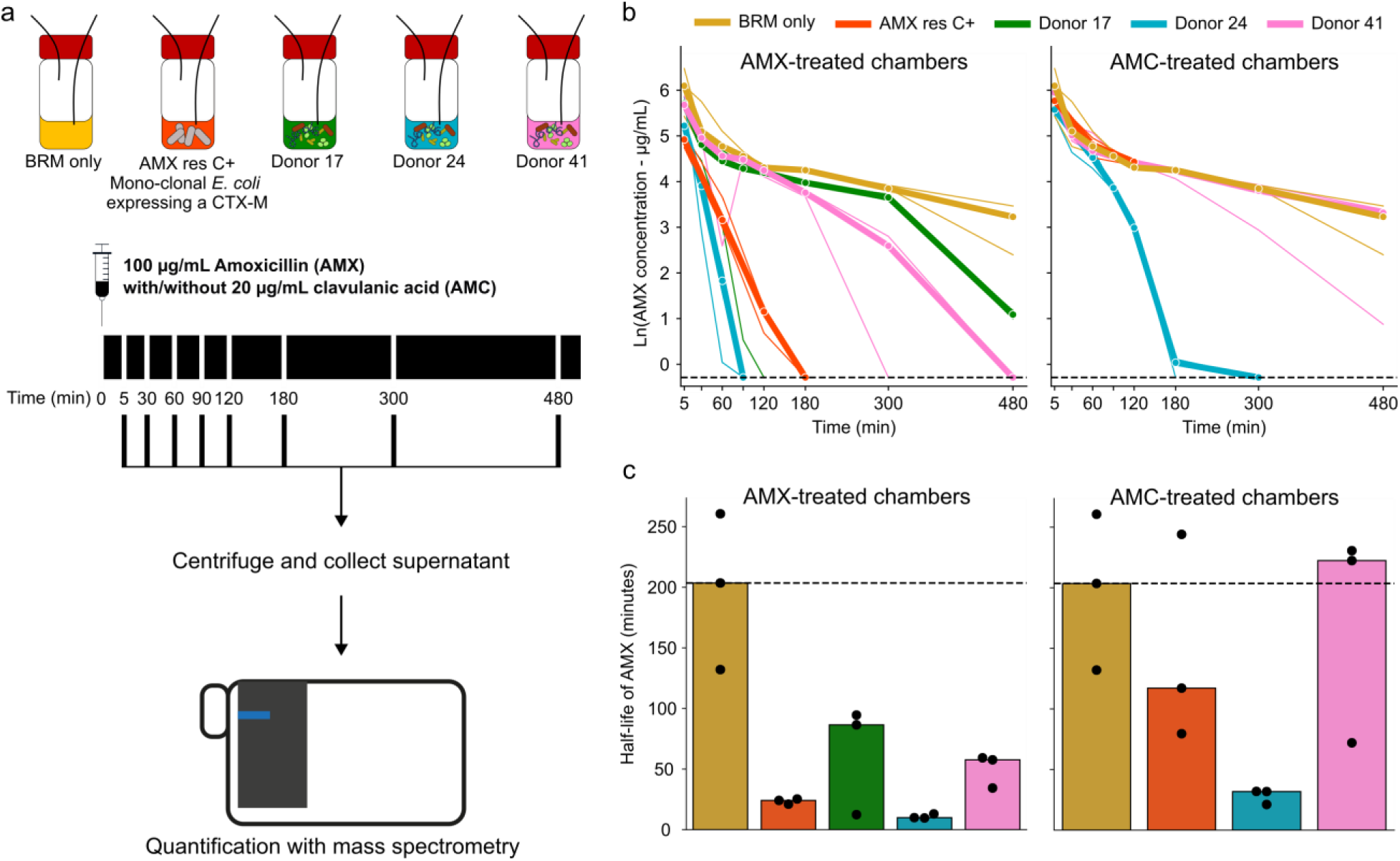
Microbiotas show heterogeneity in AMX depletion. **a.** Experimental set-up to quantify AMX in MBRA chambers. Microbiotas of Donors 17,24, and 41 were cultivated in MBRA chambers with the same method as previously described. After 5 days, AMX was added to the chambers to reach a final concentration of 100 µg/mL. Cultures were sampled from the chambers at the specified time points. Samples were then centrifuged and AMX was quantified in the supernatants with mass spectrometry (triple-quadrupole). A negative control (BRM only) and a positive control (Mono-culture of a CTX-M producing *E. coli*) were also used. All conditions were repeated three times. For Donors 41 and 24 and the positive control, three more chambers were inoculated with AMC. **b.** Evolution of AMX concentrations in each chamber in relation to time (minutes). Concentrations are given in Ln(µg/mL). Bold lines represent the median of the triplicates which are individually shown as thin lines. On the left are chambers treated with AMX and on the right are chambers treated with AMC. The BRM only control chambers, treated solely with AMX, are shown on the right for comparison purposes (same data on both graphs). The dashed line corresponds to the AMX detection limit (0.75 µg/mL). The curves of each replicate was then used to generate a linear regression, by default between the 30 minutes and the 480 minutes time points. Any data point with a concentration falling below the detection limit of 0.75 µg/mL, except the first one for each chamber, is removed from the linear regression calculation. Coefficients of the obtained regressions are then used to extrapolate an AMX decay constant, given in min-1. These constants are then used to extrapolate a half-life of AMX in each chamber. **c.** Plotting of the AMX half-life for each chamber, showing individual chambers as single dots and medians as full bars. The dashed line corresponds to the median of the BRM-only control chambers. On the left are AMX-treated chambers and on the right are AMC-treated chambers, with BRM-only chambers given for comparison purposes (same data as on the left). Statistical analysis of this data is given in Data Table (S28).

Microbiotas were cultivated as previously described. After stabilisation, triplicate chambers were inoculated with AMX to reach a final concentration of 100 µg/mL. For Donors 41 and 24, as well as for the positive control, AMC was supplemented in three more chambers to quantify AMX degradation in the presence of clavulanic acid. Samples were taken at multiple time points (5/30/60/90/120/180/300/480 min) during the 8 hours post-injection, and AMX was quantified in the supernatants after centrifugation (Fig 5b and Fig. S24). For each condition, an AMX decay constant λ (min^-1^) was calculated using the change of AMX concentration (µg/mL) over time (see Material and methods).

To compare donors, we then used the decay constants to estimate the half-life (HF) of AMX in each chamber, given in minutes (Fig. 5c). HFs varied greatly according to the various microbiotas and controls. First, we observed a median HF of 204 min in the BRM-only control. This can be explained by the MBRA dilution rate (1.875 ml/h) and a 10h HF of the antibiotic in BRM medium, likely due to non-enzymatic degradation (Supplementary note 6, Fig. S25). Oppositely, the median AMX HF in the positive control chambers was 8 times lower (25 min) and demonstrated that a monoculture of β-lactamase producer can rapidly affect the concentration of the antibiotic. A median AMX HF of 10 min was determined for microbiotas derived from Donor 24, indicating fast AMX depletion. Donors 17 and 41 had close AMX HFs (respective medians of 86 min and 58 min). Addition of clavulanic acid almost abolished AMX depletion in the positive control (median AMX HF of 118 min) as well as the one observed in Donor 41 which was now identical to the BRM-only control (median AMX HF of 222 min). However, Donor 24 kept a high depletion capacity (median AMX HF of 32 min). While all these results validate the idea of a protective effect in the robust microbiotas, one interesting observation from this data is that Donors 17 and 41 were not strongly different in their AMX depletion capacities despite important differences in the perturbation they endured when challenged by AMX.

One important outcome of the depletion may be the time a microbiota spends at an antibiotic concentration below the minimum inhibitory concentration (MIC) of some species. To compute this time in our setting, we simulated the dynamics of AMX concentrations in microbiotas of the three donors within a 24-hour treatment regimen as used in our experiment. We started with an 8-hour treatment period, followed by microbiota sampling and further addition of 100 µg/mL AMX and incubation for 16h (Fig. S26a). Models used the median AMX decay constant of each condition (Donor 17: 0.008 min^-1^; Donor 24: 0.07 min^-1;^ Donor 41: 0.012 min^-1;^ BRM only: 0.0034 min^-1^). As expected, the models predicted that microbiotas derived from Donor 24 spent limited amount of time at high antibiotic concentration (1266 minutes under 0.25 µg/mL). Furthermore, it revealed that mild differences in AMX decay constants may result in substantial gaps between time spent at low concentrations of antibiotics. For instance, within a 24-hour treatment regimen, Donor 41 microbiotas are expected to spend 252 more minutes than Donor 17 microbiotas under the AMX concentration threshold of 0.25 µg/mL, 271 more minutes under 4 µg/mL, and 212 more minutes under 8 µg/mL (Fig. S26b). In addition, the models predicts that even in the most perturbed microbiotas (e.g Donor 17), more than 800 minutes are spent bellow 8 µg/mL of AMX, especially within the 16h window. This may explain why *Enterobacteriaceae* populations were stable in our Drigalski agar counting data: sensitive *Enterobacteriaceae* may deplete in the first hours of treatment, but quickly recover when the antibiotic concentrations decrease bellow the MIC (Supplementary note 7 – Fig. S23). Overall, this data supports the hypothesis that microbiotas can deplete AMX in our MBRA chamber, most likely through the expression of β-lactamases. It further reveals the speed at which these changes in concentration can happen and how modest changes in depletion rate may change the amount of time microbiotas can recover in between AMX injections.

## Discussion

While the gut microbiota is highly exposed to antibiotic during treatment periods and presumably a major terrain for resistance selection, our understanding of antibiotic impact in this specific niche remains limited. Multiple studies have highlighted that gut microbiotas responses to antibiotics can be highly heterogeneous^6,21–23,26,72–74^, suggesting that the process is not as straightforward as expected. Multiple factors may contribute to such heterogeneity, many of which are host-related, including metabolic activity, transit rate, immunity, and diet^14,27,28^. In the present study, we focused on another potential source of heterogeneity: the microbiota itself. To this end, we used a controlled artificial *in vitro* gut system (MBRA) inoculated with human-derived stool to eliminate host-related confounding factors, while preserving complex gut microbiota composition^42,43^. Since most drugs are taken orally in human community settings, we used one of the most prescribed antibiotics, amoxicillin (AMX)^45^. We sampled faeces from 16 healthy individuals from the NutriNet-Santé cohort^46^, first stabilising the gut microbiotas in our system for 5 days. Afterwards, we administered two AMX pulses per day for five days, mimicking human exposure during antibiotic regimen. As we were able to cultivate distinct microbiotas with low variability between chambers from the same inoculum, the MBRA system allowed us to compare responses of multiple microbiotas to AMX treatment.

Combining a control condition with multiple replications, we consistently observed a gradient of AMX perturbation intensity across donors. This shows that even under tightly controlled conditions, the gut microbiota alone can shape the impact of antibiotic treatment. This heterogeneity of responses revealed several nuances. First, despite more than a 30-fold variation in perturbation intensity according to our metric, microbiotas could be grouped into two main categories. The AMX-Robust category (5 donors) comprised microbiotas showing almost no change in composition nor diversity during treatment. By contrast, the AMX-sensitive category was markedly heterogeneous, encompassing both modestly perturbed donors (101, 67) and highly perturbed ones (95, 81). These results indicate that microbiota composition alone drives much of the heterogeneity in treatment responses across models, and that the MBRA system can effectively identify the underlying determinants.

One such determinant may be the basal composition of the gut microbiota, i.e before treatment^30,31^. Indeed, each bacterial strain is characterised by its own minimal inhibitory concentration (MIC) towards a given antibiotic. Importantly, although we used AMX concentrations intended to exceed the MIC of most species (100 µg/mL), our mass spectrometry dynamic quantification of AMX showed that, within an 8-hour window, AMX concentrations decreased from 100 µg/mL to below 20 µg/mL in non-inoculated MBRA chambers, and dropped further in the presence of select microbiotas. Such declines may reduce the impact of AMX on many species during treatment, allowing partial recovery after initial exposure^75^. Day-to-day microbiota composition would thus result from a complex interplay involving differential resistance, lysis rate, growth and recovery speed—influenced by competitive release. Despite these dynamics, one may expect the overall intensity of perturbation to depend on the MIC distribution of constituent species at the time of exposure.

Indeed, members of the gut microbiota can display distinct ranges of AMX MICs. For example, sensitive strains of *Escherichia coli* typically have MICs between 2 and 8 µg/mL^a^. By contrast, members of the *Clostridia* class (such as *Lachnospiraceae* and *Oscillospiraceae_88309*) tend to have lower MICs (0.1–0.25 µg/mL), whereas members of the *Bacteroidaceae* family, such as *Bacteroides fragilis*, exhibit higher values (16–128 µg/mL)^a^. However, substantial variation exists even within families. *Salmonella enterica*, another member of *Enterobacteriaceae*, typically shows MICs of 0.5–1 µg/mL for most clinical isolates^a^, several-fold lower than those of *E. coli*. Additional examples from our dataset include marked heterogeneity within *Desulfovibrionaceae* and *Ruminococcaceae* families in microbiotas from AMX-Sensitive donors. Such diversity likely explains the absence of strong correlations between overall microbiota composition—or individual families—and perturbation intensity. This is well illustrated by Donors 13 and 95, which had similar microbiotas compositions but exhibited opposite perturbation patterns. Increasing taxonomic resolution to genus or species level may reduce within-family heterogeneity, but it also amplifies inter-donor differences and sequencing noise, creating further challenges for correlation analyses. Moreover, even within a given species, resistance can vary depending on the presence of specific resistance genes^76^.

Despite these limitations, we observed a strong and significant correlation between perturbation intensity and the *Lachnospiracea*e-to-*Bacteroidaceae* frequency ratio. This likely reflects both the high abundance of these families in most gut microbiotas^77^ and their relatively consistent AMX sensitivity (observed here and reported elsewhere^52,53,55^). As *Lachnospiraceae* are often described as highly sensitive to AMX, this supports the idea that strong initial killing can substantially disrupt the microbiota. However, this perturbation is not limited to the loss of a single family. For example, when perturbation intensity was recalculated after excluding *Lachnospiraceae*, it remained largely unchanged (Fig. S27).

This indicates that it is the loss of multiple species that reshapes the broader organisation of the microbiota. To explore this further, we used the MBRA system to investigate the dynamics of microbiota changes during treatment. Consistent with the observations above, we found a strong and significant correlation between perturbation intensity and the loss of two families: *Oscillospiraceae_88309* and *Lachnospiraceae*. Like *Lachnospiraceae*, *Oscillospiraceae_88309* is rarely reported to include AMX-resistant species^78^. The high abundance of *Lachnospiraceae* (20.5% on average before treatment) may partly explain its strong association with perturbation. However, *Oscillospiraceae_88309*, despite its lower abundance (7.4% on average), also showed a strong relationship. This suggests a potential structural role within the microbiota: its loss may trigger ecological imbalances that promote the expansion or decline of other families, such as *Enterobacteriaceae*, and contribute to dysbiosis. Consistently, we observed recovery of both *Lachnospiraceae* and *Oscillospiraceae_88309* after cessation of treatment, followed by restoration of community composition and diversity. This supports that members of *Oscillospiraceae_88309* could be explored as probiotics to support post-antibiotic recovery of the gut microbiota^79^. Further work will be needed to characterise this family in that context.

Using *Enterobacteriaceae* counts, we could rule out that the stability of *Lachnospiraceae* and *Oscillospiraceae_88309* in AMX-Robust microbiotas was due to ubiquitous resistance. Instead, it likely reflected a collective protection provided by other bacterial families in AMX-Robust microbiotas. This hypothesis was supported by supplementation of AMX treatment with clavulanic acid in MBRA chambers. The addition of this β-lactamase inhibitor, enzymes hydrolising AMX, markedly increased the impact of treatment. This indicates that β-lactamases expressed by some microbiota members provide a form of collective protection. This is consistent with models proposed in the literature, whereby β-lactamases hydrolyse AMX and reduce its effective concentration, thereby protecting more sensitive species^22,29,80^. However, the extent and dynamics of this protective effect have remained largely unexplored.

To investigate this further, we directly quantified AMX concentrations in gut microbiotas from three donors using mass spectrometry. We found that microbiotas differed markedly in their ability to deplete AMX, ranging from near-complete depletion within 1h30 (Donor 24) to minimal depletion (Donor 17). Clavulanic acid reduced this depletion capacity, indicating that it is partly mediated by clavulanic-acid-sensitive β-lactamases (e.g. Extended Spectrum β-lactamases - ESBLs)^70^. Two aspects of these experiments were notable. First, in Donor 24, depletion occurred at a remarkably rapid rate, even faster than in a pure culture of a β-lactamase-producing strain. However, our 16S data highlighted that this donor did not have a family exceeding 45% abundance in the 16S data. This suggests that strong and rapid protective effects can be either exerted by a single bacterial family not in absolute majority, or combinations of families. Second, the difference in depletion between Donors 17 and 41 was modest, despite their classification as AMX-Sensitive and AMX-Robust, respectively. Modelling AMX concentrations based on measured half-lives showed that even small differences in depletion capacity can substantially alter the time spent below critical concentration thresholds. For instance, microbiotas from Donor 41 spent more than seven hours per day below 0.25 µg/mL whereas those from donor 17 spent 3-hours and a half below that threshold. These results indicate that collective protection, through its dynamic modulation of antibiotic concentration, is a key determinant of both the intensity and variability of microbiota perturbation.

To our knowledge, this study is among the first to propose such a high-resolution dynamic view of antibiotic concentration and microbiota perturbation. A notable previous study explored such dynamics with three β-lactams and 121 human patients, using modelling to compensate for reduced resolution as available here with our MBRA system^81^. Future work could extend this approach to other antibiotic classes beyond β-lactams. For example, a study in morifloxacin-treated human patients combined plasma and faeces antibiotic dosage with microbiotas diversity monitoring, showing substantial dynamics and heterogeneities between patients^82^. In particular, it would be informative to determine whether collective protection is specific to antibiotics than can be enzymatically degraded, or whether similar effects arise for antibiotics subjected to modifications (e.g aminoglycosides, macrolides and tetracyclines) or with no known or widespread degradation pathways (quinolones)^83^. Such studies may reveal previously unrecognised mechanisms of collective protection within microbiotas. We advocate that the MBRA system, because it provides a suitable framework for high-resolution dynamics, is ideal to complement other modelling approaches in human patients.

A further objective of this study was to examine the dynamics of antibiotic resistance selection across the 16 donor microbiotas. Coherently with the aforementioned results, we observed heterogeneous selection of AMX resistance within *Enterobacteriaceae* populations in MBRA chambers. As expected, no clear selection of resistant *Enterobacteriaceae* occurred in AMX-robust donors. By contrast, in AMX-sensitive microbiotas, we frequently observed the emergence of resistant *Enterobacteriaceae*, often accompanied by stable or increasing total counts. These findings indicate that collective protection can modulate, and in some cases limit, the selection of resistance.

Interestingly, resistance selection was less frequent in chambers treated with amoxicillin–clavulanic acid (AMC). The addition of clavulanic acid led to a marked decline in total *Enterobacteriaceae* counts, followed by recovery after treatment cessation. This suggests that most *Enterobacteriaceae* selected under AMX treatment express β-lactamases, consistent with the reduced selection observed in the presence of clavulanic acid. Once established, such strains may confer protection to the broader community, reducing perturbation and accelerating recovery. This may explain that peak perturbation occurred around day 6 in 7 out of 11 AMX-sensitive donors. Furthermore, we observed a decline of AMX-resistant *Enterobacteriaceae* after AMX removal, which hints those resistant strains have a fitness defect in recovered microbiotas. By contrast, AMC treatment produced more sustained perturbation and slower recovery accompanied by maintenance of resistant populations, which is in line with this model. We also noted other interesting patterns (Supplementary note 8). These results underline that resistance selection is shaped by collective protection and follows highly dynamic trajectories.

Collectively, our findings indicate that the equilibrium between sensitive and resistant bacterial families (*Lachnospiraceae*, *Oscillospiraceae* and *Bacteroidaceae*) as well as collective protection plays a central role in shaping gut microbiota responses to AMX treatment, including both dysbiosis and resistance selection. This effect is mediated through dynamic changes in microbiota composition as well as in antibiotic concentration experienced by the microbiota. These observations have several implications. Our mass spectrometry measurements indicate that relatively small shifts in antibiotic concentration can be sufficient to influence dysbiosis and resistance selection. This supports the use of probiotics to provide collective protection (β-lactamases producers), which could be naturally producing gut microbiotas members (e.g *Bacteroidaceae*) rather than engineered highly-efficient strains of which applications often face regulatory barriers^84^. Another potential clinical application of our findings could be to evaluate the degree of collective protection within a patient gut microbiota before choosing an antibiotic therapy. This would guide personalised antibiotic choices—for instance, favouring β-lactams that are readily degraded by the patient’s microbiota. This could be evaluated using β-lactamase activity assays on faecal samples (as with the nitrocefin assay^85^), and may be particularly relevant for immunocompromised patients for which enteric pathogen colonisation (e.g. *Clostridioides difficile* infection^86^), and the spread of antibiotic resistance are priority problems^87,88^.

## Material and methods

### Selection of 16 donors from the NutriNet-Santé cohort

Fecal samples were obtained from 103 donors enrolled in the NutriNet-Santé cohort with previously documented dietary fiber intake. Detailed cohort description, dietary assessment, microbiological analyses, and associated results will be reported elsewhere. Donors were stratified into three dietary fiber intake categories: high fiber (Fib+), low fiber (Fib−), and intermediate or heterogeneous intake (Random). For each fecal sample, a single *Escherichia coli* isolate was recovered and assigned to a phylogenetic group. In parallel, the pooled aerobic Gram-negative population growing on Drigalski agar was subcultured on agar supplemented with 8 mg/L amoxicillin in order to assess resistance structure. Five qualitative ecological configurations were defined according to the relative abundance and amoxicillin susceptibility of Gram-negative bacteria and *E. coli* populations. One configuration, characterised by a dominant susceptible *E. coli* population coexisting with a subdominant resistant *E. coli*, was considered particularly informative for studying selective-pressure dynamics. A stratified random subset of 16 donors was generated to preserve fiber-intake structure while ensuring representation of phylogenetic diversity and resistance configurations. The final subset included 5 Fib+, 5 Fib−, and 6 Random donors and was further divided into two balanced sets of eight samples for downstream experimental analyses. Samples from smokers with atypical microbiota profiles (n = 2) and samples lacking detectable *E. coli* (n = 4) were excluded prior to sampling because the study aimed to investigate treatment-driven dynamics within *E. coli* populations. Three independent random samplings were evaluated. The first two were discarded due to over-representation of phylogroup D strains. The third sampling maintained balanced dietary distribution, phylogenetic diversity, and resistance configurations and was therefore retained for subsequent analyses. In graphical representations, selected donors are indicated by color-coded markers corresponding to fiber-intake categories: red for Fib+, green for Fib−, and blue for Random.

Among the 103 fecal samples with documented dietary fiber intake, donors were distributed across the three predefined fiber-consumption groups. Microbiological characterisation identified recoverable *Escherichia coli* isolates in most samples and revealed heterogeneous resistance structures within aerobic Gram-negative communities grown on Drigalski agar supplemented with amoxicillin. Qualitative classification defined multiple ecological configurations reflecting variation in dominance patterns and amoxicillin susceptibility among Gram-negative bacteria and *E. coli* populations. In particular, the coexistence of dominant susceptible and subdominant resistant *E. coli* populations highlighted potential gradients of selective pressure within the gut ecosystem. obtain an experimentally tractable yet representative subset, 16 donors were selected using stratified random sampling that preserved fiber-intake distribution while capturing phylogenetic and resistance diversity. The resulting subset comprised 5 Fib+, 5 Fib−, and 6 Random donors and was organised into two balanced groups of eight samples. Exclusion of smokers with atypical microbiota profiles and samples lacking detectable *E. coli* ensured consistency with the study objective of analyzing treatment-associated dynamics within *E. coli* populations. Among three independent random samplings, the first two displayed phylogroup D over-representation and were therefore excluded. The third sampling provided balanced representation of dietary categories, phylogenetic diversity, and resistance configurations and was retained for all downstream analyses. Graphical visualization confirmed the balanced structure of the final subset, with selected donors represented by color-coded markers corresponding to fiber-intake groups.

### Mini BioReactor Array design and cultivation

MBRA systems were used as described previously^42,89^. Briefly, a MBRA system is composed of 24 independent chambers placed on a magnetic stand for continuous homogenisation in an anaerobic chamber. The 24 chambers are connected to 2 peristaltic pumps allowing a continuous flow in the chamber (530S drive and a 205Ca8 8 channel roller plus a 505CAX16 16 channel extension, Watson and Marlow; IPC 24 channel pump, Ismatec). After preparation, the system was sterilised and placed in the anaerobic chamber for at least 72 hours. The 24 chambers were then filled with 15 mL of BioReactor Medium^42^. The medium stayed for 5 hours in the chambers without flow or homogenization. In the meantime, the feces were prepared using a solution with 2.5% of faeces homogenised with Dulbecco‘s Phosphate Buffered Saline (14190-144, Gibco) which was vortexed during 5 min, then centrifuged at 800 rpm for 5 min. Supernatants were filtered on a 100 µm-cell strainer (352360, Falcon) to avoid chunks. Finally, 3.8 mL of this preparation was inoculated in each chamber. The experiment was designed so that one donor had two conditions (treated and untreated), with each condition in triplicate. After 16 hours, the pumps were started with a 1.875 mL/hour flow rate, which correspond to an 8-hour turnover of medium, mimicking the intestinal flow. After 5 days of stabilisation of the microbiota to the culture conditions, the corresponding chambers were treated with 100 µg/mL amoxicillin (with or without 20 µg/mL clavulanic acid) twice a day (one dose in the morning and one 8 hours later). For the control chambers, the same volume (150 µL) of water (AMX solvent) was injected instead. The amoxicillin (CIP: 3 400 956 035 885, Panpharma) was solubilised with sterile water and the amoxicillin-clavulanic acid (19456, Cayman Chemical) mix was prepared with Phosphate Buffered Saline (10010-015, Gibco) with a 5:1 ratio. After 5 days of treatment, there was a 5-days recovery phase. 400 µL-samples were collected nearly every two days during each phase (Fig 1a). The samples were stored at −80°C and separated in two different tubes. One tube was used for the 16S sequencing (200 µL), while the other 200 µL were mixed with 25% glycerol for storing in preparation of the antibiotics screening experiment.

### Antibiotics Screening on Drigalski

Drigalski agar (CM0531B, Thermo Scientific Oxoid) plates were prepared with and without antibiotics. The antibiotics used for this screening were amoxicillin (CIP: 3 400 956 035 885, Panpharma) at 32 mg/L and amoxicillin and clavulanic acid at a 5:1 ratio (19456, Cayman Chemical) at 32 mg/L. Samples at different time-points were plated: before the first treatment (D5), 1 hour post-treatment (PT), 24h after the first treatment (D6), the fourth day of treatment (D8), 8h after the last treatment (D10) and 24h after the last treatment (D11). The samples were diluted in 0.01M Magnesium sulfate heptahydrate (63138, Sigma) and plated by spotting different dilutions, from pure to 10^−5^.

### Amoxicillin dosage using mass spectrometry

Amoxicillin concentrations were determined by LC-MS/MS using a procedure previously described^90^. Amoxicillin-13C6 was used as the internal standard. For consistency with the calibration standards prepared in human plasma, all supernatants were first diluted 1:10 in blank human plasma. Briefly, sample preparation involved protein precipitation using acetonitrile followed by dilution of the supernatant. The analytical system consisted of either an UltiMate™ 3000 HPLC chain coupled to a TSQ Vantage™ mass spectrometer or a Vanquish™ HPLC system coupled to a TSQ Altis Plus™ triple quadrupole mass spectrometer (Thermo Fisher Scientific, Waltham, MA, USA). Chromatographic separation was achieved on an Acquity HSS T3 (2.1 x 50 mm, 1.8 µm) column (Waters Corp., Milford, MA, USA) using a gradient of water and acetonitrile, both acidified with 0.1% formic acid. Compounds were ionised using an electrospray source in positive mode. The calibration range extended from 0.075 µg/mL to 75 µg/mL, resulting in a lower limit of quantification (LLOQ) of 0.75 µg/mL after accounting for the dilution factor.

### Administration frequency and concentration of amoxicillin (with or without clavulanic acid) in MBRA experiments

To our knowledge, very few studies have reported fecal concentrations of amoxicillin following oral administration. One publication analysed feces from individuals who had received antibiotic treatments, including amoxicillin, in order to assess the impact on waste composting^91^. In that study, one of the tested concentrations was 100 µg per gram of dry faeces. By extrapolating grams of faeces to milliliters of MBRA medium, this concentration was considered appropriate for use in the present experiments. The administration schedule consisted of two injections per day to approximate standard clinical regimens of amoxicillin (with or without clavulanic acid), such as those used for pneumonia or upper respiratory tract infections, in which oral amoxicillin is typically prescribed two to three times daily in community settings^45^.

### Bacterial DNA Extraction

Genomic DNA was extracted from frozen MBRA suspensions using the DNeasy® 96 PowerSoil Pro QIAcube HT Kit (Qiagen, Hilden, Germany), preceded by an initial mechanical disruption step as previously described^41^. Briefly, for mechanical lysis, 100 µL of each sample was added to a PowerBead Pro Plate containing and mixed with 800 µL of CD1 lysis buffer. Samples were disrupted using a TissueLyser II (Qiagen) for 2 minutes at 25 Hz. The plates were then centrifuged at 2773g for 10 min, and 500 µL of the supernatant were collected and transferred to a new 96-well plate. Then, 250 µL of CD2 was added and mixed followed by centrifugation at 2773g for 10 min. A total of 550 µL of supernatant was transferred to a new plate. Purification with silica membrane plates and elution steps were performed using the QIAcube® HT automated platform (Qiagen) according to the manufacturer’s instructions.

### Bacterial Community Analysis by 16S rRNA Gene Sequencing

The taxonomic composition of bacterial communities was revealed by 16S rRNA gene amplicon sequencing using the Illumina MiSeq platform as described earlier^41^. The <u>V4 region</u> of the 16S rRNA gene was amplified by PCR using a forward primer 515F (5’-AATGATACGGCGACCAC-CGAGATCTACACGCTXXXXXXXXXXXXTATGGTAATTGT<u>GTGYCAGCMGCCGCGGTAA</u>-3’) and a reverse primer (806R 5’CAAGCAGAAGACGGCATACGAGATAGTCAGCCAGCC<u>G-GACTACNVGGGTWTCTAAT</u>-3’) comprising unique 12-base barcodes to enable sample-specific identification of amplicons^92,93^. Amplicons were verified by agarose gel electrophoresis and quantified using Quant-iT Picogreen dsDNA assay (Life Technologies) prior to pooling in equimolar concentrations. The resulting library was purified using AMPure magnetic beads (Agencourt, Beckman Coulter) and sequenced (Paired-end reads, 2×250bp, V3 reagents) on an Illumina MiSeq platform at the Genom’IC facility (Institut Cochin, Paris, France).

### Pre-Processing of 16S rRNA gene sequence

The 16S rRNA sequences were analysed using QIIME2 (version 2024)^47^. As an initial step, paired-end sequence data were demultiplexed using the demux plugin by assigning barcode reads to sample identifiers, following the Earth Microbiome Project (EMP)^94^ amplicon sequencing protocol. DADA2^95^ was employed for denoising and quality-based trimming of the reads, with the parameter --p-trunc-len-r set to 180 to truncate reverse reads at positions where the Phred quality score dropped below 30, enabling the identification and correction of Illumina amplicon sequencing errors and the generation of Amplicon sequence variants (ASV) table. Taxonomy was assigned to all amplicon sequence variants using a 99% pairwise identity threshold with the Greengenes2 reference database (2024.09) via the q2-feature-classifier plugin.

### Data analysis and figure construction

Taxonomic tables obtained from the sequencing and data processing pipeline aforementioned were processed using a custom Python (V 3.12.4) code. Multiple packages were used for the processing.

| Module names and versions | Function |
| --- | --- |
| Pandas (V2.2.2)<br>numpy (V1.26.4)<br>Microsoft Excel (V2108) | Data tables handling. |
| scikit-learn (V1.4.2)<br>scikit-bio (V0.6.2)<br>scipy (V1.15.3) | Statistical tests, diversity metrics and principal co-ordinates analysis. |
| seaborn (V0.13.2)<br>matplotlib (V3.8.4) | Figure making. |
| Inkscape (V1.4)<br>Microsoft Powerpoint (V2108) | Figure processing and scheme making. |

Statistical tests used for the data, with their inputs and associated Python modules are provided in the Data table (S32). Significance threshold for p-values was of 0.05. This threshold was the same for false discovery rate after using the Benjamini-Hochberg correction. All p-values and effect sizes were computed with tests that did not assume normality (Spearman correlation, Mann-Whitney U, Cliff’s delta, PERMANOVA, PERMDISP). This was decided due to the high amount of noise from our 16S amplicon sequencing dataset, and the inherent variability from the MBRA system, with one replicate often strongly deviating from the two others. We applied the same logic to the AMX quantification dataset. In the latter case, the statistical tests are not discussed in the text, as we evaluated the sample size to be too restrictive. The results of our tests can be found in the Data table provided with this paper (S28).

For read counts data analysis, each Illumina sample with less than 3500 total reads was discarded from the dataset. Furthermore, each OTU which appeared in less than 10% of all samples was discarded. However, OTU frequencies were calculated prior to this curation. We decided for this approach to reduce noise generated by rare OTUs while keeping frequencies intact for diversity metrics. For the plotting of frequencies for each donor (Fig S2 and S18), OTUs which appeared in less than 3 samples were not shown to improve the readability of the figures. The same logic was applied for plotting the frequencies changes of each family, with the removal of some OTUs which were only appearing occasionally. The data corresponding to all these OTUs can be found in the Data Table, irrespective of this curing (S20).

We frequently encountered samples with 0 reads which made impossible calculation of log2(FC) of bacterial families vs Day 5. This happened often with *Lachnospiraceae*, as this OTU was often disappearing in AMX-treated chambers during treatment. To remediate this, we considered OTU to represent an equivalent of 1 read for samples involved. If log2(FC) of two samples with 0 read was calculated, its value was directly given as 0. All 0 were then ignored for the subsequent analysis.

When using PERMANOVA and PERMDISP, we compared the Robust vs Sensitive groups. Since both groups did not have the same size, we randomly sampled from the Sensitive group to match the size of the Robust group.

Read rarefaction was done using the *skbio.stats.subsample_counts* function from scikit-bio, setting the depth to 3500 reads and seed to 42.

### Calculation of AMX decay constants and half-lives

For the calculation of AMX decay constants, AMX concentrations obtained from mass spectrometry quantification were plotted on a natural logarithm scale (Fig 5b). Resulting curves were used to calculate a regression coefficient. We noticed that the concentration of AMX did not reach 100 µg/mL in most chambers before 60-90 minutes, and reached high concentrations in the first time points. This was likely caused by a slow homogeneisation of AMX levels in the MBRA chambers due to agitation, or by presence of AMX stock solution in the sampling needle at first time points. For this reason, coefficients were calculated depending on manually selected time-windows to primarily restrict noise from the first data points of the curves (See Data table S27). This coeffcient was the decay constant λ, given in min^-1^. Following an exponential decay equation, we could then find the half-lives (t ½) of AMX in each chamber according to λ, as:

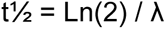

With t½ given in minutes. The same equation was used to determine the time at which AMX concentrations reached 8/4/0.25 µg/mL, respectively with Ln(12.5)/Ln(25)/Ln(400), considering a staring concentration of 100 µg/mL.

The median λ values for each triplicate chambers were used to build the model presented in Fig. S26. These values were also used to determine the time spent under each AMX concentration thresholds (8/4/0.25 µg/mL), taking into account an 8h-16h treatment regimen and that AMX concentration increases by 100 µg/mL after 8 hours of treatment.

### Exclusion of Donors 87 and 13

We opted to exclude Donors 87 and 13 from the data obtained in the AMC experiment. The two donors and their associated data points from the AMC experiment were excluded from analyses done for Fig. 4f, S21 and S22: data points corresponding to these donors’ chambers were not used to calculate the Medians, Mann-Whitney p-values and Cliff’s delta effect sizes. Data corresponding to these two donors can still be found in the data tables provided with this paper.

## Data and code availability

Python code will be made available on Github upon publication. A table containing all the data used to make the figures will be provided upon publication as a supplementary file.

## Acknowledgments

We thank Charles Burdet for discussions on data analysis. We thank Chloé Robert and Jeanne Volckaert for their help with library preparation and communities sampling. We thank the Genom’IC platform of the Cochin Institute for carrying the amplicon sequencing.

## Authors contribution

**Paul Lubrano** : Data curation, Formal analysis, Investigation, Methodology, Visualization, Writing – original draft.

**Mélanie Magnan** : Investigation, Methodology;

**Caroline Steiner** : Investigation, Methodology;

**André Birgy** : Conceptualisation, Investigation, Methodology;

**Benoit Chassaing** : Conceptualisation, Investigation, Methodology, Supervision;

**Mélanie Deschasaux-Tanguy** : Investigation, Methodology;

**Arnaud Guttierez** : Investigation, Methodology;

**Hugo C. Barreto** : Investigation, Methodology;

**Sophie Magréault** : Investigation, Methodology;

**Vincent Jullien** : Investigation, Methodology;

**Mathilde Lescat** : Conceptualisation, Investigation, Methodology, Supervision;

**Olivier Tenaillon** : Conceptualization, Funding acquisition, Investigation, Methodology, Project administration, Resources, Supervision, Validation, Writing – original draft.

## Supplementary notes

### Supplementary note 1

We first investigated α-diversity metrics, which capture the diversity of families within a sample. We focused on Shannon entropy that includes families’ frequencies in its computation of diversity. Changes in each chamber during the MBRA cultivation were evaluated with log2 fold-changes against the Day 5 α-diversity. A curve was then constructed with the resulting values, representing the changes of α-diversity across the entire experiment. We then calculated area under curve (AUC), before, over, and after the days of treatments as proxies for perturbation. As expected, this analysis revealed an overall reduction of diversity under treatment (Fig. 1c, Fig S3-4). Median of all AUCs for chambers in control condition reached −0.12, while a value of −0.94 was observed in AMX-treated chambers, with significant differences between all AUCs of both groups (p-value<0.05, Cliff’s delta=0.67). These changes were connected to treatment, as there were no significant differences between AUCs of both control and AMX groups before treatment (p-value=0.19, Cliff’s delta=0.15). We observed a recovery of initial diversity at end of the experiment, as shown by a reduction in differences between AUCs of both control and AMX groups (p-value=0.05, Cliff’s delta=0.23). Perturbation of the diversity was however heterogenous amongst the donors. For example, Donors 24 and 41 showed almost no variation during treatment compared to the control condition (respective Control/AMX medians of −0.65/-0.53 and −0.25/-0.46).

To better characterise the changes that happened in the community, we resorted to β-diversity metrics (Jensen-Shannon divergence (JS) or Bray-Curtis dissimilarity (BC) – Fig. 1d, Fig. S5-6) that measure composition shifts between communities. We used the compositions of chambers before treatment (Day 5) as reference. In the control conditions, we observed a gradual increase of β-diversity until day 15, as communities drifted away in their composition. The intensity of that process was variable across the populations. For example, control conditions of Donor 101 only slightly varied (median JS of 0.18 on Day 15) while Donor 12 controls strongly diverged from their Day 5 compositions (median JS of 0.33 on Day 15). We also observed noise in the system, with occasional replicates completely diverging from the two others. This was the case for Donor 17, of which one control replicate (Rep B) had a notable frequency increase of *Clostridiaceae_222000* from Day 6. Taken together, these two observations indicated that some communities were more volatile than others, and that our system, while robust, could also be prone to random variations. This noise validated the use of a control condition as a reference and the use of triplicates to not over-evaluate AMX perturbation.

The β-diversity metrics revealed that the AMX treatments induced overall large changes in community compositions. Median AUC of all JS curves from control chambers reached 0.87, and 1.47 from all AMX-treated chambers (p-value<0.05, Cliff’s delta=−0.7), with no significant differences between both control and AMX groups before treatment (p-value=0.53, Cliff’s delta=0.07). However, as observed for the α-diversity, the treatment response was heterogeneous between donors. For example, Donors 81 and 95 showed a strong response (respective JS Control/AMX AUC medians of 0.6/2.18 and 0.8/2.03), while Donors 12 and 24 showed almost no more variations than their controls (respective JS Control/AMX AUC medians of 1.1/1.32 and 0.7/0.85).

### Supplementary note 2

To test the sensitivity of this analysis to the choice of phylogenic level, we repeated our analysis at the class, order, and genus levels (Fig. S8). We observed a subset of donors which alternated between the robust and sensitive groups (62,67,75,81,101), likely because of strong intra-taxa heterogeneities. However, the AMX-robust donors we previously categorised (12/13/24/41/65) were not reclassified as sensitive in any taxonomic level chosen. This demonstrates that they face low-level AMX-driven perturbations to their diversities no matter the phylogenic group chosen for our analysis. Furthermore, we repeated this analytical pipeline using read rarefaction as a normalisation method, focusing on the family taxonomic level and using JS and Shannon entropy as metrics. Although FCs slightly varied between donors, donor classification was unchanged, further demonstrating the robustness of our analysis (Fig. S9).

### Supplementary note 3

Comparing the sensitive and robust groups, PERMANOVA (Pseudo-F=2.44; p-value=0.0048) was significant, but so was PERMDIST (F-value=12.17; p-value=0.0018), indicating a significant dispersion within either group (Fig. 2a). Focusing on single chambers, no correlation was found between Shannon entropy at Day 5 and perturbation, given as AUC of the perturbation curve (Control/AMX Spearman ρ= −0.07/0.131 and p-value=0.63/0.38 –Fig. S10). Additionally, comparing each chamber β-diversity in a pair-wise fashion (Fig. S11), we observed no strong correlation between pair-wise Jensen-Shannon divergence at Day 5 and Day 6 in AMX-treated chambers (Spearman ρ=0.295, p-value<0.05). That implies that, one day after the start of AMX treatment, chambers that have low community divergence at Day 5 can strongly shift from each other at day 6, confirming that composition is not predictive of perturbation. On the other hand, a strong correlation was found in Control chambers (Spearman ρ =0.783, p-value<0.05), indicating that, without AMX, communities shifts were low and divergence between donors and replicates were maintained from Day 5 to Day 6.

### Supplementary note 4

Family’s abundance changes at a sampling-day resolution showed interesting patterns (Fig. S14). For example, abundances of *Lachnospiraceae* in Robust vs Sensitive communities were no longer significantly different from Day 13 (p-value=0.276, Cliff’s delta=−0.2). This supports the previous observations that bacterial communities can generally return to their pre-treatment states^37,52,96^. Since *Lachnospiraceae* and *Oscillospiraceae_88309* are known to contain spore producers^97,98^, it is possible that they adopt such strategy during AMX treatment. This would explain their observable decrease in frequency followed by recovery after treatment ends, the DNA of *Firmicutes* spores being complex to extract without specialised kits^99^.

### Supplementary note 5

We first analysed the community compositions of our chambers and compared them to the ones of our previous experiment (Fig. S17 and Fig. S18). Half of the donors of the panel (81/101/12/44/17/95) had their community clustering remarkably well with the ones of the previous experiment. However, disparate clusters were noticed for the rest of the donors. Hence, we repeated the analysis at the genus level, expecting to highlight better the disparities between the replicate experiments (Fig. S19). At this level, replicates clustered better between donors. This highlights that the two independent MBRA experiments conserved most donors specificities although being subjected to noise regarding family frequencies. As we previously highlighted that the community phenotype was likely more relevant than frequency distributions within communities, we judged that this was not an issue for pursuing our analysis.

However, there were some inconsistencies that lead to the exclusion of two donors: Donor 87, in which large divergence between replicates was observed at Day 5, and Donor 13, largely dominated by *Enterococcus_B*, which we did not observe in the AMX replicate. Therefore, we opted to exclude the two donors from analyses involving comparison with the AMX experiments (see Material and methods for more details).

### Supplementary note 6

We observed that, in BRM-only chambers, AMX median half-life was of 203 minutes. We hypothesised that this depletion was the result of the medium renewal in the MBRA chambers (1.875 mL/h – 31.25 µL/min) associated with the degradation of AMX in BRM medium.

We first calculated the half-life of AMX, only taking into account the dilution factor of 31.25 µL/min and a starting concentration of 100 µg/mL. We obtained the value of 332 minutes. Using this value, we then determined a decay constant (See material and methods).

AMX half-life was previously measured to be 27.4h in Mueller-Hinton broth^100^. We used this as a reference for the stability of AMX in BRM medium. Then, irrespective of medium renewal, we determined a new decay constant with this half-life value. Two more theoretical half-lives of 5h and 10h were then added as additional references. This gave us three decay constants, to which we added the medium renewal decay constant previously calculated.

We finally modeled the loss of AMX by taking into account these decay constants and the BRM-only decay constant we experimentally determined, and compared the resulting curves (Fig. S25). The BRM-only curve was close to the model involving both medium renewal and an AMX half-life of 10h. Hence, we conclude that AMX likely has a 10h half-life in our MBRA system. Overall, this data predicts that, even with no communities, AMX levels can reach levels lower than 8 µg/mL after 16h of incubation in our MBRA system (Fig. S26a).

### Supplementary note 7

These short decay and fast recoveries may somehow limit the selection for antibiotic resistance, a process that seemed more marked when the perturbation was stronger with AMC (Fig. S23). Indeed, in many communities treated by AMC, *Enterobacteriaceae* total counts fell below our detection limit (10^4^) which was not the case with AMX treatment. In both Donors 17 and 41, there was an increase of AMX-resistant *Enterobacteriaceae* proportion during the treatment, reaching almost 100%. Donor 41 median AMX HF under AMC treatment being almost identical to our BRM-only condition (222 min), communities of that donor would spend less than 3 hours under the 8 µg/mL threshold, likely not enough for its *Enterobacteriaceae* populations to recover from the treatment before a new injection. This should either lead to population loss and/or strong resistance selection.

### Supplementary note 8

An interesting case is the AMX-Sensitive Donor 95 in which no AMX-resistant *Enterobacteriaceae* was detected despite having a stable *Enterobacteriaceae* population during both AMX and AMC treatments. AMX and AMC perturbations were almost identical with this Donor, suggesting no trace of collective protection and fluctuations of AMX levels similar to those in sterile MBRA chambers. That no resistance was selected and *Enterobacteriaceae* populations maintained opposes Donors 41 and 17 observations (see above). In this Donor 95 microbiotas, fluctuations in AMX levels may favor rapidly growing *Enterobacteriaceae* (absent from Donor 41) with higher MICs which quickly recover without necessarily being resistant to high AMX levels. Other interesting patterns include Donors 75 and 101 which had almost fully resistant *Enterobacteriaceae* populations respectively before AMC and AMX treatments, and were both perturbed. In these cases, it is possible that resistance was caused by other mechanisms than β-lactamases, or that the *Enterobacteriaceae* population was not capable of providing collective protection for multiple reasons (expression level, catalytic rate of the expressed β-lactamase).

## Footnotes

a The data/graph presented is from the EUCAST MIC and Zone diameter distribution website (https://mic.eucast.org), last accessed the 22/04/2026.

