## Supplementary material for "Collective protection drives human gut microbiota response to amoxicillin treatment": Suplementary figures and tables

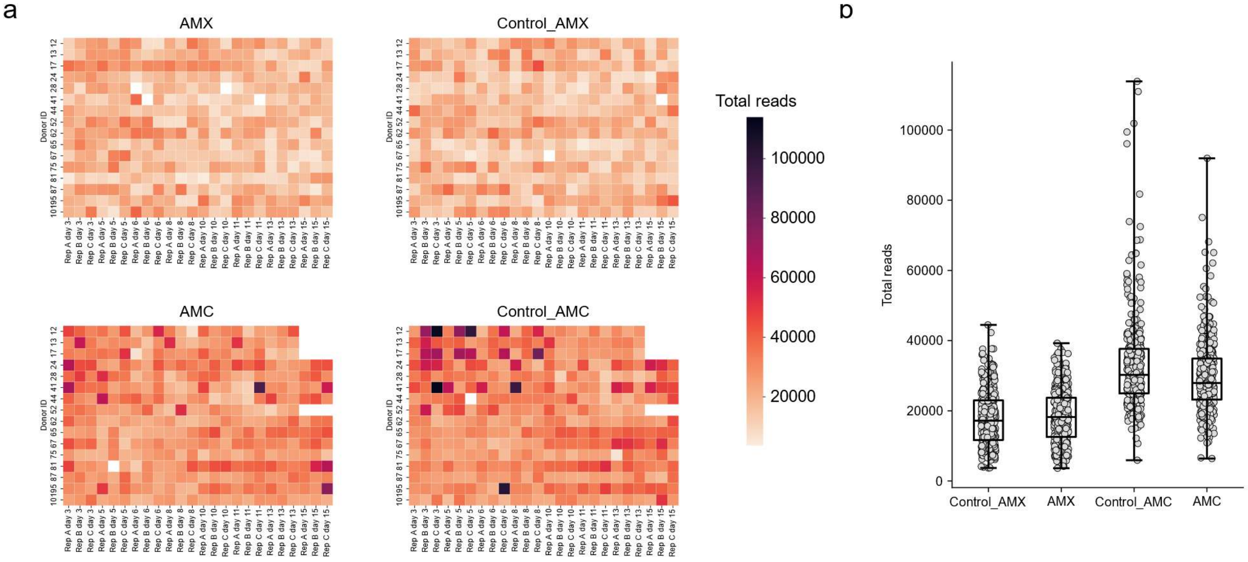


**Fig. S1**. Read counts of samples from both the AMX and AMC experiments. **a.** Heatmap showing total read counts per samples. Empty cells represent samples that were discarded out of the analysis for having less than 3500 reads. **b.** Distribution of read counts from both the AMX and AMC experiments.

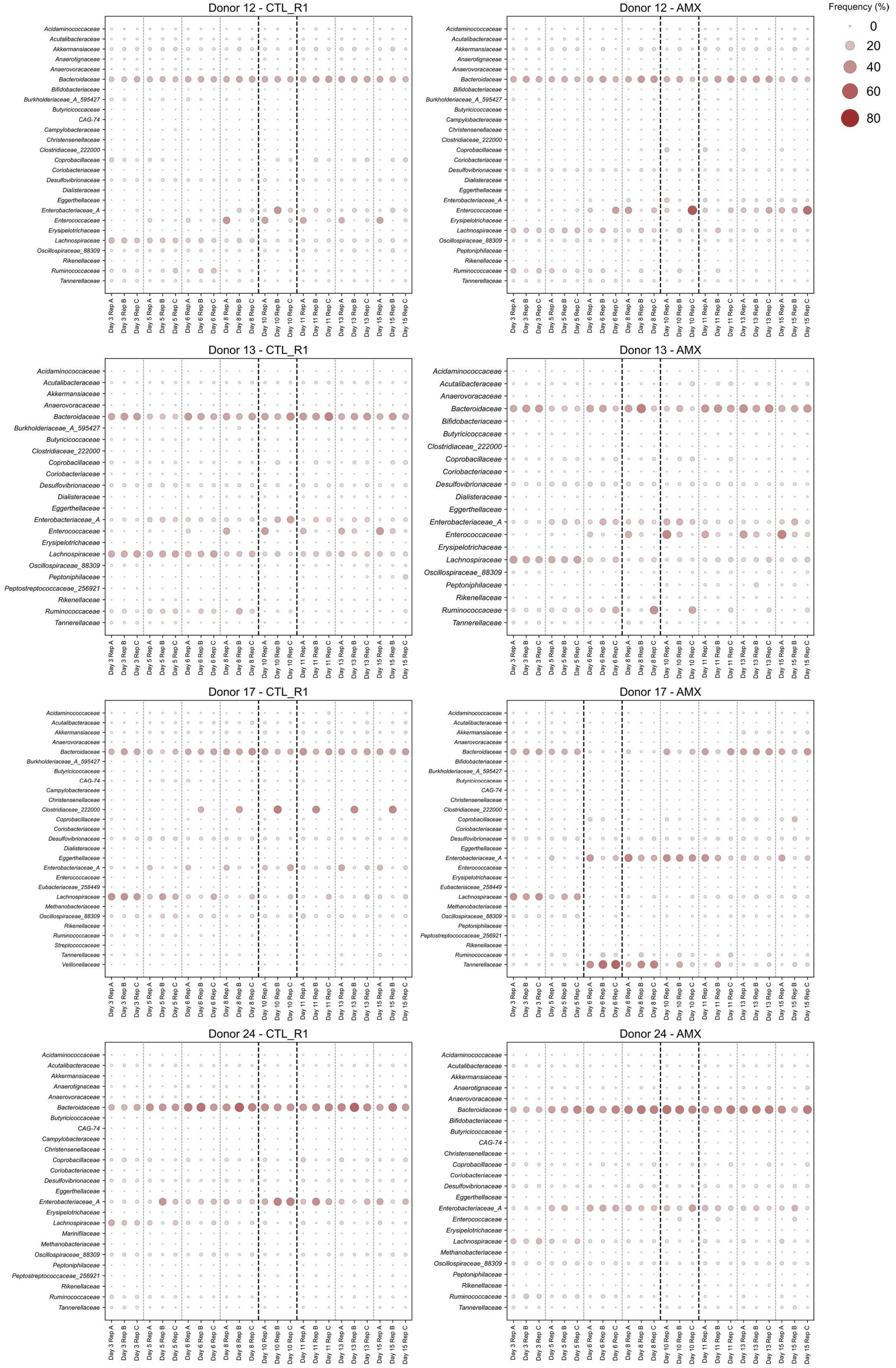
Frequency (%)
0
20
40
60
80
Donor 12 - CTL_R1
Acidaminococcaceae
Acutalibacteraceae
Akkermansiaceae
Anaerotignaceae
Anaerovoracaceae
Bacteroidaceae
Bifidobacteriaceae
Burkholderiaceae_A_595427
Butyricicoccaceae
CAG-74
Campylobacteraceae
Christensenellaceae
Clostridiaceae_222000
Coprobacillaceae
Coriobacteriaceae
Desulfovibrionaceae
Dialisteraceae
Eggerthellaceae
Enterobacteriaceae_A
Enterococcaceae
Erysipelotrichaceae
Lachnospiraceae
Oscillospiraceae_88309
Rikenellaceae
Ruminococcaceae
Tannerellaceae
Donor 12 - AMX
Acidaminococcaceae
Acutalibacteraceae
Akkermansiaceae
Anaerotignaceae
Anaerovoracaceae
Bacteroidaceae
Bifidobacteriaceae
Burkholderiaceae_A_595427
Butyricicoccaceae
Campylobacteraceae
Christensenellaceae
Clostridiaceae_222000
Coprobacillaceae
Coriobacteriaceae
Desulfovibrionaceae
Dialisteraceae
Eggerthellaceae
Enterobacteriaceae_A
Enterococcaceae
Erysipelotrichaceae
Lachnospiraceae
Oscillospiraceae_88309
Peptoniphilaceae
Rikenellaceae
Ruminococcaceae
Tannerellaceae
Donor 13 - CTL_R1
Acidaminococcaceae
Acutalibacteraceae
Akkermansiaceae
Anaerovoracaceae
Bacteroidaceae
Burkholderiaceae_A_595427
Butyricicoccaceae
Clostridiaceae_222000
Coprobacillaceae
Coriobacteriaceae
Desulfovibrionaceae
Dialisteraceae
Eggerthellaceae
Enterobacteriaceae_A
Enterococcaceae
Erysipelotrichaceae
Lachnospiraceae
Oscillospiraceae_88309
Peptoniphilaceae
Peptostreptococcaceae_256921
Rikenellaceae
Ruminococcaceae
Tannerellaceae
Donor 13 - AMX
Acidaminococcaceae
Acutalibacteraceae
Anaerovoracaceae
Bacteroidaceae
Bifidobacteriaceae
Butyricicoccaceae
Clostridiaceae_222000
Coprobacillaceae
Coriobacteriaceae
Desulfovibrionaceae
Dialisteraceae
Eggerthellaceae
Enterobacteriaceae_A
Enterococcaceae
Erysipelotrichaceae
Lachnospiraceae
Oscillospiraceae_88309
Peptoniphilaceae
Rikenellaceae
Ruminococcaceae
Tannerellaceae
Donor 17 - CTL_R1
Acidaminococcaceae
Acutalibacteraceae
Akkermansiaceae
Anaerovoracaceae
Bacteroidaceae
Burkholderiaceae_A_595427
Butyricicoccaceae
CAG-74
Campylobacteraceae
Christensenellaceae
Clostridiaceae_222000
Coprobacillaceae
Coriobacteriaceae
Desulfovibrionaceae
Dialisteraceae
Eggerthellaceae
Enterobacteriaceae_A
Enterococcaceae
Eubacteriaceae_258449
Lachnospiraceae
Methanobacteriaceae
Oscillospiraceae_88309
Rikenellaceae
Ruminococcaceae
Streptococcaceae
Tannerellaceae
Veillonellaceae
Donor 17 - AMX
Acidaminococcaceae
Acutalibacteraceae
Akkermansiaceae
Anaerovoracaceae
Bacteroidaceae
Bifidobacteriaceae
Burkholderiaceae_A_595427
Butyricicoccaceae
CAG-74
Christensenellaceae
Clostridiaceae_222000
Coprobacillaceae
Coriobacteriaceae
Desulfovibrionaceae
Eggerthellaceae
Enterobacteriaceae_A
Enterococcaceae
Erysipelotrichaceae
Eubacteriaceae_258449
Lachnospiraceae
Methanobacteriaceae
Oscillospiraceae_88309
Peptoniphilaceae
Peptostreptococcaceae_256921
Rikenellaceae
Ruminococcaceae
Tannerellaceae
Donor 24 - CTL_R1
Acidaminococcaceae
Acutalibacteraceae
Akkermansiaceae
Anaerotignaceae
Anaerovoracaceae
Bacteroidaceae
Butyricicoccaceae
CAG-74
Campylobacteraceae
Christensenellaceae
Coprobacillaceae
Coriobacteriaceae
Desulfovibrionaceae
Eggerthellaceae
Enterobacteriaceae_A
Erysipelotrichaceae
Lachnospiraceae
Marinifilaceae
Methanobacteriaceae
Oscillospiraceae_88309
Peptoniphilaceae
Peptostreptococcaceae_256921
Rikenellaceae
Ruminococcaceae
Tannerellaceae
Donor 24 - AMX
Acidaminococcaceae
Acutalibacteraceae
Akkermansiaceae
Anaerotignaceae
Anaerovoracaceae
Bacteroidaceae
Bifidobacteriaceae
Butyricicoccaceae
CAG-74
Christensenellaceae
Coprobacillaceae
Coriobacteriaceae
Desulfovibrionaceae
Eggerthellaceae
Enterobacteriaceae_A
Enterococcaceae
Erysipelotrichaceae
Lachnospiraceae
Methanobacteriaceae
Oscillospiraceae_88309
Peptoniphilaceae
Rikenellaceae
Ruminococcaceae
Tannerellaceae
Day 3 Rep A
Day 3 Rep B
Day 3 Rep C
Day 5 Rep A
Day 5 Rep B
Day 5 Rep C
Day 6 Rep A
Day 6 Rep B
Day 6 Rep C
Day 8 Rep A
Day 8 Rep B
Day 8 Rep C
Day 10 Rep A
Day 10 Rep B
Day 10 Rep C
Day 11 Rep A
Day 11 Rep B
Day 11 Rep C
Day 13 Rep A
Day 13 Rep B
Day 13 Rep C
Day 15 Rep A
Day 15 Rep B
Day 15 Rep C

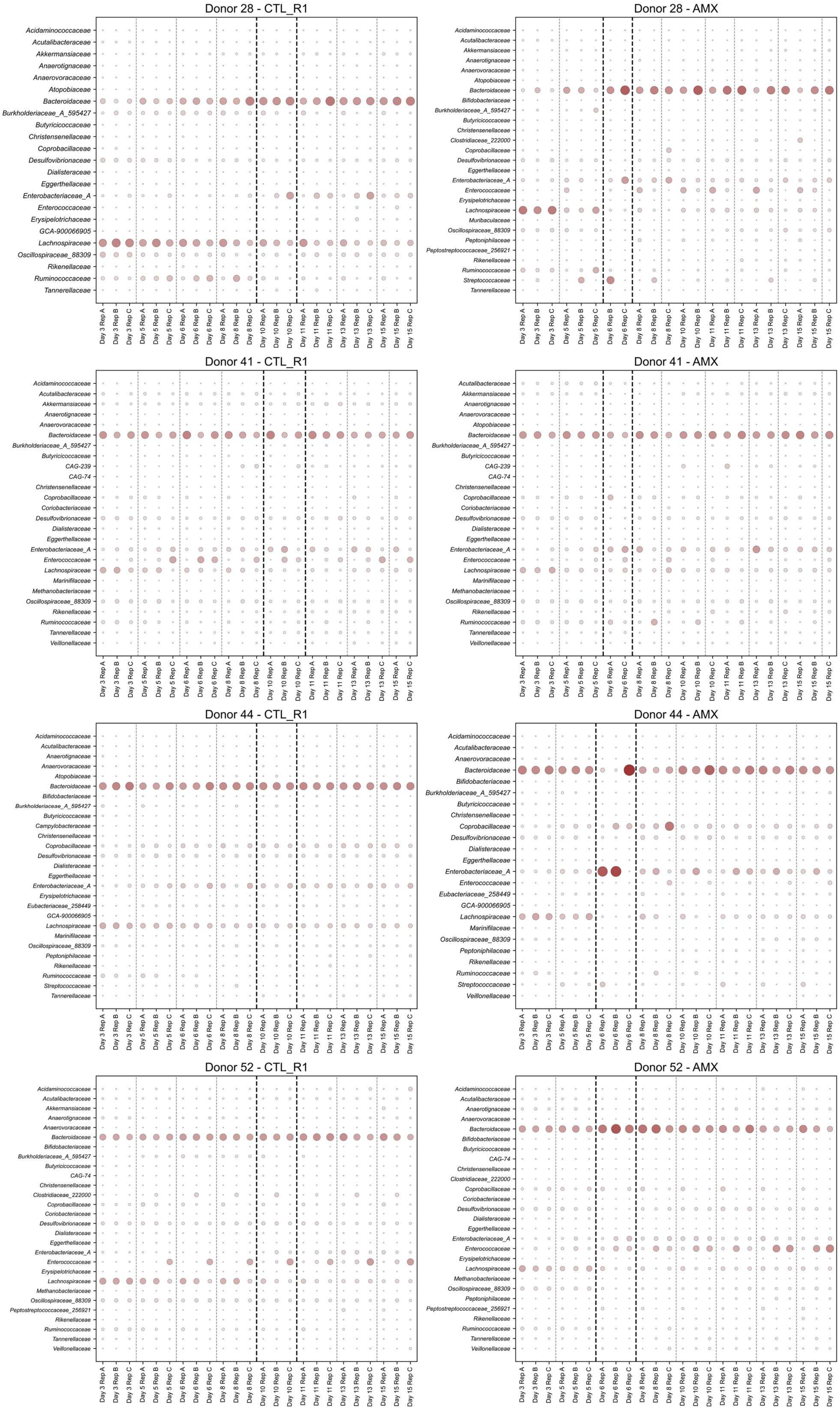
Donor 28 - CTL_R1
Acidaminococcaceae
Acutalibacteraceae
Akkermansiaceae
Anaerotignaceae
Anaerovoracaceae
Atopobiaceae
Bacteroidaceae
Burkholderiaceae_A_595427
Butyricicoccaceae
Christensenellaceae
Coprobacillaceae
Desulfovibrionaceae
Dialisteraceae
Eggerthellaceae
Enterobacteriaceae_A
Enterococcaceae
Erysipelotrichaceae
GCA-900066905
Lachnospiraceae
Oscillospiraceae_88309
Rikenellaceae
Ruminococcaceae
Tannerellaceae
Day 3 Rep A
Day 3 Rep B
Day 3 Rep C
Day 5 Rep A
Day 5 Rep B
Day 5 Rep C
Day 6 Rep A
Day 6 Rep B
Day 6 Rep C
Day 8 Rep A
Day 8 Rep B
Day 8 Rep C
Day 10 Rep A
Day 10 Rep B
Day 10 Rep C
Day 11 Rep A
Day 11 Rep B
Day 11 Rep C
Day 13 Rep A
Day 13 Rep B
Day 13 Rep C
Day 15 Rep A
Day 15 Rep B
Day 15 Rep C
Donor 28 - AMX
Acidaminococcaceae
Acutalibacteraceae
Akkermansiaceae
Anaerotignaceae
Anaerovoracaceae
Atopobiaceae
Bacteroidaceae
Bifidobacteriaceae
Burkholderiaceae_A_595427
Butyricicoccaceae
Christensenellaceae
Clostridiaceae_222000
Coprobacillaceae
Desulfovibrionaceae
Eggerthellaceae
Enterobacteriaceae_A
Enterococcaceae
Erysipelotrichaceae
Lachnospiraceae
Muribaculaceae
Oscillospiraceae_88309
Peptoniphilaceae
Peptostreptococcaceae_256921
Rikenellaceae
Ruminococcaceae
Streptococcaceae
Tannerellaceae
Day 3 Rep A
Day 3 Rep B
Day 3 Rep C
Day 5 Rep A
Day 5 Rep B
Day 5 Rep C
Day 6 Rep A
Day 6 Rep B
Day 6 Rep C
Day 8 Rep A
Day 8 Rep B
Day 8 Rep C
Day 10 Rep A
Day 10 Rep B
Day 10 Rep C
Day 11 Rep A
Day 11 Rep B
Day 11 Rep C
Day 13 Rep A
Day 13 Rep B
Day 13 Rep C
Day 15 Rep A
Day 15 Rep B
Day 15 Rep C
Donor 41 - CTL_R1
Acidaminococcaceae
Acutalibacteraceae
Akkermansiaceae
Anaerotignaceae
Anaerovoracaceae
Bacteroidaceae
Burkholderiaceae_A_595427
Butyricicoccaceae
CAG-239
CAG-74
Christensenellaceae
Coprobacillaceae
Coriobacteriaceae
Desulfovibrionaceae
Dialisteraceae
Eggerthellaceae
Enterobacteriaceae_A
Enterococcaceae
Lachnospiraceae
Marinifilaceae
Methanobacteriaceae
Oscillospiraceae_88309
Rikenellaceae
Ruminococcaceae
Tannerellaceae
Veillonellaceae
Day 3 Rep A
Day 3 Rep B
Day 3 Rep C
Day 5 Rep A
Day 5 Rep B
Day 5 Rep C
Day 6 Rep A
Day 6 Rep B
Day 6 Rep C
Day 8 Rep A
Day 8 Rep B
Day 8 Rep C
Day 10 Rep A
Day 10 Rep B
Day 10 Rep C
Day 11 Rep A
Day 11 Rep B
Day 11 Rep C
Day 13 Rep A
Day 13 Rep B
Day 13 Rep C
Day 15 Rep A
Day 15 Rep B
Day 15 Rep C
Donor 41 - AMX
Acutalibacteraceae
Akkermansiaceae
Anaerotignaceae
Anaerovoracaceae
Atopobiaceae
Bacteroidaceae
Burkholderiaceae_A_595427
Butyricicoccaceae
CAG-239
CAG-74
Christensenellaceae
Coprobacillaceae
Coriobacteriaceae
Desulfovibrionaceae
Dialisteraceae
Eggerthellaceae
Enterobacteriaceae_A
Enterococcaceae
Lachnospiraceae
Marinifilaceae
Methanobacteriaceae
Oscillospiraceae_88309
Rikenellaceae
Ruminococcaceae
Tannerellaceae
Veillonellaceae
Day 3 Rep A
Day 3 Rep B
Day 3 Rep C
Day 5 Rep A
Day 5 Rep B
Day 5 Rep C
Day 6 Rep A
Day 6 Rep B
Day 6 Rep C
Day 8 Rep A
Day 8 Rep B
Day 8 Rep C
Day 10 Rep A
Day 10 Rep B
Day 10 Rep C
Day 11 Rep A
Day 11 Rep B
Day 11 Rep C
Day 13 Rep A
Day 13 Rep B
Day 13 Rep C
Day 15 Rep A
Day 15 Rep B
Day 15 Rep C
Donor 44 - CTL_R1
Acidaminococcaceae
Acutalibacteraceae
Anaerotignaceae
Anaerovoracaceae
Atopobiaceae
Bacteroidaceae
Bifidobacteriaceae
Burkholderiaceae_A_595427
Butyricicoccaceae
Campylobacteraceae
Christensenellaceae
Coprobacillaceae
Desulfovibrionaceae
Dialisteraceae
Eggerthellaceae
Enterobacteriaceae_A
Erysipelotrichaceae
Eubacteriaceae_258449
GCA-900066905
Lachnospiraceae
Marinifilaceae
Oscillospiraceae_88309
Peptoniphilaceae
Rikenellaceae
Ruminococcaceae
Streptococcaceae
Tannerellaceae
Day 3 Rep A
Day 3 Rep B
Day 3 Rep C
Day 5 Rep A
Day 5 Rep B
Day 5 Rep C
Day 6 Rep A
Day 6 Rep B
Day 6 Rep C
Day 8 Rep A
Day 8 Rep B
Day 8 Rep C
Day 10 Rep A
Day 10 Rep B
Day 10 Rep C
Day 11 Rep A
Day 11 Rep B
Day 11 Rep C
Day 13 Rep A
Day 13 Rep B
Day 13 Rep C
Day 15 Rep A
Day 15 Rep B
Day 15 Rep C
Donor 44 - AMX
Acidaminococcaceae
Acutalibacteraceae
Anaerovoracaceae
Bacteroidaceae
Bifidobacteriaceae
Burkholderiaceae_A_595427
Butyricicoccaceae
Christensenellaceae
Coprobacillaceae
Desulfovibrionaceae
Dialisteraceae
Eggerthellaceae
Enterobacteriaceae_A
Enterococcaceae
Eubacteriaceae_258449
GCA-900066905
Lachnospiraceae
Marinifilaceae
Oscillospiraceae_88309
Peptoniphilaceae
Rikenellaceae
Ruminococcaceae
Streptococcaceae
Veillonellaceae
Day 3 Rep A
Day 3 Rep B
Day 3 Rep C
Day 5 Rep A
Day 5 Rep B
Day 5 Rep C
Day 6 Rep A
Day 6 Rep B
Day 6 Rep C
Day 8 Rep A
Day 8 Rep B
Day 8 Rep C
Day 10 Rep A
Day 10 Rep B
Day 10 Rep C
Day 11 Rep A
Day 11 Rep B
Day 11 Rep C
Day 13 Rep A
Day 13 Rep B
Day 13 Rep C
Day 15 Rep A
Day 15 Rep B
Day 15 Rep C
Donor 52 - CTL_R1
Acidaminococcaceae
Acutalibacteraceae
Akkermansiaceae
Anaerotignaceae
Anaerovoracaceae
Bacteroidaceae
Bifidobacteriaceae
Burkholderiaceae_A_595427
Butyricicoccaceae
CAG-74
Christensenellaceae
Clostridiaceae_222000
Coprobacillaceae
Coriobacteriaceae
Desulfovibrionaceae
Dialisteraceae
Eggerthellaceae
Enterobacteriaceae_A
Enterococcaceae
Erysipelotrichaceae
Lachnospiraceae
Methanobacteriaceae
Oscillospiraceae_88309
Peptostreptococcaceae_256921
Rikenellaceae
Ruminococcaceae
Tannerellaceae
Veillonellaceae
Day 3 Rep A
Day 3 Rep B
Day 3 Rep C
Day 5 Rep A
Day 5 Rep B
Day 5 Rep C
Day 6 Rep A
Day 6 Rep B
Day 6 Rep C
Day 8 Rep A
Day 8 Rep B
Day 8 Rep C
Day 10 Rep A
Day 10 Rep B
Day 10 Rep C
Day 11 Rep A
Day 11 Rep B
Day 11 Rep C
Day 13 Rep A
Day 13 Rep B
Day 13 Rep C
Day 15 Rep A
Day 15 Rep B
Day 15 Rep C
Donor 52 - AMX
Acidaminococcaceae
Acutalibacteraceae
Anaerotignaceae
Anaerovoracaceae
Bacteroidaceae
Bifidobacteriaceae
Butyricicoccaceae
CAG-74
Christensenellaceae
Clostridiaceae_222000
Coprobacillaceae
Coriobacteriaceae
Desulfovibrionaceae
Dialisteraceae
Eggerthellaceae
Enterobacteriaceae_A
Enterococcaceae
Erysipelotrichaceae
Lachnospiraceae
Methanobacteriaceae
Oscillospiraceae_88309
Peptoniphilaceae
Peptostreptococcaceae_256921
Rikenellaceae
Ruminococcaceae
Tannerellaceae
Veillonellaceae
Day 3 Rep A
Day 3 Rep B
Day 3 Rep C
Day 5 Rep A
Day 5 Rep B
Day 5 Rep C
Day 6 Rep A
Day 6 Rep B
Day 6 Rep C
Day 8 Rep A
Day 8 Rep B
Day 8 Rep C
Day 10 Rep A
Day 10 Rep B
Day 10 Rep C
Day 11 Rep A
Day 11 Rep B
Day 11 Rep C
Day 13 Rep A
Day 13 Rep B
Day 13 Rep C
Day 15 Rep A
Day 15 Rep B
Day 15 Rep C

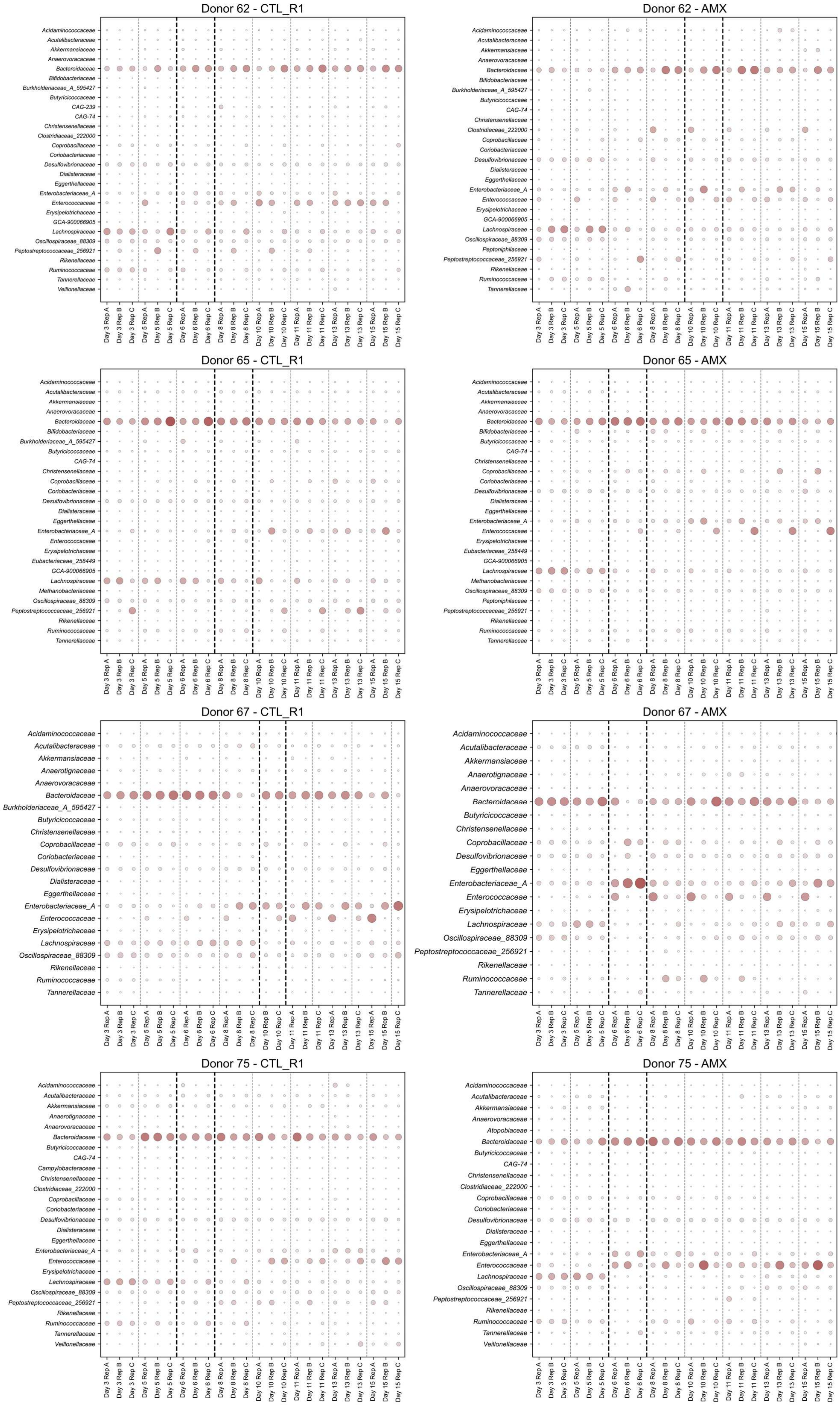

Donor 62 - CTL_R1
Acidaminococcaceae
Acutalibacteraceae
Akkermansiaceae
Anaerovoracaceae
Bacteroidaceae
Bifidobacteriaceae
Burkholderiaceae_A_595427
Butyricicoccaceae
CAG-239
CAG-74
Christensenellaceae
Clostridiaceae_222000
Coprobacillaceae
Coriobacteriaceae
Desulfovibrionaceae
Dialisteraceae
Eggerthellaceae
Enterobacteriaceae_A
Enterococcaceae
Erysipelotrichaceae
GCA-900066905
Lachnospiraceae
Oscillospiraceae_88309
Peptostreptococcaceae_256921
Rikenellaceae
Ruminococcaceae
Tannerellaceae
Veillonellaceae
Day 3 Rep A
Day 3 Rep B
Day 3 Rep C
Day 5 Rep A
Day 5 Rep B
Day 5 Rep C
Day 6 Rep A
Day 6 Rep B
Day 6 Rep C
Day 8 Rep A
Day 8 Rep B
Day 8 Rep C
Day 10 Rep A
Day 10 Rep B
Day 10 Rep C
Day 11 Rep A
Day 11 Rep B
Day 11 Rep C
Day 13 Rep A
Day 13 Rep B
Day 13 Rep C
Day 15 Rep A
Day 15 Rep B
Day 15 Rep C
Donor 62 - AMX
Acidaminococcaceae
Acutalibacteraceae
Akkermansiaceae
Anaerovoracaceae
Bacteroidaceae
Bifidobacteriaceae
Burkholderiaceae_A_595427
Butyricicoccaceae
CAG-74
Christensenellaceae
Clostridiaceae_222000
Coprobacillaceae
Coriobacteriaceae
Desulfovibrionaceae
Dialisteraceae
Eggerthellaceae
Enterobacteriaceae_A
Enterococcaceae
Erysipelotrichaceae
GCA-900066905
Lachnospiraceae
Oscillospiraceae_88309
Peptoniphilaceae
Peptostreptococcaceae_256921
Rikenellaceae
Ruminococcaceae
Tannerellaceae
Donor 65 - CTL_R1
Acidaminococcaceae
Acutalibacteraceae
Akkermansiaceae
Anaerovoracaceae
Bacteroidaceae
Bifidobacteriaceae
Burkholderiaceae_A_595427
Butyricicoccaceae
CAG-74
Christensenellaceae
Coprobacillaceae
Coriobacteriaceae
Desulfovibrionaceae
Dialisteraceae
Eggerthellaceae
Enterobacteriaceae_A
Enterococcaceae
Erysipelotrichaceae
Eubacteriaceae_258449
GCA-900066905
Lachnospiraceae
Methanobacteriaceae
Oscillospiraceae_88309
Peptostreptococcaceae_256921
Rikenellaceae
Ruminococcaceae
Tannerellaceae
Donor 65 - AMX
Acidaminococcaceae
Acutalibacteraceae
Akkermansiaceae
Anaerovoracaceae
Bacteroidaceae
Bifidobacteriaceae
Butyricicoccaceae
CAG-74
Christensenellaceae
Coprobacillaceae
Coriobacteriaceae
Desulfovibrionaceae
Dialisteraceae
Eggerthellaceae
Enterobacteriaceae_A
Enterococcaceae
Erysipelotrichaceae
Eubacteriaceae_258449
GCA-900066905
Lachnospiraceae
Methanobacteriaceae
Oscillospiraceae_88309
Peptoniphilaceae
Peptostreptococcaceae_256921
Rikenellaceae
Ruminococcaceae
Tannerellaceae
Donor 67 - CTL_R1
Acidaminococcaceae
Acutalibacteraceae
Akkermansiaceae
Anaerotignaceae
Anaerovoracaceae
Bacteroidaceae
Burkholderiaceae_A_595427
Butyricicoccaceae
Christensenellaceae
Coprobacillaceae
Coriobacteriaceae
Desulfovibrionaceae
Dialisteraceae
Eggerthellaceae
Enterobacteriaceae_A
Enterococcaceae
Erysipelotrichaceae
Lachnospiraceae
Oscillospiraceae_88309
Rikenellaceae
Ruminococcaceae
Tannerellaceae
Donor 67 - AMX
Acidaminococcaceae
Acutalibacteraceae
Akkermansiaceae
Anaerotignaceae
Anaerovoracaceae
Bacteroidaceae
Butyricicoccaceae
Christensenellaceae
Coprobacillaceae
Desulfovibrionaceae
Eggerthellaceae
Enterobacteriaceae_A
Enterococcaceae
Erysipelotrichaceae
Lachnospiraceae
Oscillospiraceae_88309
Peptostreptococcaceae_256921
Rikenellaceae
Ruminococcaceae
Tannerellaceae
Donor 75 - CTL_R1
Acidaminococcaceae
Acutalibacteraceae
Akkermansiaceae
Anaerotignaceae
Anaerovoracaceae
Bacteroidaceae
Butyricicoccaceae
CAG-74
Campylobacteraceae
Christensenellaceae
Clostridiaceae_222000
Coprobacillaceae
Coriobacteriaceae
Desulfovibrionaceae
Dialisteraceae
Eggerthellaceae
Enterobacteriaceae_A
Enterococcaceae
Erysipelotrichaceae
Lachnospiraceae
Oscillospiraceae_88309
Peptostreptococcaceae_256921
Rikenellaceae
Ruminococcaceae
Tannerellaceae
Veillonellaceae
Donor 75 - AMX
Acidaminococcaceae
Acutalibacteraceae
Akkermansiaceae
Anaerovoracaceae
Atopobiaceae
Bacteroidaceae
Butyricicoccaceae
CAG-74
Christensenellaceae
Clostridiaceae_222000
Coprobacillaceae
Coriobacteriaceae
Desulfovibrionaceae
Dialisteraceae
Eggerthellaceae
Enterobacteriaceae_A
Enterococcaceae
Lachnospiraceae
Oscillospiraceae_88309
Peptostreptococcaceae_256921
Rikenellaceae
Ruminococcaceae
Tannerellaceae
Veillonellaceae

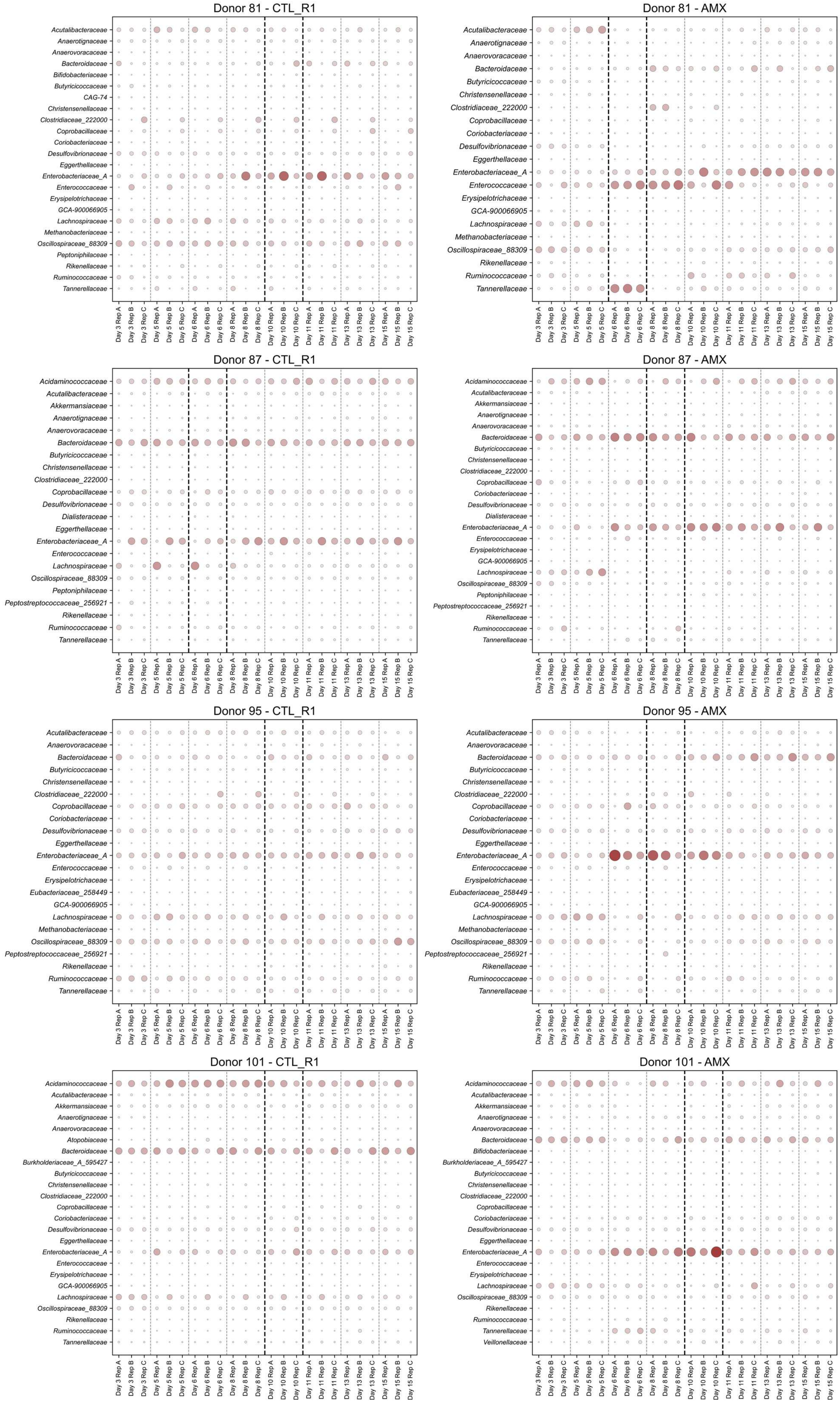
Donor 81 - CTL_R1
Acutalibacteraceae
Anaerotignaceae
Anaerovoracaceae
Bacteroidaceae
Bifidobacteriaceae
Butyricicoccaceae
CAG-74
Christensenellaceae
Clostridiaceae_222000
Coprobacillaceae
Coriobacteriaceae
Desulfovibrionaceae
Eggerthellaceae
Enterobacteriaceae_A
Enterococcaceae
Erysipelotrichaceae
GCA-900066905
Lachnospiraceae
Methanobacteriaceae
Oscillospiraceae_88309
Peptoniphilaceae
Rikenellaceae
Ruminococcaceae
Tannerellaceae
Day 3 Rep A
Day 3 Rep B
Day 3 Rep C
Day 5 Rep A
Day 5 Rep B
Day 5 Rep C
Day 6 Rep A
Day 6 Rep B
Day 6 Rep C
Day 8 Rep A
Day 8 Rep B
Day 8 Rep C
Day 10 Rep A
Day 10 Rep B
Day 10 Rep C
Day 11 Rep A
Day 11 Rep B
Day 11 Rep C
Day 13 Rep A
Day 13 Rep B
Day 13 Rep C
Day 15 Rep A
Day 15 Rep B
Day 15 Rep C
Donor 81 - AMX
Acutalibacteraceae
Anaerotignaceae
Anaerovoracaceae
Bacteroidaceae
Butyricicoccaceae
Christensenellaceae
Clostridiaceae_222000
Coprobacillaceae
Coriobacteriaceae
Desulfovibrionaceae
Eggerthellaceae
Enterobacteriaceae_A
Enterococcaceae
Erysipelotrichaceae
GCA-900066905
Lachnospiraceae
Methanobacteriaceae
Oscillospiraceae_88309
Rikenellaceae
Ruminococcaceae
Tannerellaceae
Day 3 Rep A
Day 3 Rep B
Day 3 Rep C
Day 5 Rep A
Day 5 Rep B
Day 5 Rep C
Day 6 Rep A
Day 6 Rep B
Day 6 Rep C
Day 8 Rep A
Day 8 Rep B
Day 8 Rep C
Day 10 Rep A
Day 10 Rep B
Day 10 Rep C
Day 11 Rep A
Day 11 Rep B
Day 11 Rep C
Day 13 Rep A
Day 13 Rep B
Day 13 Rep C
Day 15 Rep A
Day 15 Rep B
Day 15 Rep C
Donor 87 - CTL_R1
Acidaminococcaceae
Acutalibacteraceae
Akkermansiaceae
Anaerotignaceae
Anaerovoracaceae
Bacteroidaceae
Butyricicoccaceae
Christensenellaceae
Clostridiaceae_222000
Coprobacillaceae
Desulfovibrionaceae
Dialisteraceae
Eggerthellaceae
Enterobacteriaceae_A
Enterococcaceae
Lachnospiraceae
Oscillospiraceae_88309
Peptoniphilaceae
Peptostreptococcaceae_256921
Rikenellaceae
Ruminococcaceae
Tannerellaceae
Day 3 Rep A
Day 3 Rep B
Day 3 Rep C
Day 5 Rep A
Day 5 Rep B
Day 5 Rep C
Day 6 Rep A
Day 6 Rep B
Day 6 Rep C
Day 8 Rep A
Day 8 Rep B
Day 8 Rep C
Day 10 Rep A
Day 10 Rep B
Day 10 Rep C
Day 11 Rep A
Day 11 Rep B
Day 11 Rep C
Day 13 Rep A
Day 13 Rep B
Day 13 Rep C
Day 15 Rep A
Day 15 Rep B
Day 15 Rep C
Donor 87 - AMX
Acidaminococcaceae
Acutalibacteraceae
Akkermansiaceae
Anaerotignaceae
Anaerovoracaceae
Bacteroidaceae
Butyricicoccaceae
Christensenellaceae
Clostridiaceae_222000
Coprobacillaceae
Coriobacteriaceae
Desulfovibrionaceae
Dialisteraceae
Enterobacteriaceae_A
Enterococcaceae
Erysipelotrichaceae
GCA-900066905
Lachnospiraceae
Oscillospiraceae_88309
Peptoniphilaceae
Peptostreptococcaceae_256921
Rikenellaceae
Ruminococcaceae
Tannerellaceae
Day 3 Rep A
Day 3 Rep B
Day 3 Rep C
Day 5 Rep A
Day 5 Rep B
Day 5 Rep C
Day 6 Rep A
Day 6 Rep B
Day 6 Rep C
Day 8 Rep A
Day 8 Rep B
Day 8 Rep C
Day 10 Rep A
Day 10 Rep B
Day 10 Rep C
Day 11 Rep A
Day 11 Rep B
Day 11 Rep C
Day 13 Rep A
Day 13 Rep B
Day 13 Rep C
Day 15 Rep A
Day 15 Rep B
Day 15 Rep C
Donor 95 - CTL_R1
Acutalibacteraceae
Anaerovoracaceae
Bacteroidaceae
Butyricicoccaceae
Christensenellaceae
Clostridiaceae_222000
Coprobacillaceae
Coriobacteriaceae
Desulfovibrionaceae
Eggerthellaceae
Enterobacteriaceae_A
Enterococcaceae
Erysipelotrichaceae
Eubacteriaceae_258449
GCA-900066905
Lachnospiraceae
Methanobacteriaceae
Oscillospiraceae_88309
Peptostreptococcaceae_256921
Rikenellaceae
Ruminococcaceae
Tannerellaceae
Day 3 Rep A
Day 3 Rep B
Day 3 Rep C
Day 5 Rep A
Day 5 Rep B
Day 5 Rep C
Day 6 Rep A
Day 6 Rep B
Day 6 Rep C
Day 8 Rep A
Day 8 Rep B
Day 8 Rep C
Day 10 Rep A
Day 10 Rep B
Day 10 Rep C
Day 11 Rep A
Day 11 Rep B
Day 11 Rep C
Day 13 Rep A
Day 13 Rep B
Day 13 Rep C
Day 15 Rep A
Day 15 Rep B
Day 15 Rep C
Donor 95 - AMX
Acutalibacteraceae
Anaerovoracaceae
Bacteroidaceae
Butyricicoccaceae
Christensenellaceae
Clostridiaceae_222000
Coprobacillaceae
Coriobacteriaceae
Desulfovibrionaceae
Eggerthellaceae
Enterobacteriaceae_A
Enterococcaceae
Erysipelotrichaceae
Eubacteriaceae_258449
GCA-900066905
Lachnospiraceae
Methanobacteriaceae
Oscillospiraceae_88309
Peptostreptococcaceae_256921
Rikenellaceae
Ruminococcaceae
Tannerellaceae
Day 3 Rep A
Day 3 Rep B
Day 3 Rep C
Day 5 Rep A
Day 5 Rep B
Day 5 Rep C
Day 6 Rep A
Day 6 Rep B
Day 6 Rep C
Day 8 Rep A
Day 8 Rep B
Day 8 Rep C
Day 10 Rep A
Day 10 Rep B
Day 10 Rep C
Day 11 Rep A
Day 11 Rep B
Day 11 Rep C
Day 13 Rep A
Day 13 Rep B
Day 13 Rep C
Day 15 Rep A
Day 15 Rep B
Day 15 Rep C
Donor 101 - CTL_R1
Acidaminococcaceae
Acutalibacteraceae
Akkermansiaceae
Anaerotignaceae
Anaerovoracaceae
Atopobiaceae
Bacteroidaceae
Burkholderiaceae_A_595427
Butyricicoccaceae
Christensenellaceae
Clostridiaceae_222000
Coprobacillaceae
Coriobacteriaceae
Desulfovibrionaceae
Eggerthellaceae
Enterobacteriaceae_A
Enterococcaceae
Erysipelotrichaceae
GCA-900066905
Lachnospiraceae
Oscillospiraceae_88309
Rikenellaceae
Ruminococcaceae
Tannerellaceae
Day 3 Rep A
Day 3 Rep B
Day 3 Rep C
Day 5 Rep A
Day 5 Rep B
Day 5 Rep C
Day 6 Rep A
Day 6 Rep B
Day 6 Rep C
Day 8 Rep A
Day 8 Rep B
Day 8 Rep C
Day 10 Rep A
Day 10 Rep B
Day 10 Rep C
Day 11 Rep A
Day 11 Rep B
Day 11 Rep C
Day 13 Rep A
Day 13 Rep B
Day 13 Rep C
Day 15 Rep A
Day 15 Rep B
Day 15 Rep C
Donor 101 - AMX
Acidaminococcaceae
Acutalibacteraceae
Akkermansiaceae
Anaerotignaceae
Anaerovoracaceae
Bacteroidaceae
Bifidobacteriaceae
Burkholderiaceae_A_595427
Butyricicoccaceae
Christensenellaceae
Clostridiaceae_222000
Coprobacillaceae
Coriobacteriaceae
Desulfovibrionaceae
Eggerthellaceae
Enterobacteriaceae_A
Enterococcaceae
Erysipelotrichaceae
Lachnospiraceae
Oscillospiraceae_88309
Rikenellaceae
Ruminococcaceae
Tannerellaceae
Veillonellaceae
Day 3 Rep A
Day 3 Rep B
Day 3 Rep C
Day 5 Rep A
Day 5 Rep B
Day 5 Rep C
Day 6 Rep A
Day 6 Rep B
Day 6 Rep C
Day 8 Rep A
Day 8 Rep B
Day 8 Rep C
Day 10 Rep A
Day 10 Rep B
Day 10 Rep C
Day 11 Rep A
Day 11 Rep B
Day 11 Rep C
Day 13 Rep A
Day 13 Rep B
Day 13 Rep C
Day 15 Rep A
Day 15 Rep B
Day 15 Rep C

**Fig. S2**. Frequencies of most represented OTUs per donors during the AMX experiment. Shown are microbiotas' variations in the Control (CTL_R1) and AMX-treated chambers. Sampling days which have their boundaries in bold indicate the day of maximum perturbation (median perturbation strength for the replicates) observed for a given triplicate within the treatment window (days 5-10).

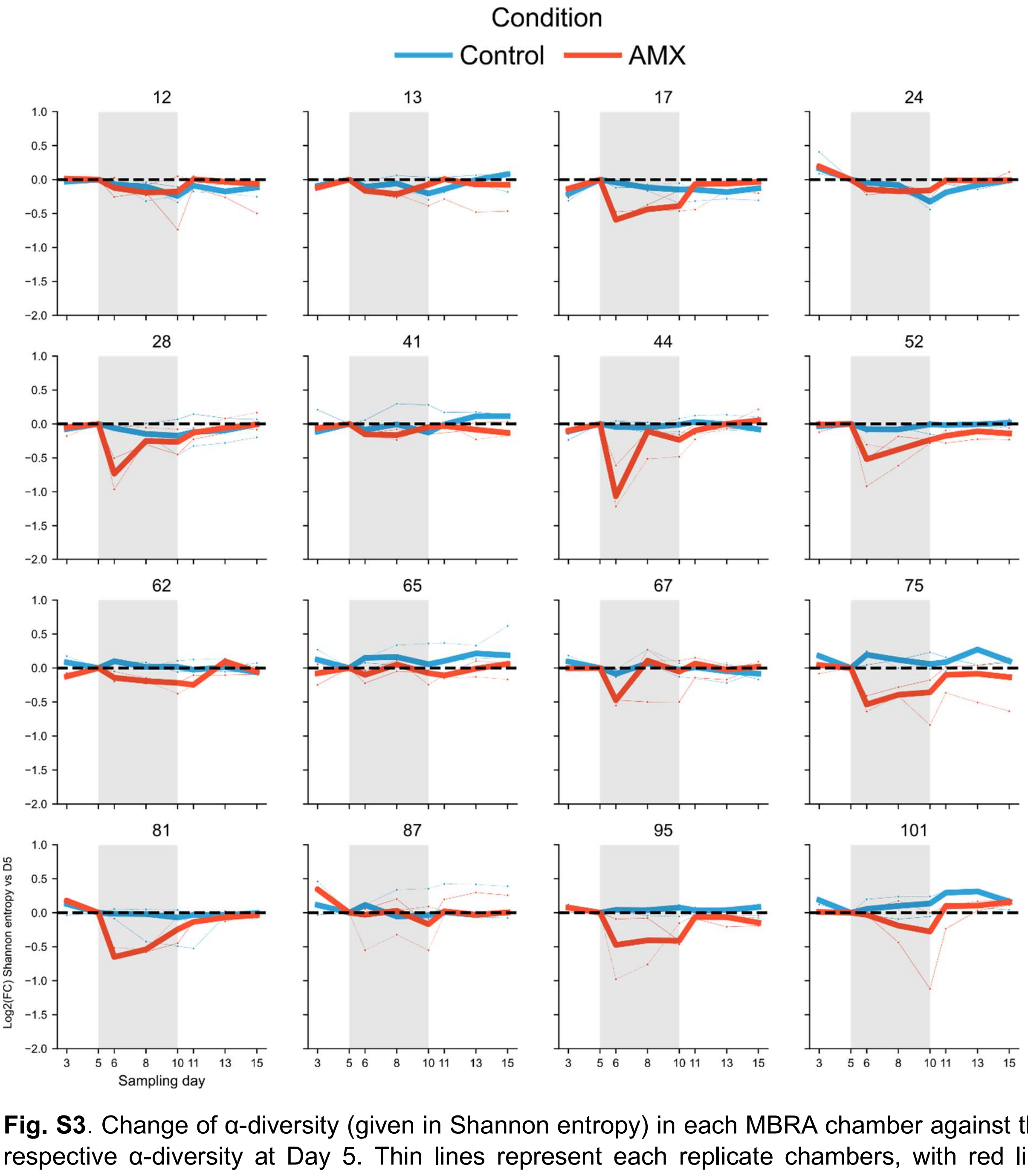


**Fig. S3**. Change of α-diversity (given in Shannon entropy) in each MBRA chamber against their respective α-diversity at Day 5. Thin lines represent each replicate chambers, with red lines representing AMX-treated chambers and blue lines representing Control chambers. Bold lines are median of the triplicate chambers. The grey rectangle corresponds to the treatment window (days 5-10). The bold dark line is a reference for log2(FC) = 0. Numbers above each plot refer to the ID of the donors. Data shown for Donors 24 and 81 are the same as in Fig. 1d.

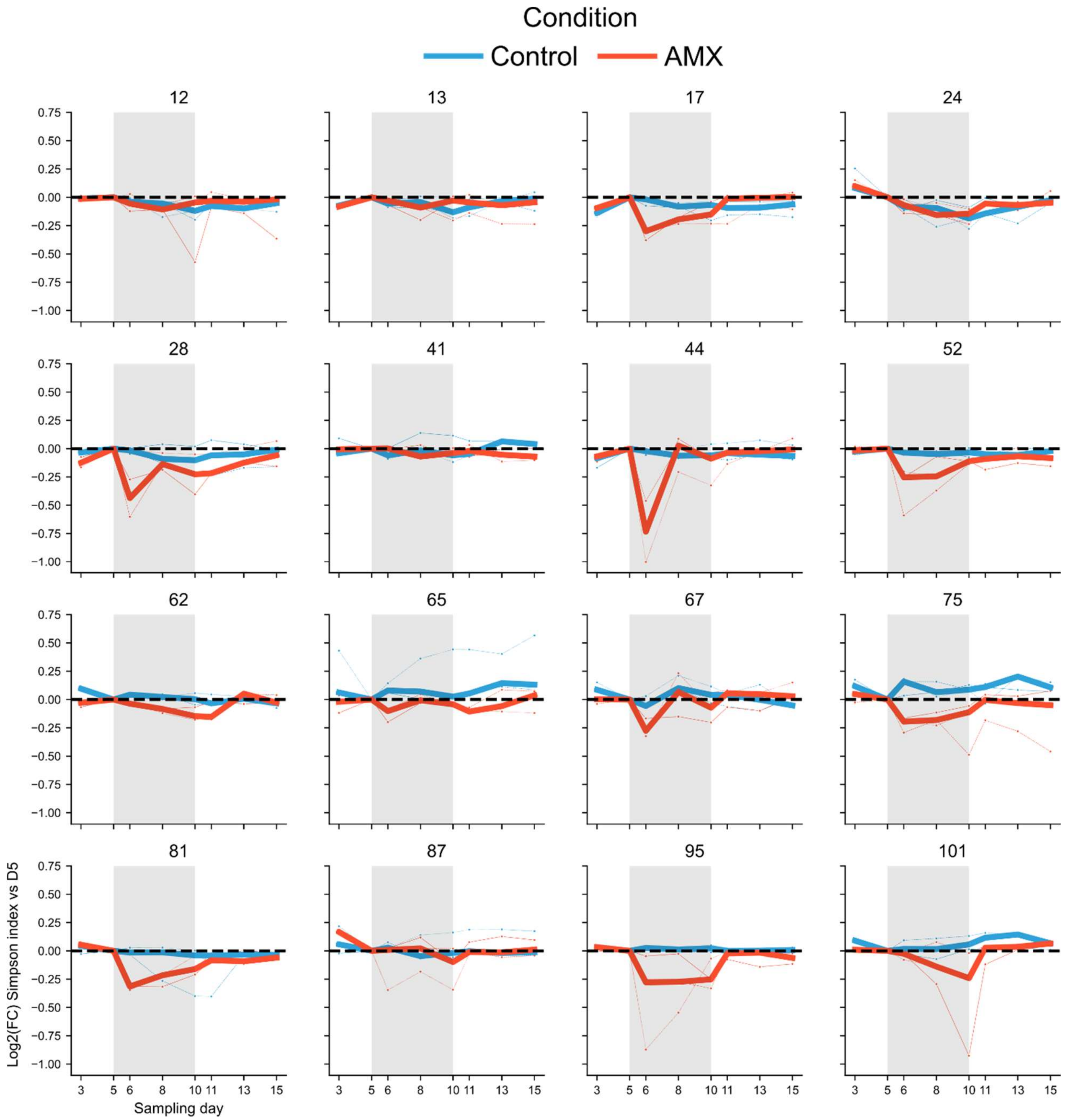


**Fig. S4**. Change of α-diversity (given in Simpson index) in each MBRA chamber against their respective α-diversity at Day 5. Thin lines represent each replicate chambers, with red lines representing AMX-treated chambers and blue lines representing Control chambers. Bold lines are median of the triplicate chambers. The grey rectangle corresponds to the treatment window (days 5-10). The bold dark line is a reference for log2(FC) = 0.

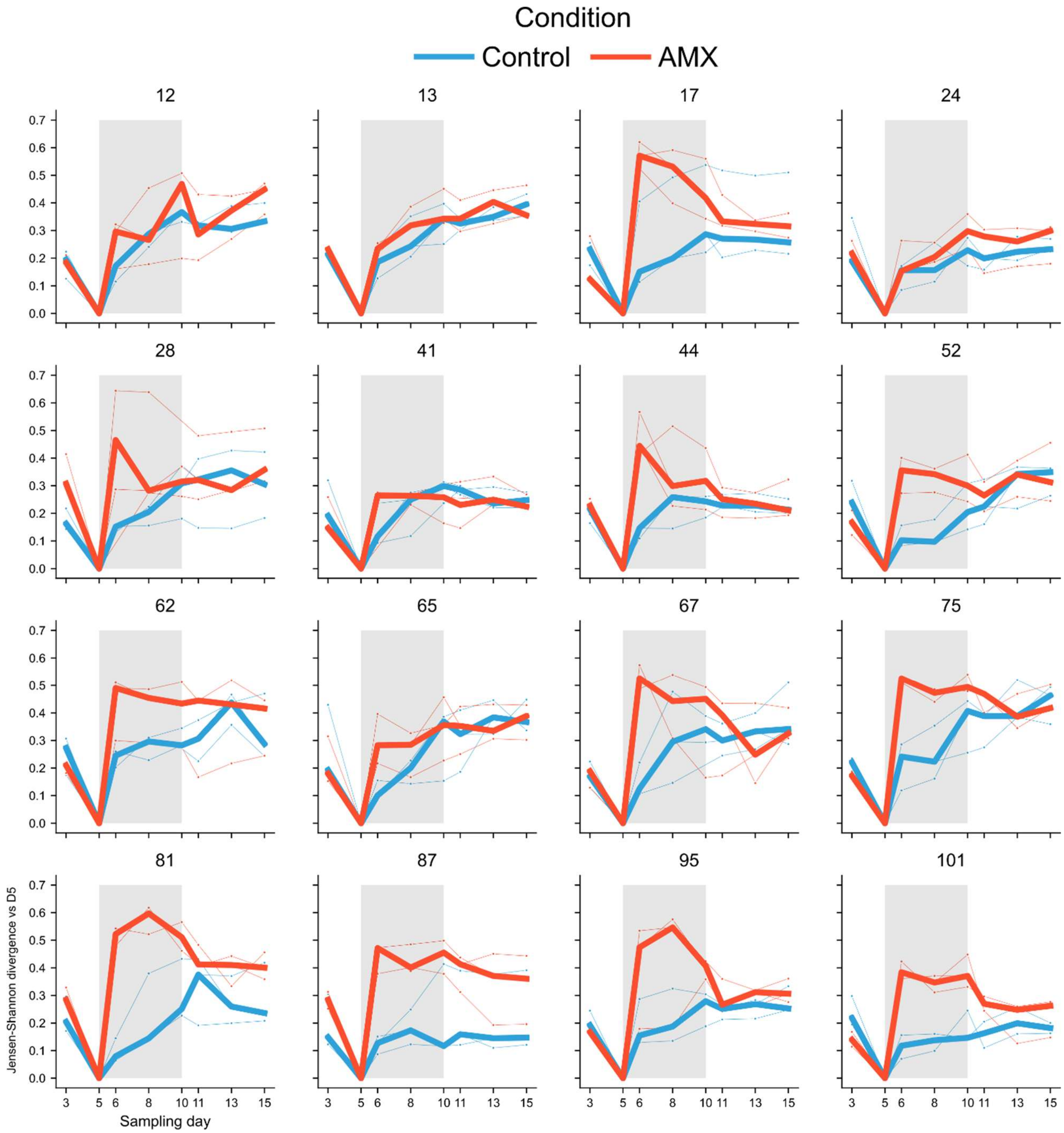


**Fig. S5**. Evolution of β-diversity in each MBRA chamber, given as their respective Jensen-Shannon divergence against their composition on Day 5. Thin lines represent each replicate chambers, with red lines representing AMX-treated chambers and blue lines representing Control chambers. Bold lines are median of the triplicate chambers. The grey rectangle corresponds to the treatment window (days 5-10). Numbers above each plot refer to the ID of the donors. Data shown for Donors 24 and 81 are the same as in Fig. 1c.

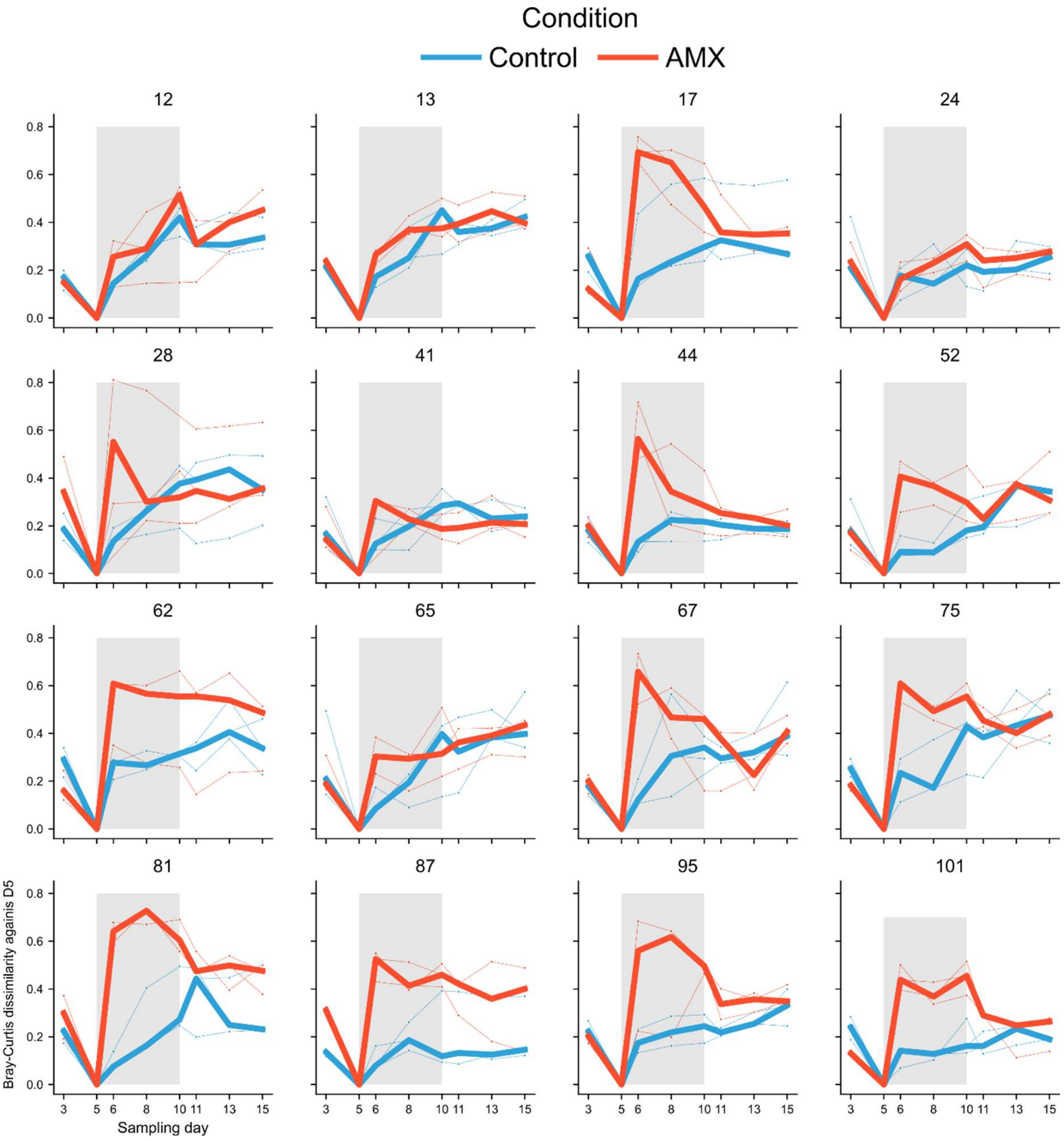


**Fig. S6**. Evolution of β-diversity in each MBRA chamber, given as their respective Bray-Curtis dissimilarity against their composition on Day 5. Thin lines represent each replicate chambers, with red lines representing AMX-treated chambers and blue lines representing Control chambers. Bold lines are median of the triplicate chambers. The grey rectangle corresponds to the treatment window (days 5-10). Numbers above each plot refer to the ID of the donors.

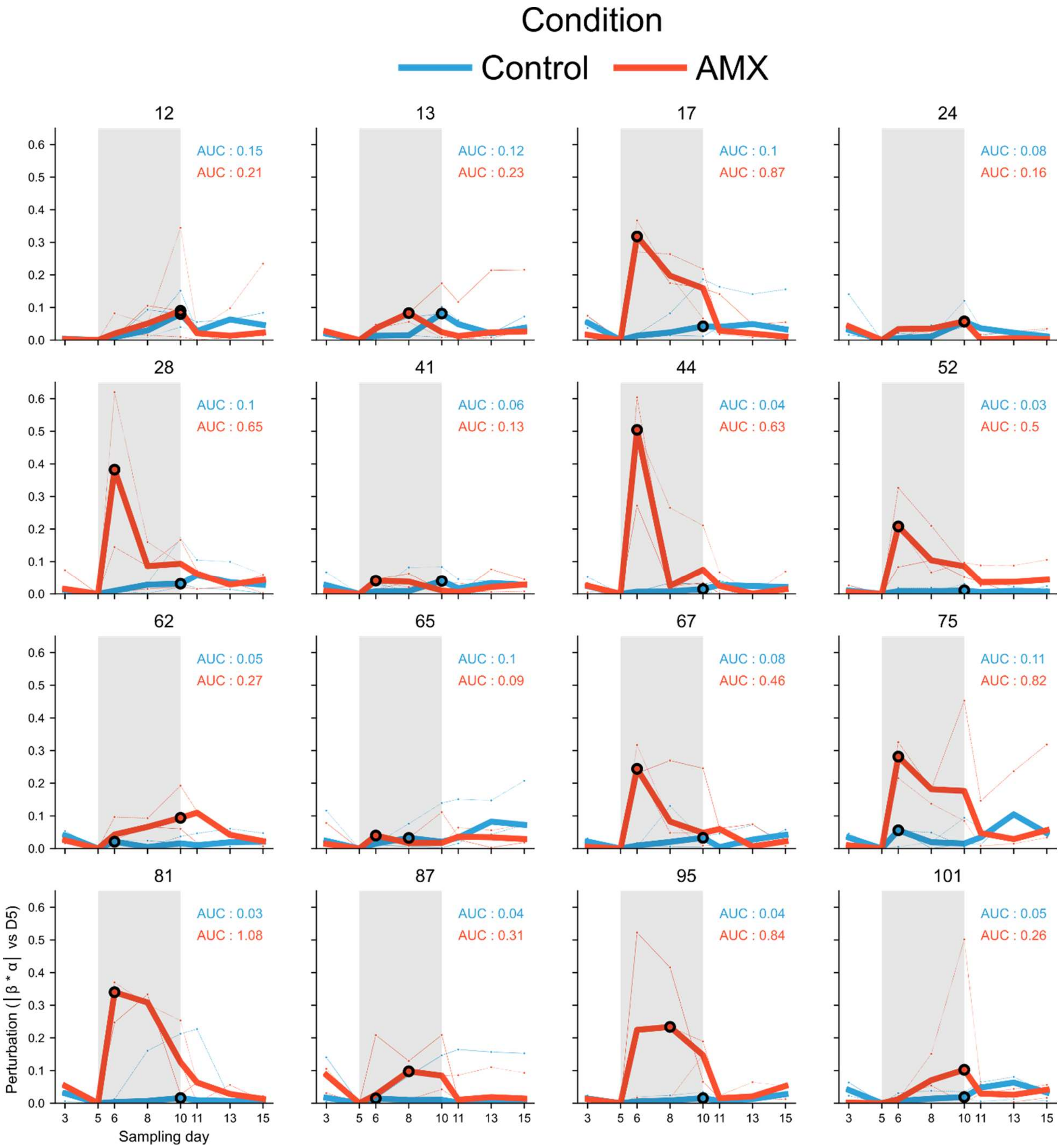


**Fig. S7**. Perturbation observed in each MBRA chamber, given as |β * α| metrics against respective Day 5 compositions and α-diversity. For β-diversity, Jensen-Shannon divergence was used, and for α-diversity, Shannon entropy. Thin lines represent each replicate chambers, with red lines representing AMX-treated chambers and blue lines representing Control chambers. Bold lines are median of the triplicate chambers. The grey rectangle corresponds to the treatment window (days 5-10). Numbers above each plot refer to the ID of the donors. Area Under Curve (AUC) of the median curves are indicated on each plot. Data shown for Donors 24 and 81 are the same as in Fig. 1d.

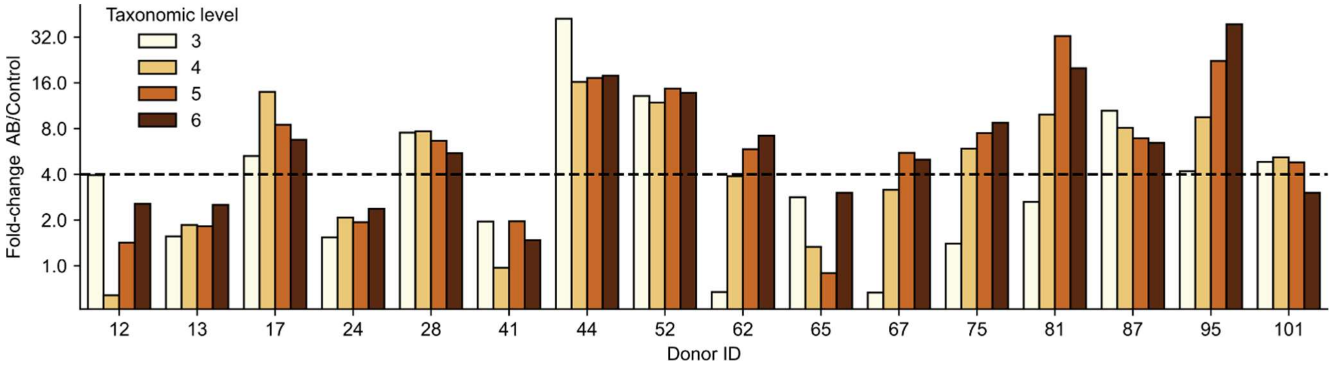


**Fig. S8**. Bar plot representing the strength of AMX perturbation across all the 16 donors of the panel. The values refer to the perturbation calculated when using data from various taxonomic levels (3: Class; 4: Order; 5: Family; 6: Genus). Values for the Family taxonomic level are the same as in Fig. 1f and used as reference. The bold dashed line corresponds to the Fold-change of 4 that was chosen to separate AMX-Sensitive donors from AMX-Robust donors.

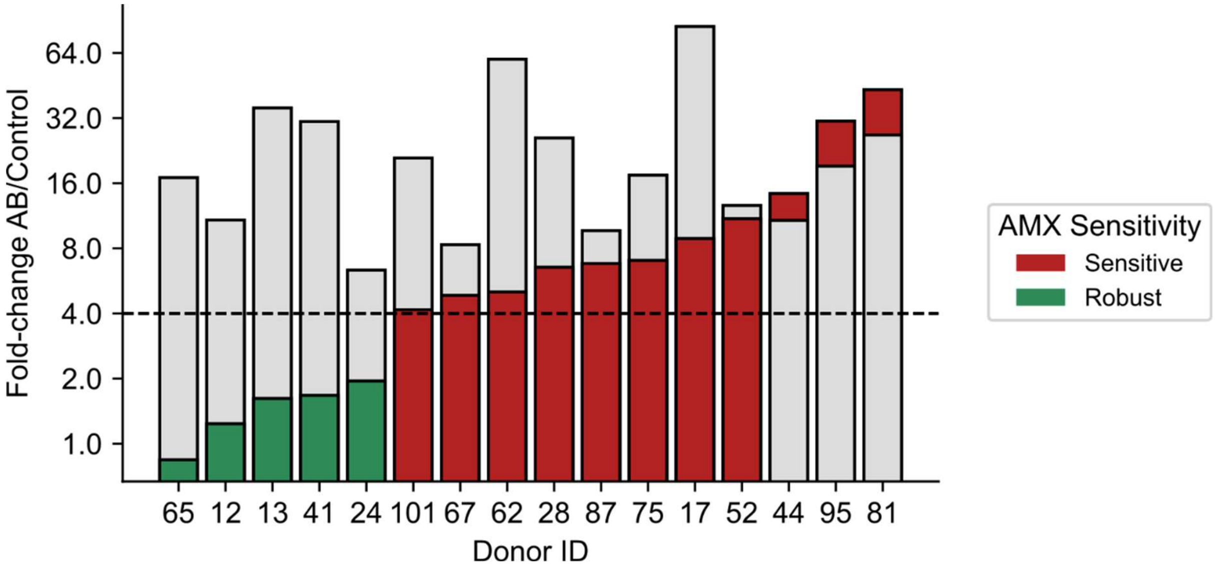


**Fig. S9**. Bar plot representing the strength of AMX and AMC perturbations across all the 16 donors of the panel. These values were obtained by repeating the analytical pipeline but using read rarefaction as a normalisation method. Donors are categorised between AMX-Sensitive or AMX-Robust groups depending on the value (Sensitive: FC>4).

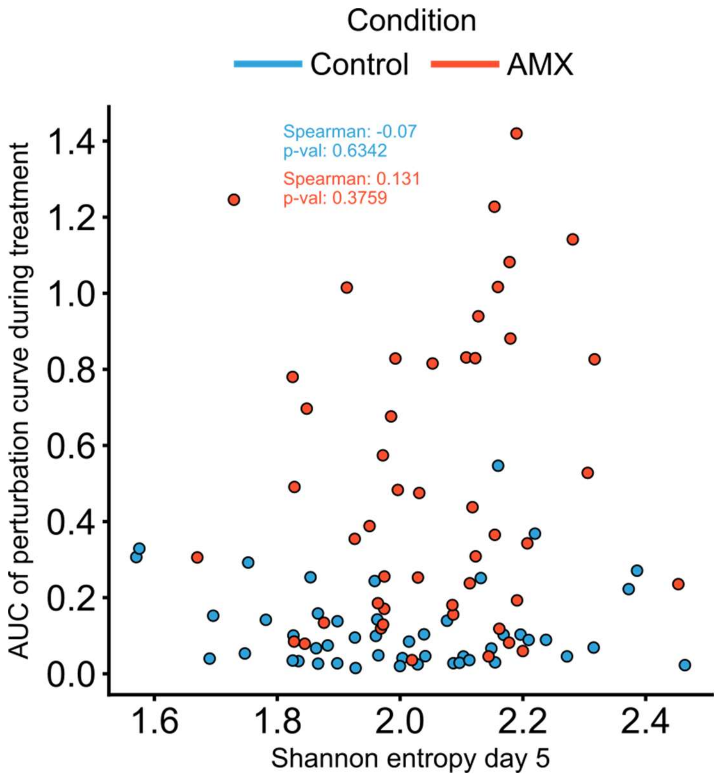


**Fig. S10**: Relationship between the Shannon entropy (α-diversity) of each chamber at Day 5 against its perturbation during treatment, measured as the AUC of its perturbation curve (see Fig. 1). AUC values are given for each replicate. Red dots correspond to chambers treated with AMX and blue dots to the Control chambers. Spearman correlation coefficients between the two plotted variables are given for either condition, with associated p-values.

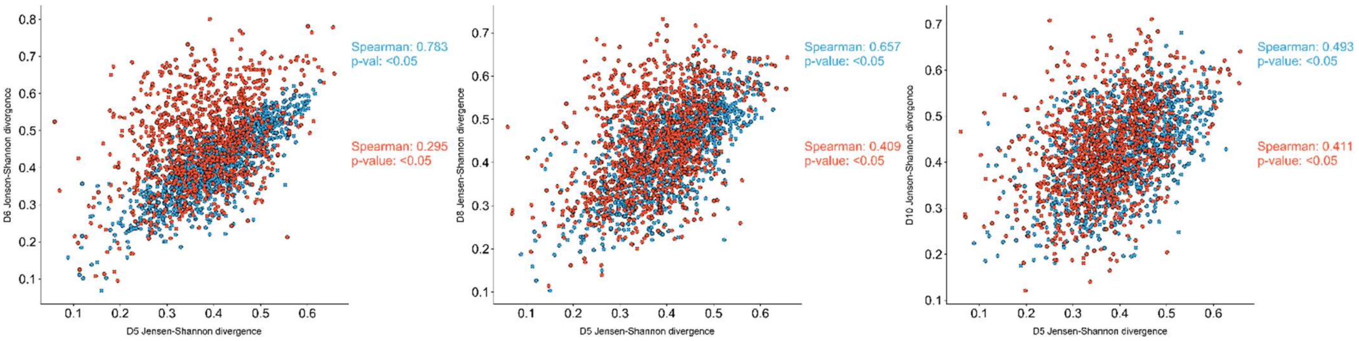


**Fig. S11**. Comparison of Jensen-Shannon divergence between pairs of MBRA chambers. Each dot represents a pair of MBRA chambers, with associated β-diversity at Day 5 or any of Days 6,8 or 10. Red dots represent pairs of AMX-treated chambers and blue dots represent pairs of Control chambers. Associated Spearman correlation coefficients and p-values are given in the same color scheme.

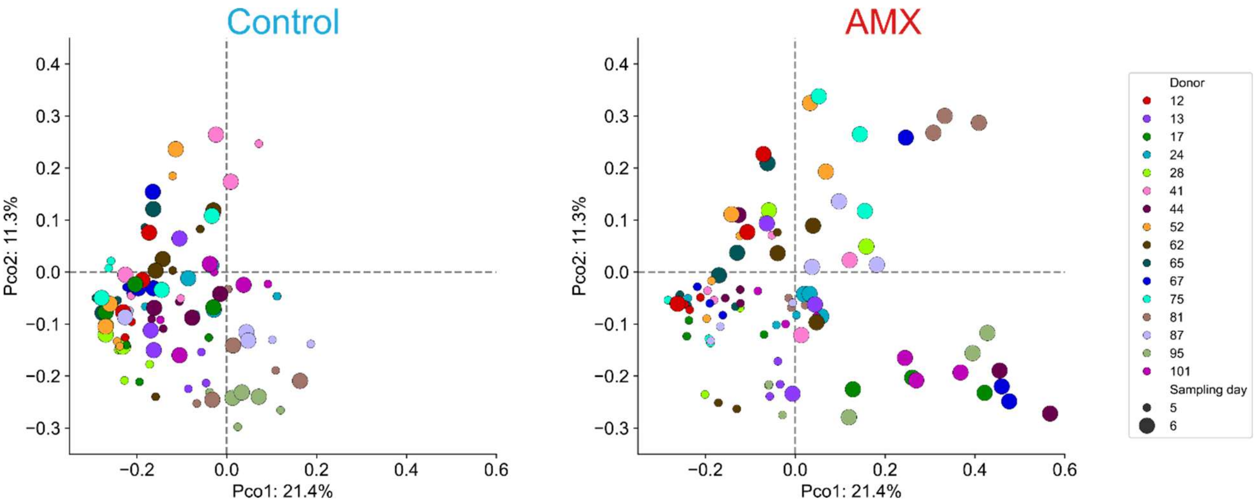


**Fig. S12**. Principal coordinate analysis of the various microbiotas of this study. All microbiotas of all chambers were analysed together using a Bray-Curtis dissimilarity matrix. On the left are shown microbiotas in control chambers at Day 5 and Day 6. On the right, microbiotas treated with AMX are shown at Day 5 and Day 6. Dot size corresponds to the displayed day. Dots are colored according to the Donor ID associated with the chamber.

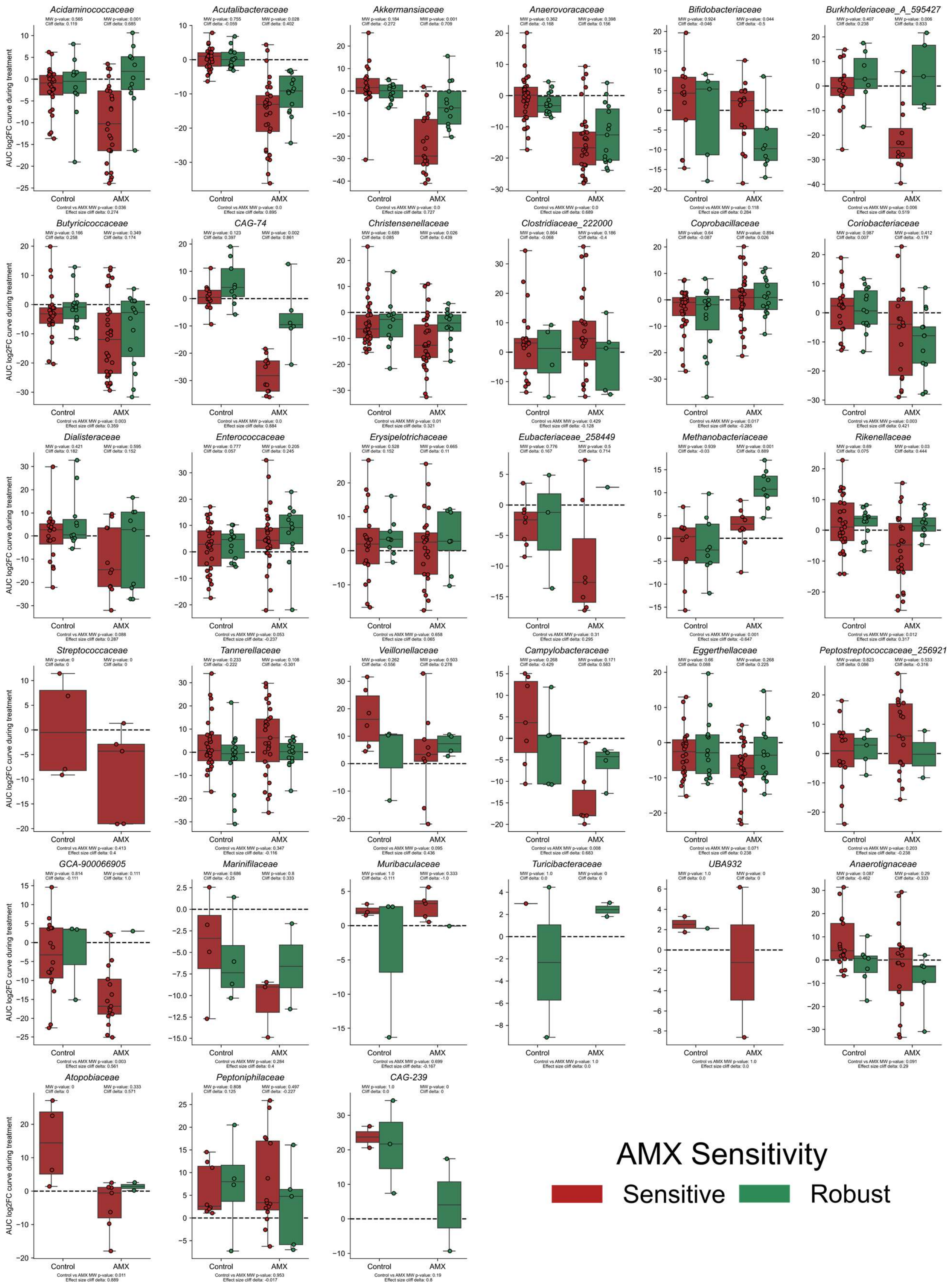
Acidaminococcaceae
Acutalibacteraceae
Akkermansiaceae
Anaerovoracaceae
Bifidobacteriaceae
Burkholderiaceae_A_595427
Butyricicoccaceae
CAG-74
Christensenellaceae
Clostridiaceae_222000
Coprobacillaceae
Coriobacteriaceae
Dialisteraceae
Enterococcaceae
Erysipelotrichaceae
Eubacteriaceae_258449
Methanobacteriaceae
Rikenellaceae
Streptococcaceae
Tannerellaceae
Veillonellaceae
Campylobacteraceae
Eggerthellaceae
Peptostreptococcaceae_256921
GCA-900066905
Marinifilaceae
Muribaculaceae
Turicibacteraceae
UBA932
Anaerotignaceae
Atopobiaceae
Peptoniphilaceae
CAG-239
AUC log2FC curve during treatment
Control
AMX
AMX Sensitivity
Sensitive
Robust

**Fig. S13**. AUC values of log2FC abundance change of all remaining families (not in Fig. 3b) in chambers either treated with AMX or untreated (Control condition). In green are chambers of AMX-Robust donors, and in red chambers of AMX-Sensitive donors. Mann-Whitney (MW) p-values and Cliff's delta effect size are provided, testing the differences between AMX-Sensitive chambers vs AMX-Robust chambers in both conditions, and between all chambers treated by AMX or untreated.

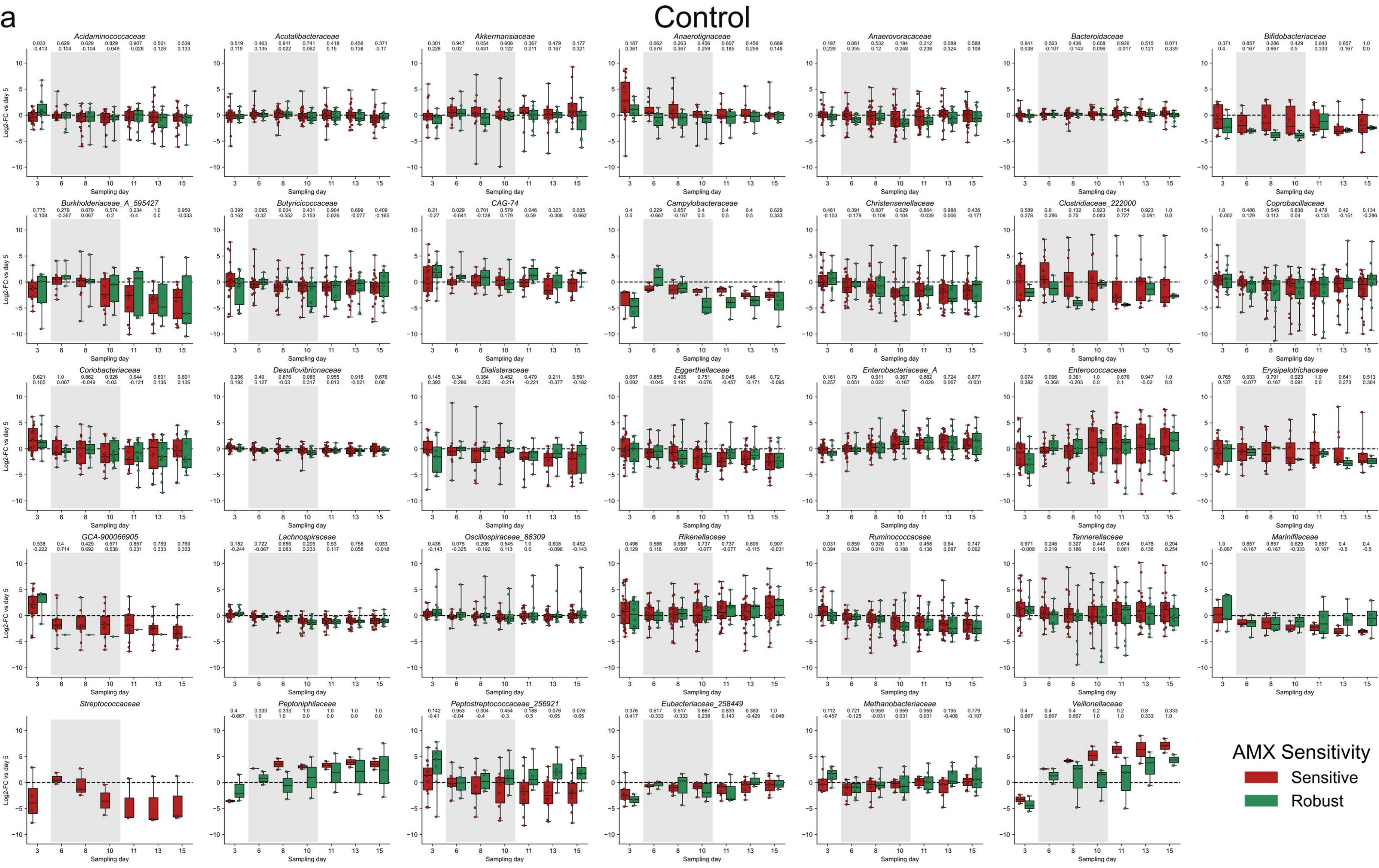
a
Control
Acidaminococcaceae
Acutalibacteraceae
Akkermansiaceae
Anaerotignaceae
Anaerovoracaceae
Bacteroidaceae
Bifidobacteriaceae
Burkholderiaceae_A_595427
Butyricicoccaceae
CAG-74
Campylobacteraceae
Christensenellaceae
Clostridiaceae_222000
Coprobacillaceae
Coriobacteriaceae
Desulfovibrionaceae
Dialisteraceae
Eggerthellaceae
Enterobacteriaceae_A
Enterococcaceae
Erysipelotrichaceae
GCA-900066905
Lachnospiraceae
Oscillospiraceae_88309
Rikenellaceae
Ruminococcaceae
Tannerellaceae
Marinifilaceae
Streptococcaceae
Peptoniphilaceae
Peptostreptococcaceae_256921
Eubacteriaceae_258449
Methanobacteriaceae
Veillonellaceae
Log2-FC vs day 5
Sampling day
AMX Sensitivity
Sensitive
Robust

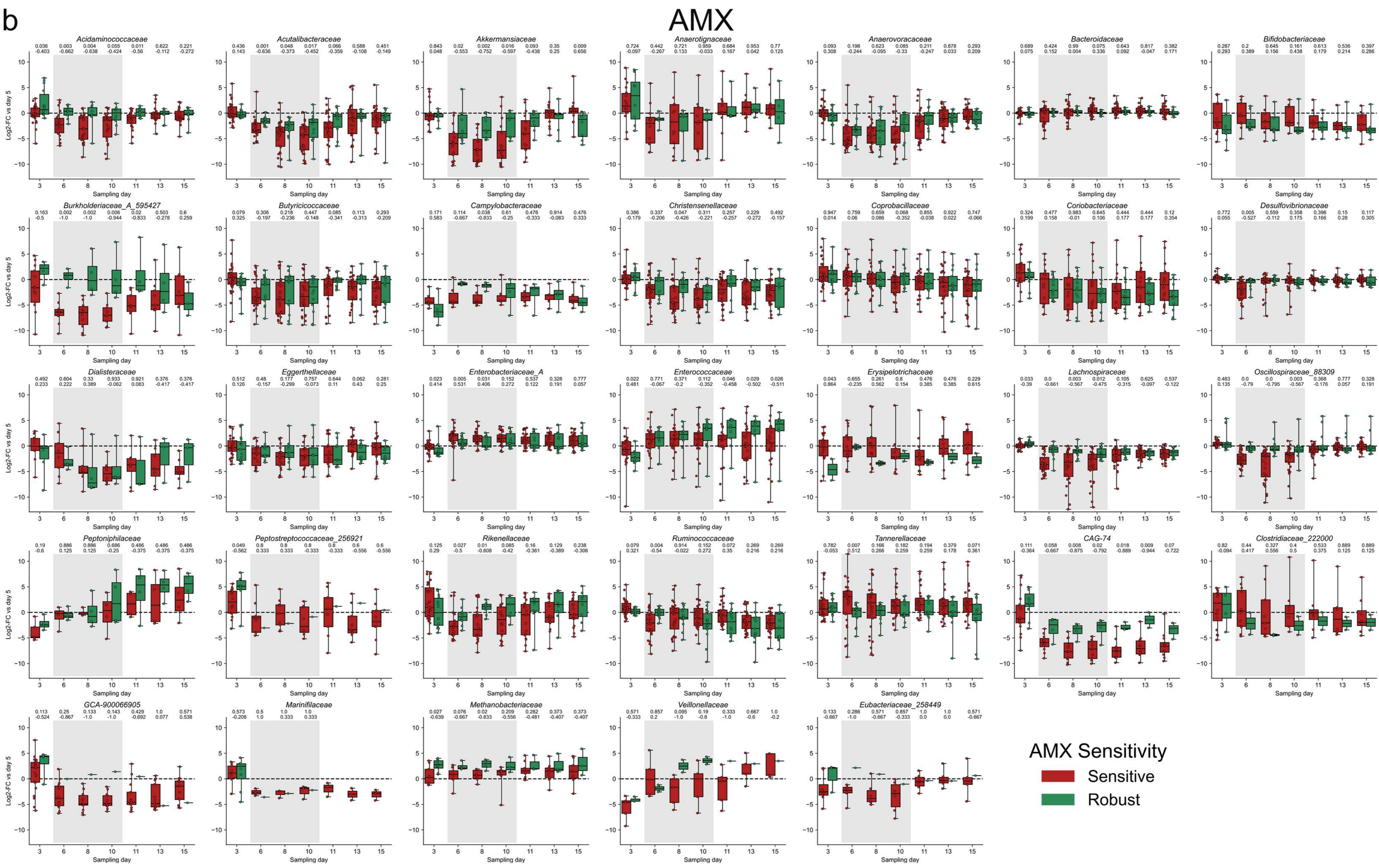
b
AMX
Acidaminococcaceae
Acutalibacteraceae
Akkermansiaceae
Anaerotignaceae
Anaerovoracaceae
Bacteroidaceae
Bifidobacteriaceae
Burkholderiaceae_A_595427
Butyricicoccaceae
Campylobacteraceae
Christensenellaceae
Coprobacillaceae
Coriobacteriaceae
Desulfovibrionaceae
Dialisteraceae
Eggerthellaceae
Enterobacteriaceae_A
Enterococcaceae
Erysipelotrichaceae
Lachnospiraceae
Oscillospiraceae_88309
Peptoniphilaceae
Peptostreptococcaceae_256921
Rikenellaceae
Ruminococcaceae
Tannerellaceae
CAG-74
Clostridiaceae_222000
GCA-900066905
Marinifilaceae
Methanobacteriaceae
Veillonellaceae
Eubacteriaceae_258449
Log2-FC vs day 5
Sampling day
AMX Sensitivity
Sensitive
Robust

**Fig S14**. Day-to-day log2FC abundance change of all families in chambers either untreated (control condition - a) or treated with AMX (b). In green are chambers of AMX-Robust donors, and in red chambers of AMX-Sensitive donors. Above each sampling day, Mann-Whitney p-values (top) and Cliff's delta effect sizes (bottom) are provided, testing the differences between AMX-Sensitive chambers vs AMX-Robust chambers at the given sampling day.

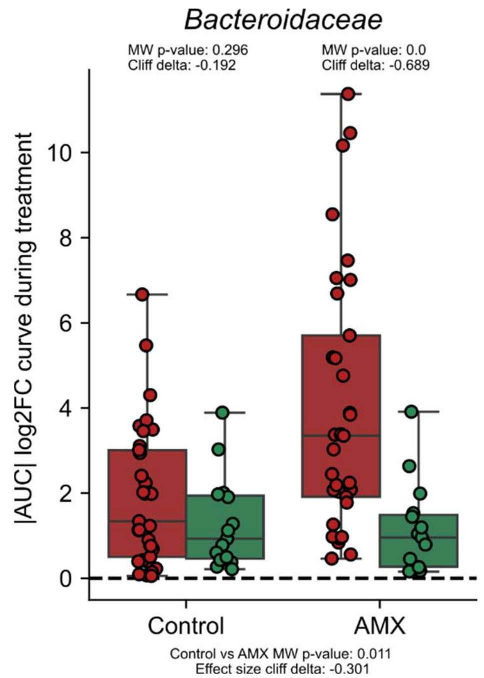


**Fig. S15**. Absolute AUC values of log2FC abundance change of the *Bacteroideaceae* family in chambers either treated with AMX or untreated (control condition). In green are chambers of AMX-Robust donors, and in red chambers of AMX-Sensitive donors. Mann-Whitney (MW) p-values and Cliff's delta effect size are provided, testing the differences between AMX-Sensitive chambers vs AMX-Robust chambers in both conditions, and between all chambers treated by AMX or untreated.

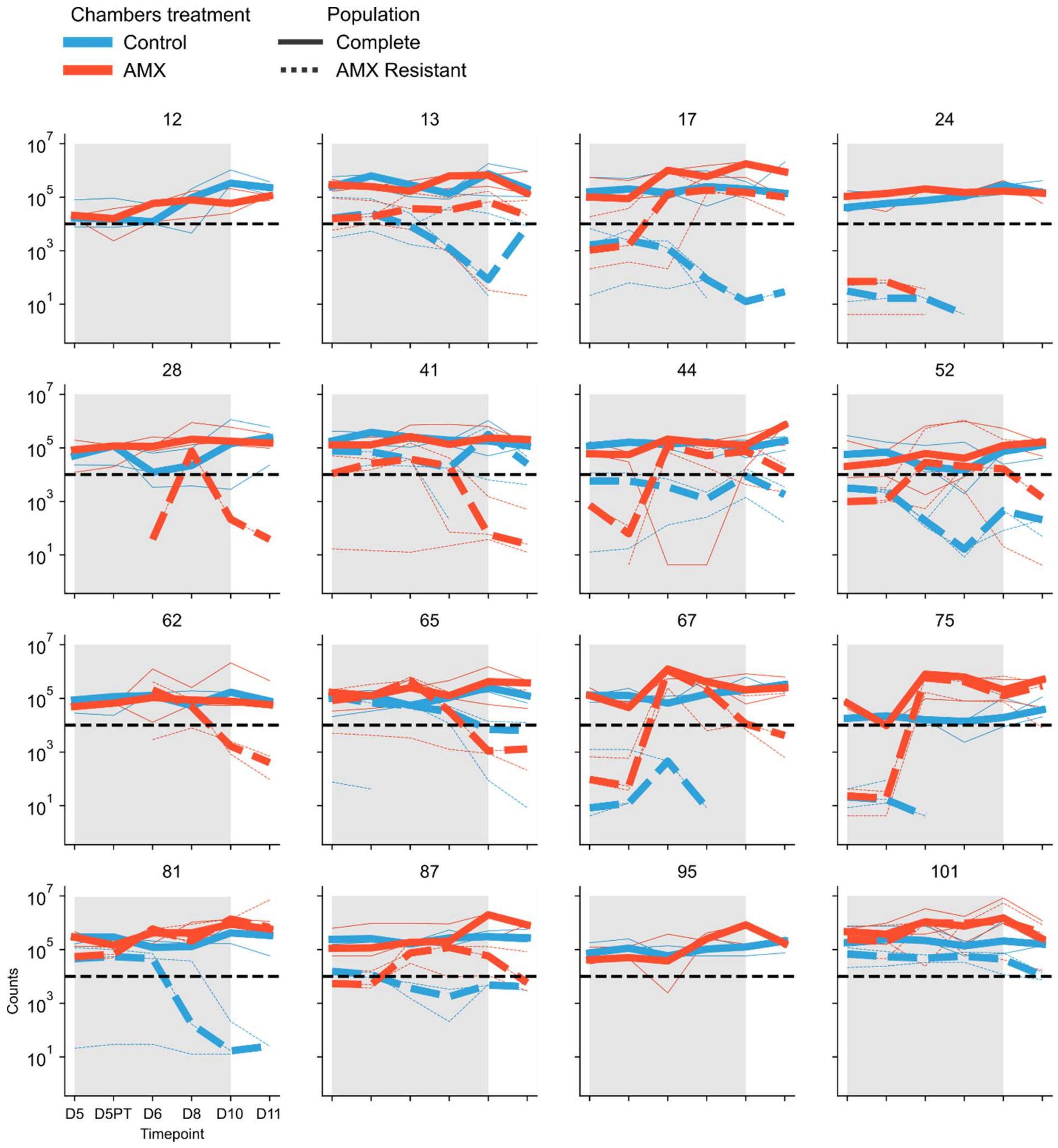


**Fig. S16**. Change in *Enterobacteriaceae* population counts over time in each MBRA chamber. For each donor, populations from Control (blue) or AMX-treated (red) chambers are shown. Since populations are plated on Drigalski agar with or without AMX, we could also count the total population (full lines) or AMX-resistant population (dashed lines) in each chamber. For each triplicate, replicates are shown as thin lines and the median is shown as a bold line. The detection limit of $10^4$ is shown as a dark bold dashed line.

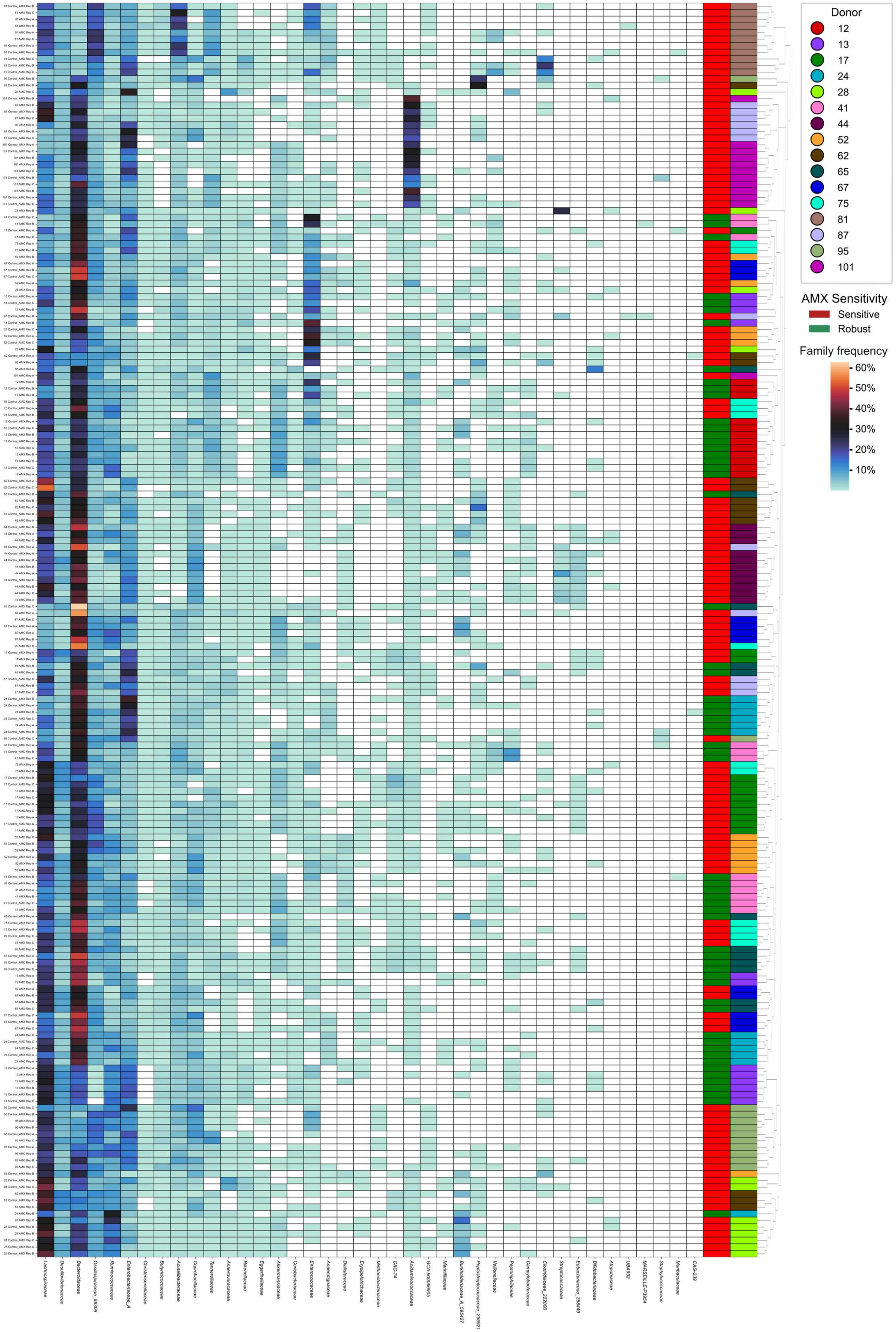

Donor
12
13
17
24
28
41
44
52
62
65
67
75
81
87
95
101
AMX Sensitivity
Sensitive
Robust
Family frequency
60%
50%
40%
30%
20%
10%
Lachnospiraceae
Desulfovibrionaceae
Bacteroidaceae
Oscillospiraceae_88309
Ruminococcaceae
Enterobacteriaceae_A
Christensenellaceae
Butyricicoccaceae
Acutalibacteraceae
Coprobacillaceae
Tannerellaceae
Anaerovoracaceae
Rikenellaceae
Eggerthellaceae
Akkermansiaceae
Coriobacteriaceae
Enterococcaceae
Anaerotignaceae
Dialisteraceae
Erysipelotrichaceae
Methanobacteriaceae
CAG-74
Acidaminococcaceae
GCA-900066905
Marinifilaceae
Burkholderiaceae_A_595427
Veillonellaceae
Peptoniphilaceae
Campylobacteraceae
Clostridiaceae_222000
Streptococcaceae
Eubacteriaceae_256449
Bifidobacteriaceae
Atopobiaceae
UBA932
MARSEILLE-P3954
Staphylococcaceae
Muribaculaceae
CAG-239

**Fig. S17.** Clustermap representing the microbiota composition of each chamber at Day 5, including both the AMX and AMC experiments (12 replicates per donor in total). Hierarchical clustering of the microbiotas was performed using a Jensen-Shannon divergence matrix. Donor IDs are indicated in colors above the heatmap. Donor AMX sensitivity is also indicated in colors above the heatmap, using the classification of Fig. 1. Heatmap cells are colored according to the abundance of the indicated bacterial family (in % of the total microbiota).

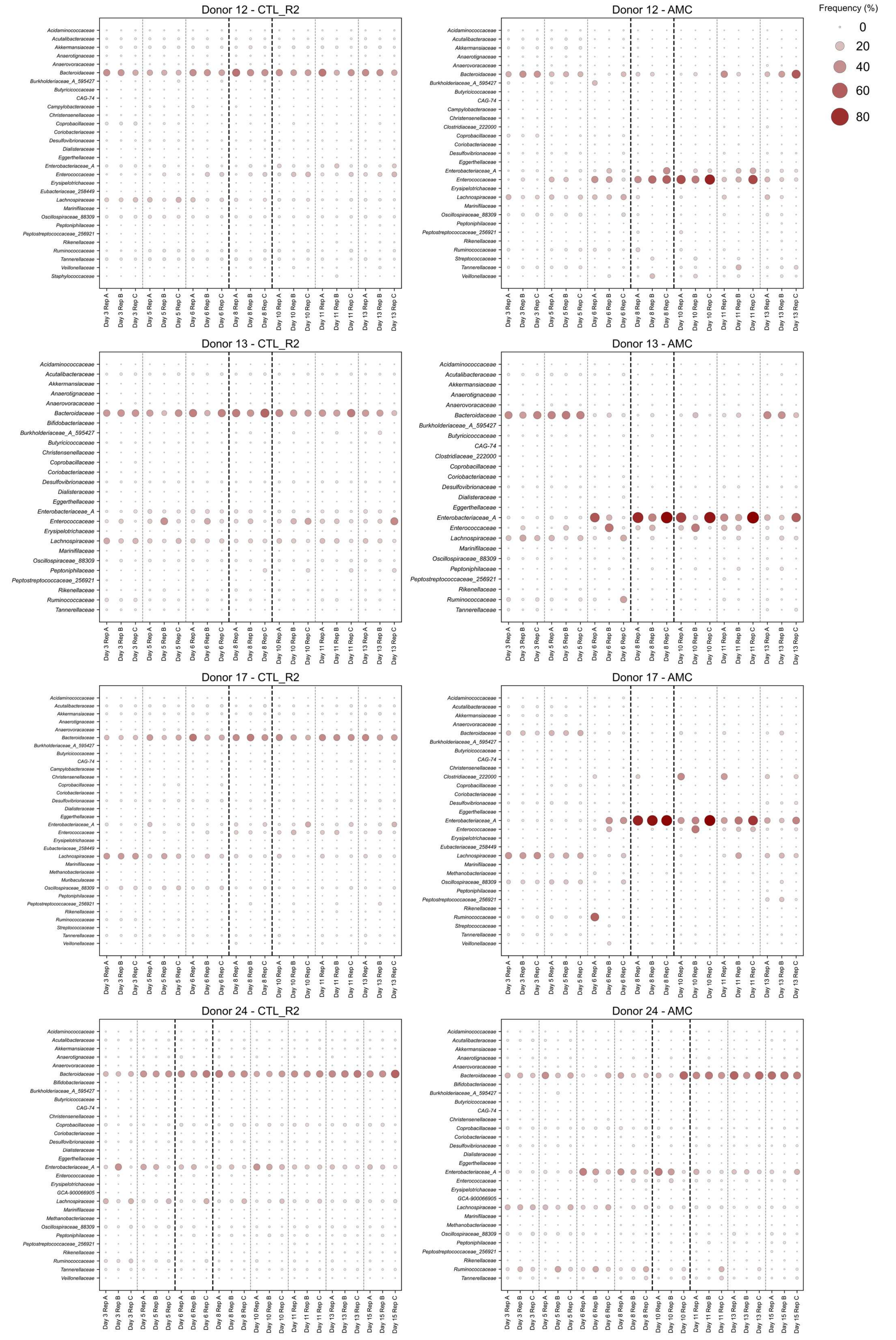
Donor 12 - CTL_R2
Donor 12 - AMC
Donor 13 - CTL_R2
Donor 13 - AMC
Donor 17 - CTL_R2
Donor 17 - AMC
Donor 24 - CTL_R2
Donor 24 - AMC
Frequency (%)
0
20
40
60
80
Acidaminococcaceae
Acutalibacteraceae
Akkermansiaceae
Anaerotignaceae
Anaerovoracaceae
Bacteroidaceae
Bifidobacteriaceae
Burkholderiaceae_A_595427
Butyricicoccaceae
CAG-74
Campylobacteraceae
Christensenellaceae
Clostridiaceae_222000
Coprobacillaceae
Coriobacteriaceae
Desulfovibrionaceae
Dialisteraceae
Eggerthellaceae
Enterobacteriaceae_A
Enterococcaceae
Erysipelotrichaceae
Eubacteriaceae_258449
GCA-900066905
Lachnospiraceae
Marinifilaceae
Methanobacteriaceae
Muribaculaceae
Oscillospiraceae_88309
Peptoniphilaceae
Peptostreptococcaceae_256921
Rikenellaceae
Ruminococcaceae
Streptococcaceae
Tannerellaceae
Veillonellaceae
Staphylococcaceae
Day 3 Rep A
Day 3 Rep B
Day 3 Rep C
Day 5 Rep A
Day 5 Rep B
Day 5 Rep C
Day 6 Rep A
Day 6 Rep B
Day 6 Rep C
Day 8 Rep A
Day 8 Rep B
Day 8 Rep C
Day 10 Rep A
Day 10 Rep B
Day 10 Rep C
Day 11 Rep A
Day 11 Rep B
Day 11 Rep C
Day 13 Rep A
Day 13 Rep B
Day 13 Rep C
Day 15 Rep A
Day 15 Rep B
Day 15 Rep C

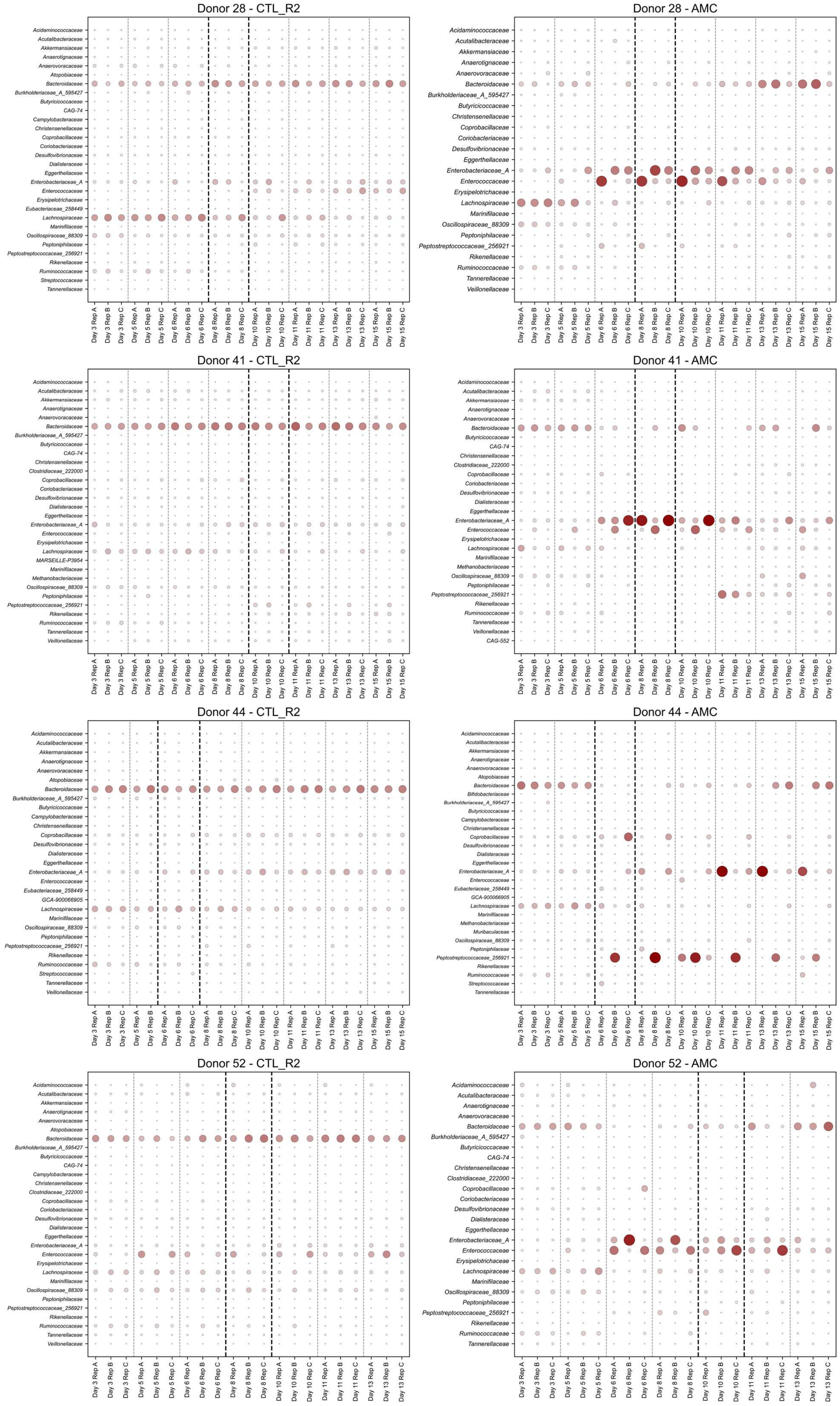

Donor 28 - CTL_R2
Acidaminococcaceae
Acutalibacteraceae
Akkermansiaceae
Anaerotignaceae
Anaerovoracaceae
Atopobiaceae
Bacteroidaceae
Burkholderiaceae_A_595427
Butyricicoccaceae
CAG-74
Campylobacteraceae
Christensenellaceae
Coprobacillaceae
Coriobacteriaceae
Desulfovibrionaceae
Dialisteraceae
Eggerthellaceae
Enterobacteriaceae_A
Enterococcaceae
Erysipelotrichaceae
Eubacteriaceae_258449
Lachnospiraceae
Marinifilaceae
Oscillospiraceae_88309
Peptoniphilaceae
Peptostreptococcaceae_256921
Rikenellaceae
Ruminococcaceae
Streptococcaceae
Tannerellaceae
Day 3 Rep A
Day 3 Rep B
Day 3 Rep C
Day 5 Rep A
Day 5 Rep B
Day 5 Rep C
Day 6 Rep A
Day 6 Rep B
Day 6 Rep C
Day 8 Rep A
Day 8 Rep B
Day 8 Rep C
Day 10 Rep A
Day 10 Rep B
Day 10 Rep C
Day 11 Rep A
Day 11 Rep B
Day 11 Rep C
Day 13 Rep A
Day 13 Rep B
Day 13 Rep C
Day 15 Rep A
Day 15 Rep B
Day 15 Rep C
Donor 28 - AMC
Acidaminococcaceae
Acutalibacteraceae
Akkermansiaceae
Anaerotignaceae
Anaerovoracaceae
Bacteroidaceae
Burkholderiaceae_A_595427
Butyricicoccaceae
Christensenellaceae
Coprobacillaceae
Coriobacteriaceae
Desulfovibrionaceae
Eggerthellaceae
Enterobacteriaceae_A
Enterococcaceae
Erysipelotrichaceae
Lachnospiraceae
Marinifilaceae
Oscillospiraceae_88309
Peptoniphilaceae
Peptostreptococcaceae_256921
Rikenellaceae
Ruminococcaceae
Tannerellaceae
Veillonellaceae
Donor 41 - CTL_R2
Acidaminococcaceae
Acutalibacteraceae
Akkermansiaceae
Anaerotignaceae
Anaerovoracaceae
Bacteroidaceae
Burkholderiaceae_A_595427
Butyricicoccaceae
CAG-74
Christensenellaceae
Clostridiaceae_222000
Coprobacillaceae
Coriobacteriaceae
Desulfovibrionaceae
Dialisteraceae
Eggerthellaceae
Enterobacteriaceae_A
Enterococcaceae
Erysipelotrichaceae
Lachnospiraceae
MARSEILLE-P3954
Marinifilaceae
Methanobacteriaceae
Oscillospiraceae_88309
Peptoniphilaceae
Peptostreptococcaceae_256921
Rikenellaceae
Ruminococcaceae
Tannerellaceae
Veillonellaceae
Donor 41 - AMC
Acidaminococcaceae
Acutalibacteraceae
Akkermansiaceae
Anaerotignaceae
Anaerovoracaceae
Bacteroidaceae
Butyricicoccaceae
CAG-74
Christensenellaceae
Clostridiaceae_222000
Coprobacillaceae
Coriobacteriaceae
Desulfovibrionaceae
Dialisteraceae
Eggerthellaceae
Enterobacteriaceae_A
Enterococcaceae
Erysipelotrichaceae
Lachnospiraceae
Marinifilaceae
Methanobacteriaceae
Oscillospiraceae_88309
Peptoniphilaceae
Peptostreptococcaceae_256921
Rikenellaceae
Ruminococcaceae
Tannerellaceae
Veillonellaceae
CAG-552
Donor 44 - CTL_R2
Acidaminococcaceae
Acutalibacteraceae
Akkermansiaceae
Anaerotignaceae
Anaerovoracaceae
Atopobiaceae
Bacteroidaceae
Burkholderiaceae_A_595427
Butyricicoccaceae
Campylobacteraceae
Christensenellaceae
Coprobacillaceae
Desulfovibrionaceae
Dialisteraceae
Eggerthellaceae
Enterobacteriaceae_A
Enterococcaceae
Eubacteriaceae_258449
GCA-900066905
Lachnospiraceae
Marinifilaceae
Oscillospiraceae_88309
Peptoniphilaceae
Peptostreptococcaceae_256921
Rikenellaceae
Ruminococcaceae
Streptococcaceae
Tannerellaceae
Veillonellaceae
Donor 44 - AMC
Acidaminococcaceae
Acutalibacteraceae
Akkermansiaceae
Anaerotignaceae
Anaerovoracaceae
Atopobiaceae
Bacteroidaceae
Bifidobacteriaceae
Burkholderiaceae_A_595427
Butyricicoccaceae
Campylobacteraceae
Christensenellaceae
Coprobacillaceae
Desulfovibrionaceae
Dialisteraceae
Eggerthellaceae
Enterobacteriaceae_A
Enterococcaceae
Eubacteriaceae_258449
GCA-900066905
Lachnospiraceae
Marinifilaceae
Methanobacteriaceae
Muribaculaceae
Oscillospiraceae_88309
Peptoniphilaceae
Peptostreptococcaceae_256921
Rikenellaceae
Ruminococcaceae
Streptococcaceae
Tannerellaceae
Donor 52 - CTL_R2
Acidaminococcaceae
Acutalibacteraceae
Akkermansiaceae
Anaerotignaceae
Anaerovoracaceae
Atopobiaceae
Bacteroidaceae
Burkholderiaceae_A_595427
Butyricicoccaceae
CAG-74
Campylobacteraceae
Christensenellaceae
Clostridiaceae_222000
Coprobacillaceae
Coriobacteriaceae
Desulfovibrionaceae
Dialisteraceae
Eggerthellaceae
Enterobacteriaceae_A
Enterococcaceae
Erysipelotrichaceae
Lachnospiraceae
Marinifilaceae
Oscillospiraceae_88309
Peptoniphilaceae
Peptostreptococcaceae_256921
Rikenellaceae
Ruminococcaceae
Tannerellaceae
Veillonellaceae
Day 3 Rep A
Day 3 Rep B
Day 3 Rep C
Day 5 Rep A
Day 5 Rep B
Day 5 Rep C
Day 6 Rep A
Day 6 Rep B
Day 6 Rep C
Day 8 Rep A
Day 8 Rep B
Day 8 Rep C
Day 10 Rep A
Day 10 Rep B
Day 10 Rep C
Day 11 Rep A
Day 11 Rep B
Day 11 Rep C
Day 13 Rep A
Day 13 Rep B
Day 13 Rep C
Donor 52 - AMC
Acidaminococcaceae
Acutalibacteraceae
Anaerotignaceae
Anaerovoracaceae
Bacteroidaceae
Burkholderiaceae_A_595427
Butyricicoccaceae
CAG-74
Christensenellaceae
Clostridiaceae_222000
Coprobacillaceae
Coriobacteriaceae
Desulfovibrionaceae
Dialisteraceae
Eggerthellaceae
Enterobacteriaceae_A
Enterococcaceae
Erysipelotrichaceae
Lachnospiraceae
Marinifilaceae
Oscillospiraceae_88309
Peptoniphilaceae
Peptostreptococcaceae_256921
Rikenellaceae
Ruminococcaceae
Tannerellaceae

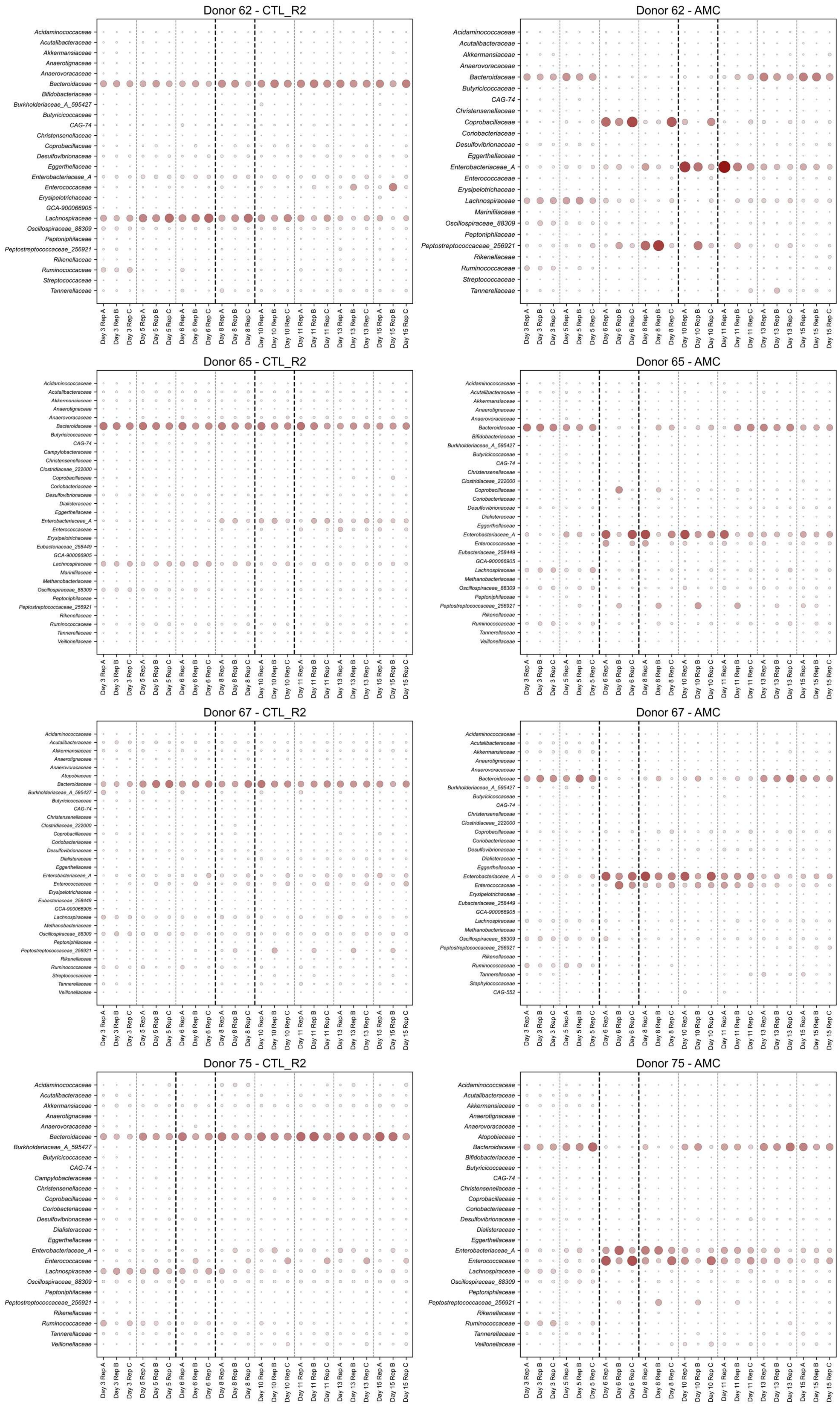

Donor 62 - CTL_R2
Acidaminococcaceae
Acutalibacteraceae
Akkermansiaceae
Anaerotignaceae
Anaerovoracaceae
Bacteroidaceae
Bifidobacteriaceae
Burkholderiaceae_A_595427
Butyricicoccaceae
CAG-74
Christensenellaceae
Coprobacillaceae
Desulfovibrionaceae
Eggerthellaceae
Enterobacteriaceae_A
Enterococcaceae
Erysipelotrichaceae
GCA-900066905
Lachnospiraceae
Oscillospiraceae_88309
Peptoniphilaceae
Peptostreptococcaceae_256921
Rikenellaceae
Ruminococcaceae
Streptococcaceae
Tannerellaceae
Day 3 Rep A
Day 3 Rep B
Day 3 Rep C
Day 5 Rep A
Day 5 Rep B
Day 5 Rep C
Day 6 Rep A
Day 6 Rep B
Day 6 Rep C
Day 8 Rep A
Day 8 Rep B
Day 8 Rep C
Day 10 Rep A
Day 10 Rep B
Day 10 Rep C
Day 11 Rep A
Day 11 Rep B
Day 11 Rep C
Day 13 Rep A
Day 13 Rep B
Day 13 Rep C
Day 15 Rep A
Day 15 Rep B
Day 15 Rep C
Donor 62 - AMC
Acidaminococcaceae
Acutalibacteraceae
Akkermansiaceae
Anaerovoracaceae
Bacteroidaceae
Butyricicoccaceae
CAG-74
Christensenellaceae
Coprobacillaceae
Coriobacteriaceae
Desulfovibrionaceae
Eggerthellaceae
Enterobacteriaceae_A
Enterococcaceae
Erysipelotrichaceae
Lachnospiraceae
Marinifilaceae
Oscillospiraceae_88309
Peptoniphilaceae
Peptostreptococcaceae_256921
Rikenellaceae
Ruminococcaceae
Streptococcaceae
Tannerellaceae
Donor 65 - CTL_R2
Acidaminococcaceae
Acutalibacteraceae
Akkermansiaceae
Anaerotignaceae
Anaerovoracaceae
Bacteroidaceae
Butyricicoccaceae
CAG-74
Campylobacteraceae
Christensenellaceae
Clostridiaceae_222000
Coprobacillaceae
Coriobacteriaceae
Desulfovibrionaceae
Dialisteraceae
Eggerthellaceae
Enterobacteriaceae_A
Enterococcaceae
Erysipelotrichaceae
Eubacteriaceae_258449
GCA-900066905
Lachnospiraceae
Marinifilaceae
Methanobacteriaceae
Oscillospiraceae_88309
Peptoniphilaceae
Peptostreptococcaceae_256921
Rikenellaceae
Ruminococcaceae
Tannerellaceae
Veillonellaceae
Donor 65 - AMC
Acidaminococcaceae
Acutalibacteraceae
Akkermansiaceae
Anaerotignaceae
Anaerovoracaceae
Bacteroidaceae
Bifidobacteriaceae
Burkholderiaceae_A_595427
Butyricicoccaceae
CAG-74
Christensenellaceae
Clostridiaceae_222000
Coprobacillaceae
Coriobacteriaceae
Desulfovibrionaceae
Dialisteraceae
Eggerthellaceae
Enterobacteriaceae_A
Enterococcaceae
Eubacteriaceae_258449
GCA-900066905
Lachnospiraceae
Methanobacteriaceae
Oscillospiraceae_88309
Peptoniphilaceae
Peptostreptococcaceae_256921
Rikenellaceae
Ruminococcaceae
Tannerellaceae
Veillonellaceae
Donor 67 - CTL_R2
Acidaminococcaceae
Acutalibacteraceae
Akkermansiaceae
Anaerotignaceae
Anaerovoracaceae
Atopobiaceae
Bacteroidaceae
Burkholderiaceae_A_595427
Butyricicoccaceae
CAG-74
Christensenellaceae
Clostridiaceae_222000
Coprobacillaceae
Coriobacteriaceae
Desulfovibrionaceae
Dialisteraceae
Eggerthellaceae
Enterobacteriaceae_A
Enterococcaceae
Erysipelotrichaceae
Eubacteriaceae_258449
GCA-900066905
Lachnospiraceae
Methanobacteriaceae
Oscillospiraceae_88309
Peptoniphilaceae
Peptostreptococcaceae_256921
Rikenellaceae
Ruminococcaceae
Streptococcaceae
Tannerellaceae
Veillonellaceae
Donor 67 - AMC
Acidaminococcaceae
Acutalibacteraceae
Akkermansiaceae
Anaerotignaceae
Anaerovoracaceae
Bacteroidaceae
Burkholderiaceae_A_595427
Butyricicoccaceae
CAG-74
Christensenellaceae
Clostridiaceae_222000
Coprobacillaceae
Coriobacteriaceae
Desulfovibrionaceae
Dialisteraceae
Eggerthellaceae
Enterobacteriaceae_A
Enterococcaceae
Erysipelotrichaceae
Eubacteriaceae_258449
GCA-900066905
Lachnospiraceae
Methanobacteriaceae
Oscillospiraceae_88309
Peptostreptococcaceae_256921
Rikenellaceae
Ruminococcaceae
Tannerellaceae
Staphylococcaceae
CAG-552
Donor 75 - CTL_R2
Acidaminococcaceae
Acutalibacteraceae
Akkermansiaceae
Anaerotignaceae
Anaerovoracaceae
Bacteroidaceae
Burkholderiaceae_A_595427
Butyricicoccaceae
CAG-74
Campylobacteraceae
Christensenellaceae
Coprobacillaceae
Coriobacteriaceae
Desulfovibrionaceae
Dialisteraceae
Eggerthellaceae
Enterobacteriaceae_A
Enterococcaceae
Lachnospiraceae
Oscillospiraceae_88309
Peptoniphilaceae
Peptostreptococcaceae_256921
Rikenellaceae
Ruminococcaceae
Tannerellaceae
Veillonellaceae
Donor 75 - AMC
Acidaminococcaceae
Acutalibacteraceae
Akkermansiaceae
Anaerotignaceae
Anaerovoracaceae
Atopobiaceae
Bacteroidaceae
Bifidobacteriaceae
Butyricicoccaceae
CAG-74
Christensenellaceae
Coprobacillaceae
Coriobacteriaceae
Desulfovibrionaceae
Dialisteraceae
Eggerthellaceae
Enterobacteriaceae_A
Enterococcaceae
Lachnospiraceae
Oscillospiraceae_88309
Peptoniphilaceae
Peptostreptococcaceae_256921
Rikenellaceae
Ruminococcaceae
Tannerellaceae
Veillonellaceae

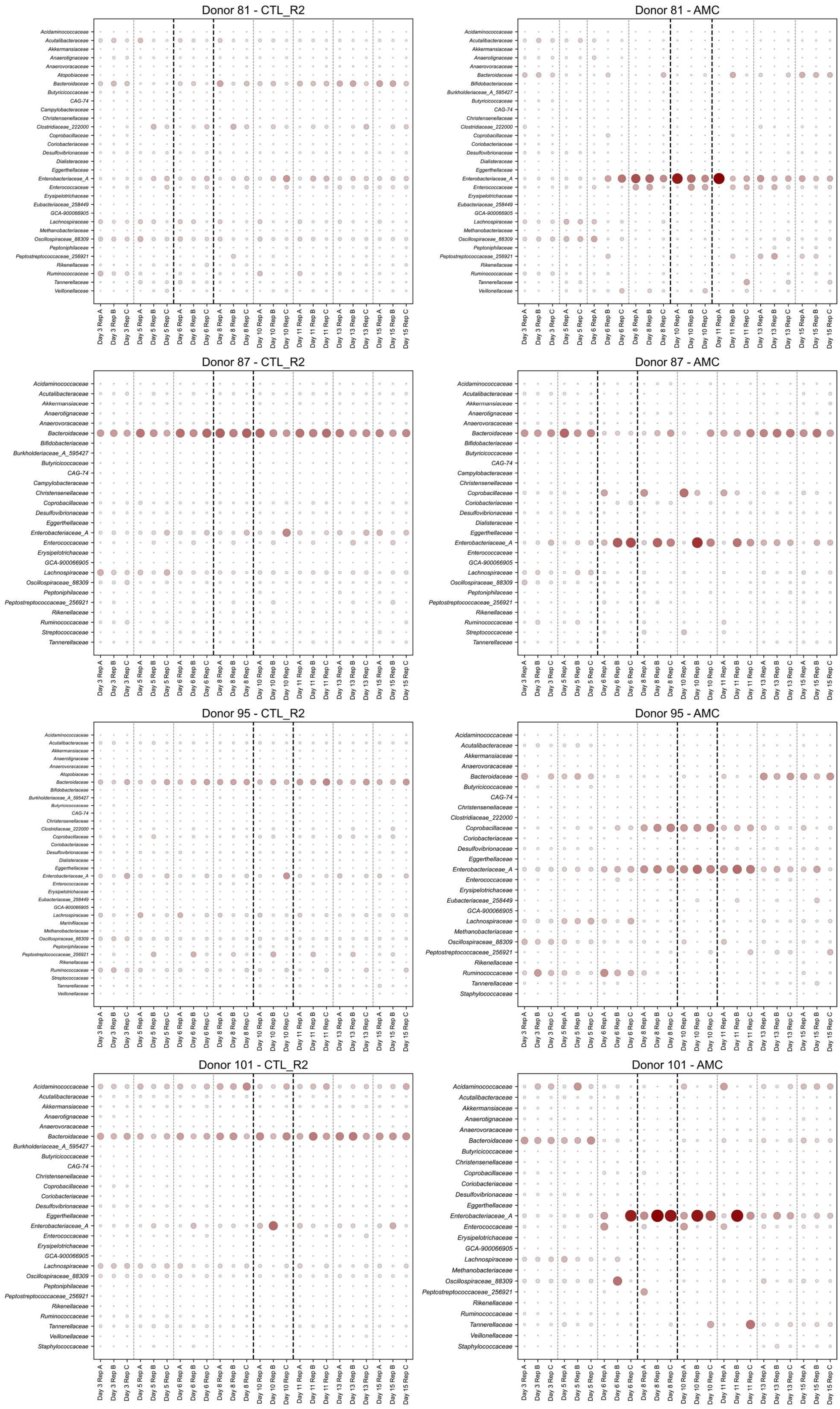
Donor 81 - CTL_R2
Acidaminococcaceae
Acutalibacteraceae
Akkermansiaceae
Anaerotignaceae
Anaerovoracaceae
Atopobiaceae
Bacteroidaceae
Butyricicoccaceae
CAG-74
Campylobacteraceae
Christensenellaceae
Clostridiaceae_222000
Coprobacillaceae
Coriobacteriaceae
Desulfovibrionaceae
Dialisteraceae
Eggerthellaceae
Enterobacteriaceae_A
Enterococcaceae
Erysipelotrichaceae
Eubacteriaceae_258449
GCA-900066905
Lachnospiraceae
Methanobacteriaceae
Oscillospiraceae_88309
Peptoniphilaceae
Peptostreptococcaceae_256921
Rikenellaceae
Ruminococcaceae
Tannerellaceae
Veillonellaceae
Day 3 Rep A
Day 3 Rep B
Day 3 Rep C
Day 5 Rep A
Day 5 Rep B
Day 5 Rep C
Day 6 Rep A
Day 6 Rep B
Day 6 Rep C
Day 8 Rep A
Day 8 Rep B
Day 8 Rep C
Day 10 Rep A
Day 10 Rep B
Day 10 Rep C
Day 11 Rep A
Day 11 Rep B
Day 11 Rep C
Day 13 Rep A
Day 13 Rep B
Day 13 Rep C
Day 15 Rep A
Day 15 Rep B
Day 15 Rep C
Donor 81 - AMC
Acidaminococcaceae
Acutalibacteraceae
Akkermansiaceae
Anaerotignaceae
Anaerovoracaceae
Bacteroidaceae
Bifidobacteriaceae
Burkholderiaceae_A_595427
Butyricicoccaceae
CAG-74
Christensenellaceae
Clostridiaceae_222000
Coprobacillaceae
Coriobacteriaceae
Desulfovibrionaceae
Dialisteraceae
Eggerthellaceae
Enterobacteriaceae_A
Enterococcaceae
Erysipelotrichaceae
Eubacteriaceae_258449
GCA-900066905
Lachnospiraceae
Methanobacteriaceae
Oscillospiraceae_88309
Peptoniphilaceae
Peptostreptococcaceae_256921
Rikenellaceae
Ruminococcaceae
Tannerellaceae
Veillonellaceae
Day 3 Rep A
Day 3 Rep B
Day 3 Rep C
Day 5 Rep A
Day 5 Rep B
Day 5 Rep C
Day 6 Rep A
Day 6 Rep B
Day 6 Rep C
Day 8 Rep A
Day 8 Rep B
Day 8 Rep C
Day 10 Rep A
Day 10 Rep B
Day 10 Rep C
Day 11 Rep A
Day 11 Rep B
Day 11 Rep C
Day 13 Rep A
Day 13 Rep B
Day 13 Rep C
Day 15 Rep A
Day 15 Rep B
Day 15 Rep C
Donor 87 - CTL_R2
Acidaminococcaceae
Acutalibacteraceae
Akkermansiaceae
Anaerotignaceae
Anaerovoracaceae
Bacteroidaceae
Bifidobacteriaceae
Burkholderiaceae_A_595427
Butyricicoccaceae
CAG-74
Campylobacteraceae
Christensenellaceae
Coprobacillaceae
Desulfovibrionaceae
Eggerthellaceae
Enterobacteriaceae_A
Enterococcaceae
Erysipelotrichaceae
GCA-900066905
Lachnospiraceae
Oscillospiraceae_88309
Peptoniphilaceae
Peptostreptococcaceae_256921
Rikenellaceae
Ruminococcaceae
Streptococcaceae
Tannerellaceae
Day 3 Rep A
Day 3 Rep B
Day 3 Rep C
Day 5 Rep A
Day 5 Rep B
Day 5 Rep C
Day 6 Rep A
Day 6 Rep B
Day 6 Rep C
Day 8 Rep A
Day 8 Rep B
Day 8 Rep C
Day 10 Rep A
Day 10 Rep B
Day 10 Rep C
Day 11 Rep A
Day 11 Rep B
Day 11 Rep C
Day 13 Rep A
Day 13 Rep B
Day 13 Rep C
Day 15 Rep A
Day 15 Rep B
Day 15 Rep C
Donor 87 - AMC
Acidaminococcaceae
Acutalibacteraceae
Akkermansiaceae
Anaerotignaceae
Anaerovoracaceae
Bacteroidaceae
Bifidobacteriaceae
Butyricicoccaceae
CAG-74
Campylobacteraceae
Christensenellaceae
Coprobacillaceae
Coriobacteriaceae
Desulfovibrionaceae
Dialisteraceae
Eggerthellaceae
Enterobacteriaceae_A
Enterococcaceae
GCA-900066905
Lachnospiraceae
Oscillospiraceae_88309
Peptoniphilaceae
Peptostreptococcaceae_256921
Rikenellaceae
Ruminococcaceae
Streptococcaceae
Tannerellaceae
Day 3 Rep A
Day 3 Rep B
Day 3 Rep C
Day 5 Rep A
Day 5 Rep B
Day 5 Rep C
Day 6 Rep A
Day 6 Rep B
Day 6 Rep C
Day 8 Rep A
Day 8 Rep B
Day 8 Rep C
Day 10 Rep A
Day 10 Rep B
Day 10 Rep C
Day 11 Rep A
Day 11 Rep B
Day 11 Rep C
Day 13 Rep A
Day 13 Rep B
Day 13 Rep C
Day 15 Rep A
Day 15 Rep B
Day 15 Rep C
Donor 95 - CTL_R2
Acidaminococcaceae
Acutalibacteraceae
Akkermansiaceae
Anaerotignaceae
Anaerovoracaceae
Atopobiaceae
Bacteroidaceae
Bifidobacteriaceae
Burkholderiaceae_A_595427
Butyricicoccaceae
CAG-74
Christensenellaceae
Clostridiaceae_222000
Coprobacillaceae
Coriobacteriaceae
Desulfovibrionaceae
Dialisteraceae
Eggerthellaceae
Enterobacteriaceae_A
Enterococcaceae
Erysipelotrichaceae
Eubacteriaceae_258449
GCA-900066905
Lachnospiraceae
Marinifilaceae
Methanobacteriaceae
Oscillospiraceae_88309
Peptoniphilaceae
Peptostreptococcaceae_256921
Rikenellaceae
Ruminococcaceae
Streptococcaceae
Tannerellaceae
Veillonellaceae
Day 3 Rep A
Day 3 Rep B
Day 3 Rep C
Day 5 Rep A
Day 5 Rep B
Day 5 Rep C
Day 6 Rep A
Day 6 Rep B
Day 6 Rep C
Day 8 Rep A
Day 8 Rep B
Day 8 Rep C
Day 10 Rep A
Day 10 Rep B
Day 10 Rep C
Day 11 Rep A
Day 11 Rep B
Day 11 Rep C
Day 13 Rep A
Day 13 Rep B
Day 13 Rep C
Day 15 Rep A
Day 15 Rep B
Day 15 Rep C
Donor 95 - AMC
Acidaminococcaceae
Acutalibacteraceae
Akkermansiaceae
Anaerovoracaceae
Bacteroidaceae
Butyricicoccaceae
CAG-74
Christensenellaceae
Clostridiaceae_222000
Coprobacillaceae
Coriobacteriaceae
Desulfovibrionaceae
Eggerthellaceae
Enterobacteriaceae_A
Enterococcaceae
Erysipelotrichaceae
Eubacteriaceae_258449
GCA-900066905
Lachnospiraceae
Methanobacteriaceae
Oscillospiraceae_88309
Peptostreptococcaceae_256921
Rikenellaceae
Ruminococcaceae
Tannerellaceae
Staphylococcaceae
Day 3 Rep A
Day 3 Rep B
Day 3 Rep C
Day 5 Rep A
Day 5 Rep B
Day 5 Rep C
Day 6 Rep A
Day 6 Rep B
Day 6 Rep C
Day 8 Rep A
Day 8 Rep B
Day 8 Rep C
Day 10 Rep A
Day 10 Rep B
Day 10 Rep C
Day 11 Rep A
Day 11 Rep B
Day 11 Rep C
Day 13 Rep A
Day 13 Rep B
Day 13 Rep C
Day 15 Rep A
Day 15 Rep B
Day 15 Rep C
Donor 101 - CTL_R2
Acidaminococcaceae
Acutalibacteraceae
Akkermansiaceae
Anaerotignaceae
Anaerovoracaceae
Bacteroidaceae
Burkholderiaceae_A_595427
Butyricicoccaceae
CAG-74
Christensenellaceae
Coprobacillaceae
Coriobacteriaceae
Desulfovibrionaceae
Eggerthellaceae
Enterobacteriaceae_A
Enterococcaceae
Erysipelotrichaceae
GCA-900066905
Lachnospiraceae
Oscillospiraceae_88309
Peptoniphilaceae
Peptostreptococcaceae_256921
Rikenellaceae
Ruminococcaceae
Tannerellaceae
Veillonellaceae
Staphylococcaceae
Day 3 Rep A
Day 3 Rep B
Day 3 Rep C
Day 5 Rep A
Day 5 Rep B
Day 5 Rep C
Day 6 Rep A
Day 6 Rep B
Day 6 Rep C
Day 8 Rep A
Day 8 Rep B
Day 8 Rep C
Day 10 Rep A
Day 10 Rep B
Day 10 Rep C
Day 11 Rep A
Day 11 Rep B
Day 11 Rep C
Day 13 Rep A
Day 13 Rep B
Day 13 Rep C
Day 15 Rep A
Day 15 Rep B
Day 15 Rep C
Donor 101 - AMC
Acidaminococcaceae
Acutalibacteraceae
Akkermansiaceae
Anaerotignaceae
Anaerovoracaceae
Bacteroidaceae
Butyricicoccaceae
Christensenellaceae
Coprobacillaceae
Coriobacteriaceae
Desulfovibrionaceae
Eggerthellaceae
Enterobacteriaceae_A
Enterococcaceae
Erysipelotrichaceae
GCA-900066905
Lachnospiraceae
Methanobacteriaceae
Oscillospiraceae_88309
Peptostreptococcaceae_256921
Rikenellaceae
Ruminococcaceae
Tannerellaceae
Veillonellaceae
Staphylococcaceae
Day 3 Rep A
Day 3 Rep B
Day 3 Rep C
Day 5 Rep A
Day 5 Rep B
Day 5 Rep C
Day 6 Rep A
Day 6 Rep B
Day 6 Rep C
Day 8 Rep A
Day 8 Rep B
Day 8 Rep C
Day 10 Rep A
Day 10 Rep B
Day 10 Rep C
Day 11 Rep A
Day 11 Rep B
Day 11 Rep C
Day 13 Rep A
Day 13 Rep B
Day 13 Rep C
Day 15 Rep A
Day 15 Rep B
Day 15 Rep C

**Fig. S18**. Frequencies of most represented OTUs per donors during the AMC experiment. Shown are microbiotas variations in the control (CTL_R2) and AMC-treated chambers. Sampling days which have their boundaries in bold indicate the day of maximum perturbation (median perturbation strength for the replicates) observed for a given triplicate within the treatment window (days 5-10).

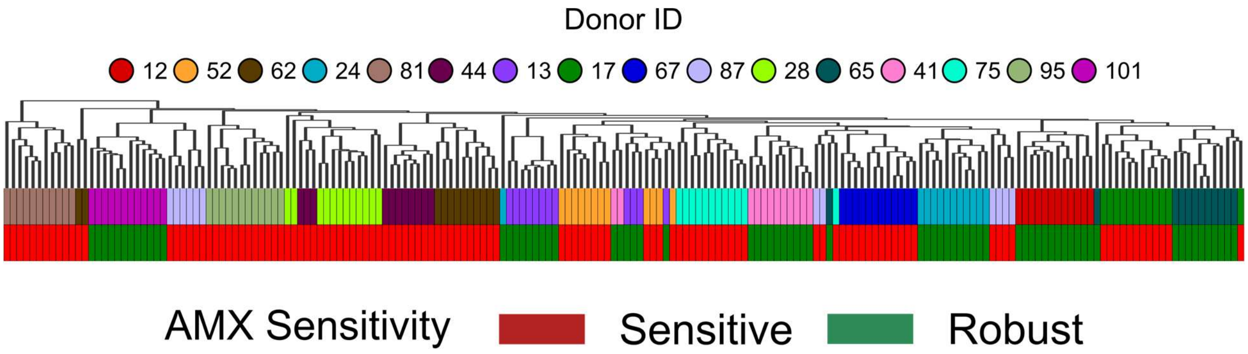


**Fig S19:** Clustermap representing the microbiota composition of each chamber at Day 5, including both the AMX and AMC experiments (12 replicates per donor in total). Hierarchical clustering of the microbiotas was performed using a Jensen-Shannon divergence matrix. Donor IDs are indicated in colors above the heatmap. Donor AMX sensitivity is also indicated in colors above the heatmap, using the classification of Figure 1. The taxonomic rank used for this cluster map is the level 6 (genus). Individual frequencies of genus are not shown for visibility convenience.

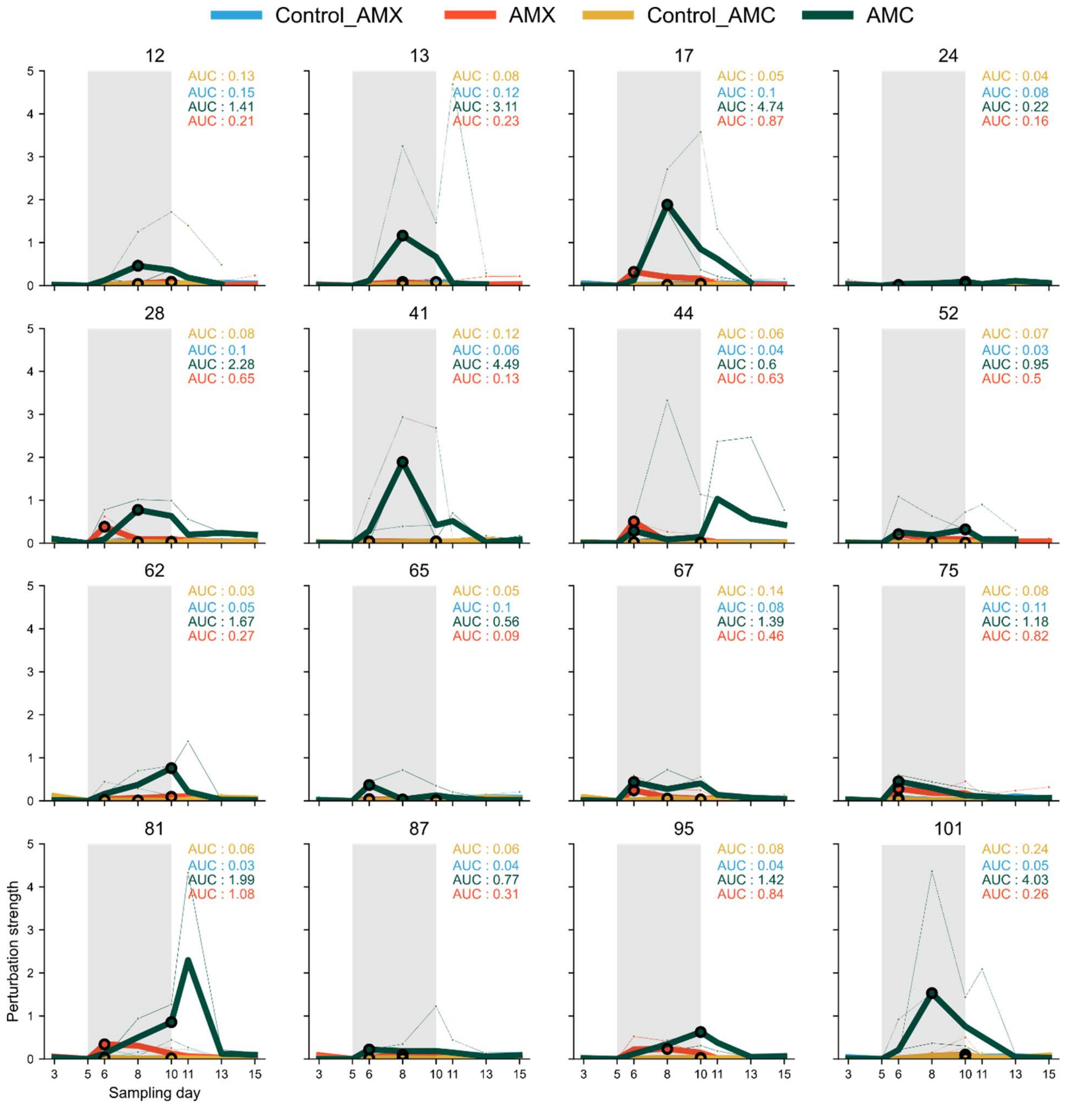


**Fig. S20**. Perturbation observed in each MBRA chamber, given as $|\beta * \alpha|$ metrics against respective Day 5 compositions and α-diversity. For β-diversity, Jensen-Shannon divergence was used, and for α-diversity, Shannon entropy. Thin lines represent each replicate chambers, with red/green lines representing AMX/AMC-treated chambers and blue/ocre lines representing Control chambers. Bold lines are medians of the triplicate chambers. The grey rectangle corresponds to the treatment window (days 5-10). Numbers above each plot refer to the ID of the donors. Area Under Curve (AUC) of the median curves are indicated on each plot. Data shown for Donors 12/24/41/65 are the same as in Fig. 4c, and the same as in Fig. S7 for each donor (AMX experiment).

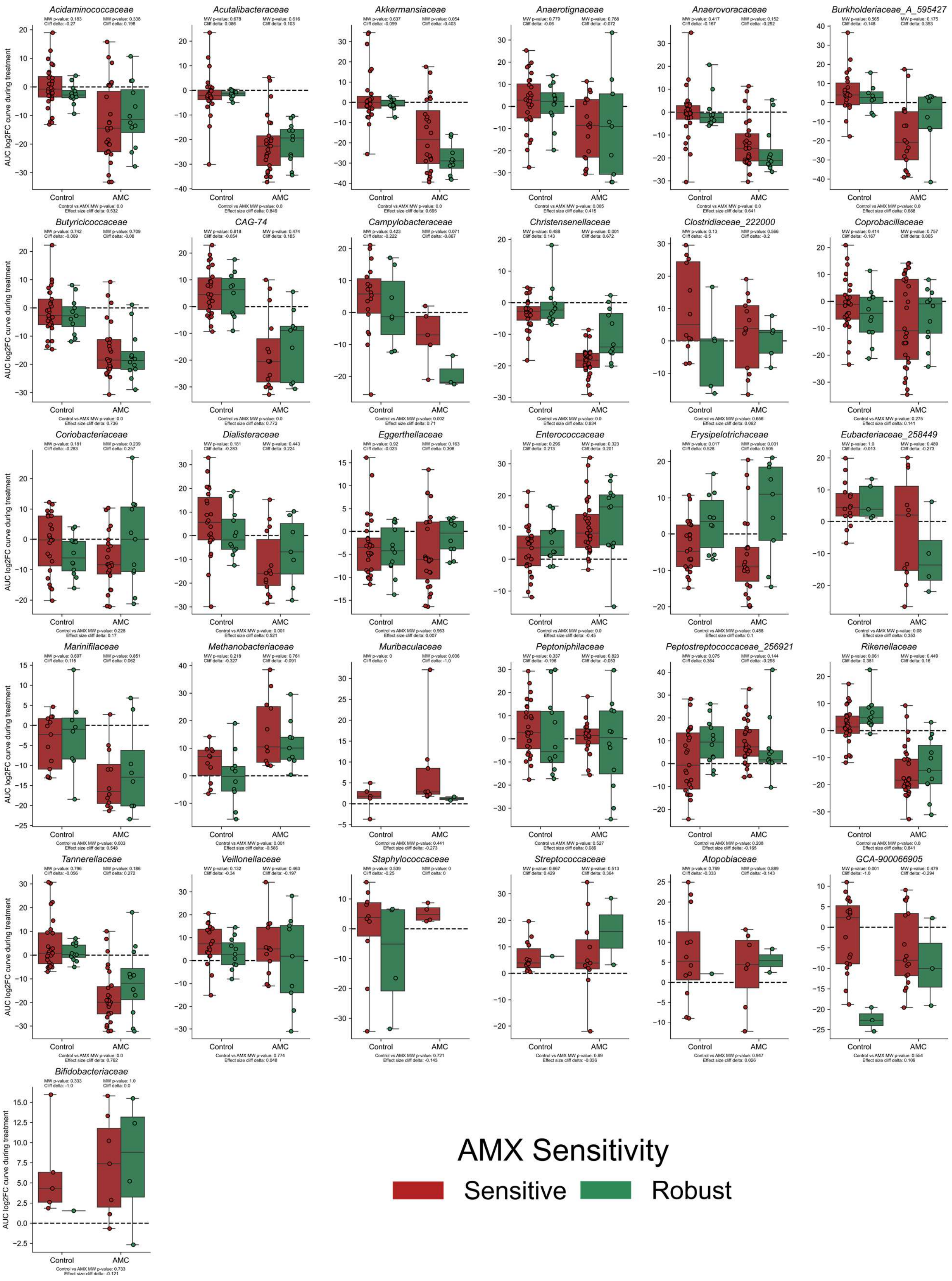
Acidaminococcaceae
Acutalibacteraceae
Akkermansiaceae
Anaerotignaceae
Anaerovoracaceae
Burkholderiaceae_A_595427
Butyricicoccaceae
CAG-74
Campylobacteraceae
Christensenellaceae
Clostridiaceae_222000
Coprobacillaceae
Coriobacteriaceae
Dialisteraceae
Eggerthellaceae
Enterococcaceae
Erysipelotrichaceae
Eubacteriaceae_258449
Marinifilaceae
Methanobacteriaceae
Muribaculaceae
Peptoniphilaceae
Peptostreptococcaceae_256921
Rikenellaceae
Tannerellaceae
Veillonellaceae
Staphylococcaceae
Streptococcaceae
Atopobiaceae
GCA-900066905
Bifidobacteriaceae
AUC log2FC curve during treatment
Control
AMC
AMX Sensitivity
Sensitive
Robust

**Fig. S21**. AUC values of log2FC abundance change of all remaining families (not in Fig. 4f) in chambers either treated with AMC or untreated (control condition). In green are chambers of AMX robust donors, and in red, chambers of AMX sensitive donors. Mann-Whitney (MW) p-values and Cliff's delta effect size are provided, testing the differences between AMX sensitive chambers vs AMX robust chambers in both conditions, and between all chambers treated by AMC or untreated.

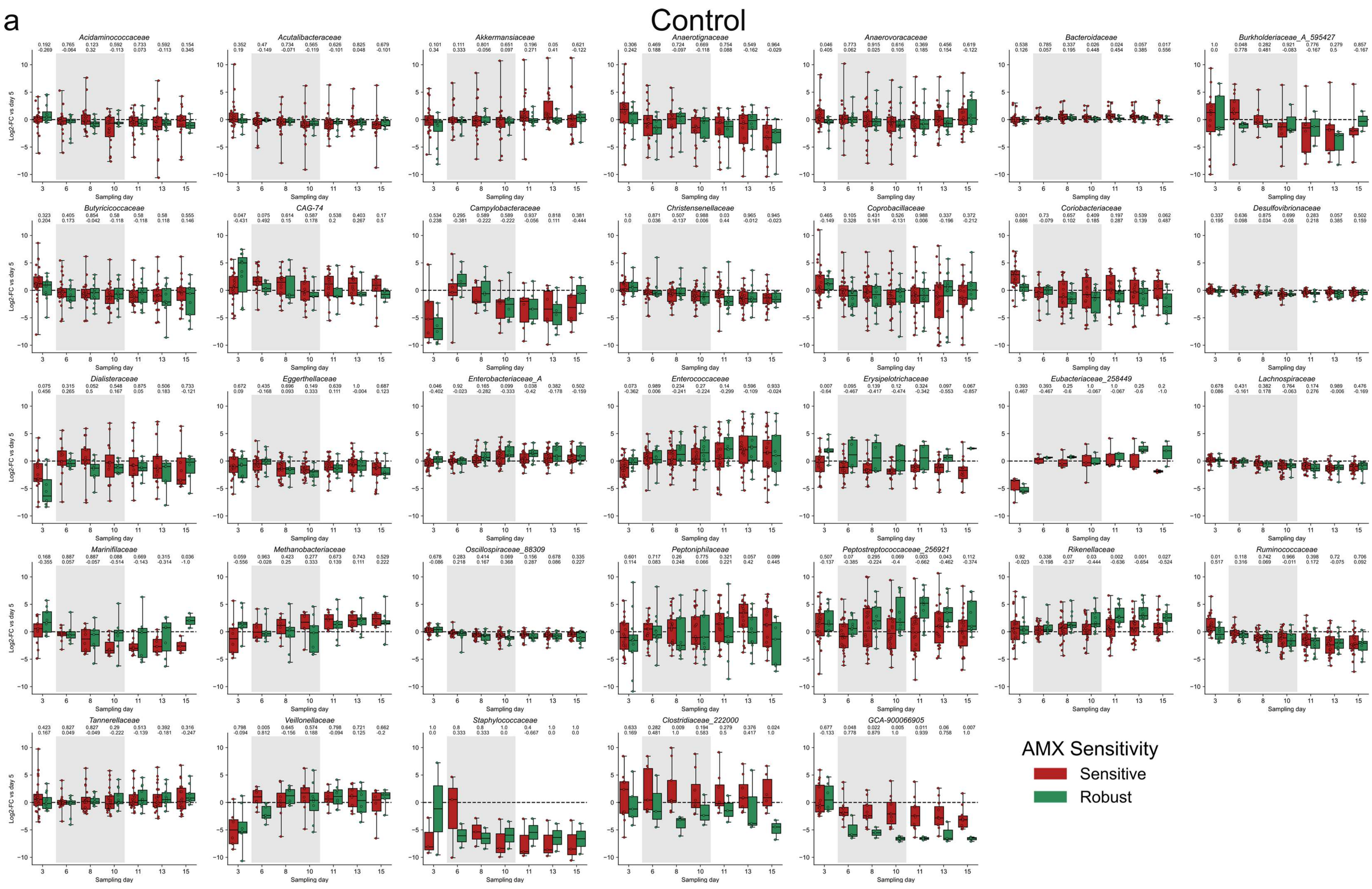
a
Control
Acidaminococcaceae
Acutalibacteraceae
Akkermansiaceae
Anaerotignaceae
Anaerovoracaceae
Bacteroidaceae
Burkholderiaceae_A_595427
Butyricicoccaceae
CAG-74
Campylobacteraceae
Christensenellaceae
Coprobacillaceae
Coriobacteriaceae
Desulfovibrionaceae
Dialisteraceae
Eggerthellaceae
Enterobacteriaceae_A
Enterococcaceae
Erysipelotrichaceae
Eubacteriaceae_258449
Lachnospiraceae
Marinifilaceae
Methanobacteriaceae
Oscillospiraceae_88309
Peptoniphilaceae
Peptostreptococcaceae_256921
Rikenellaceae
Ruminococcaceae
Tannerellaceae
Veillonellaceae
Staphylococcaceae
Clostridiaceae_222000
GCA-900066905
Log2-FC vs day 5
Sampling day
AMX Sensitivity
Sensitive
Robust

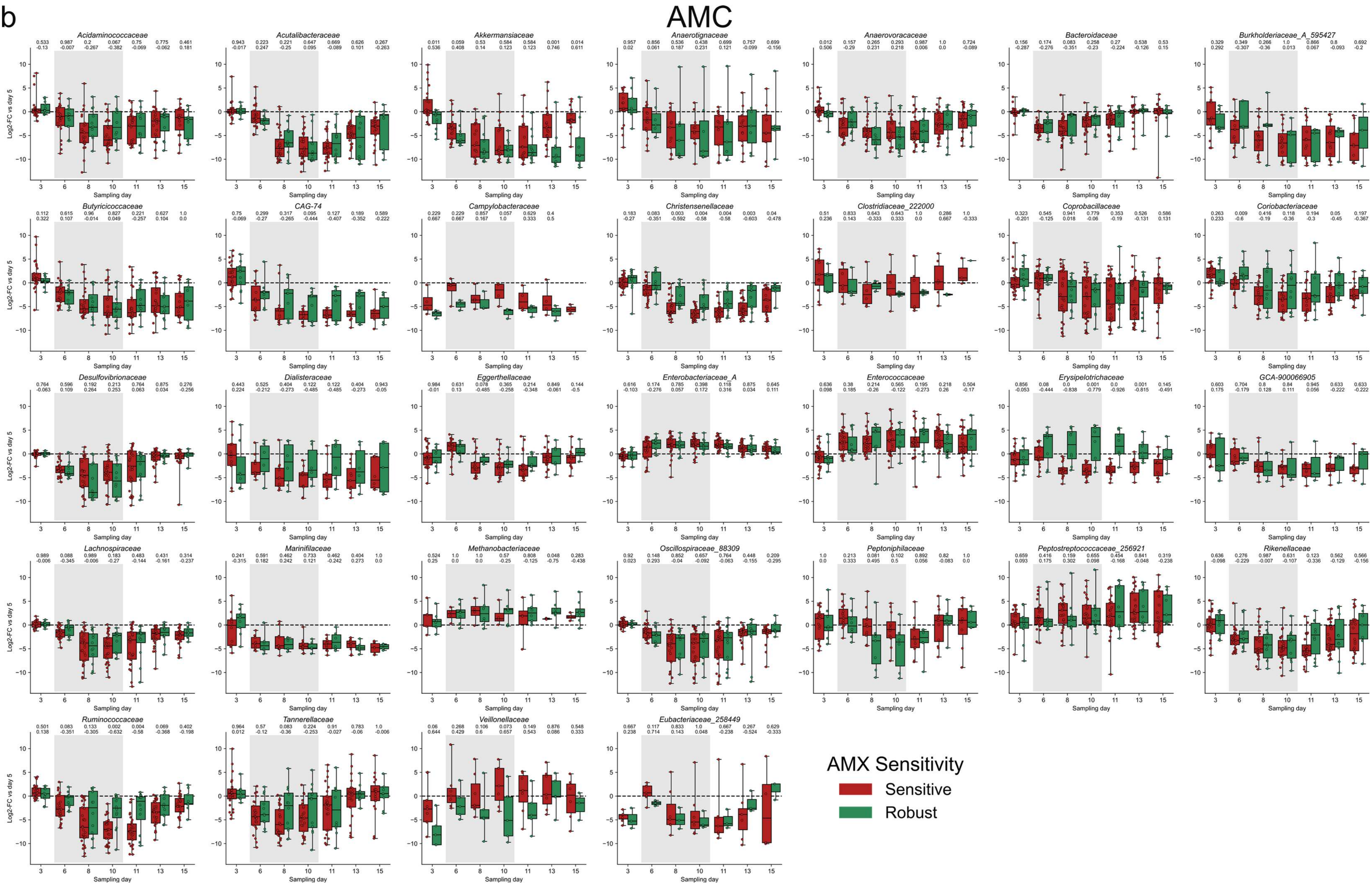
b
AMC
Acidaminococcaceae
Acutalibacteraceae
Akkermansiaceae
Anaerotignaceae
Anaerovoracaceae
Bacteroidaceae
Burkholderiaceae_A_595427
Butyricicoccaceae
CAG-74
Campylobacteraceae
Christensenellaceae
Clostridiaceae_222000
Coprobacillaceae
Coriobacteriaceae
Desulfovibrionaceae
Dialisteraceae
Eggerthellaceae
Enterobacteriaceae_A
Enterococcaceae
Erysipelotrichaceae
GCA-900066905
Lachnospiraceae
Marinifilaceae
Methanobacteriaceae
Oscillospiraceae_88309
Peptoniphilaceae
Peptostreptococcaceae_256921
Rikenellaceae
Ruminococcaceae
Tannerellaceae
Veillonellaceae
Eubacteriaceae_258449
Log2-FC vs day 5
Sampling day
AMX Sensitivity
Sensitive
Robust

**Fig S22**. Day-to-day log2FC abundance change in all families in chambers either untreated (control condition - b) or treated with AMC (b). In green are chambers of AMX-Robust donors, and in red chambers of AMX-Sensitive donors. Above each sampling day, Mann-Whitney p-values (top) and Cliff's delta effect sizes (bottom) are provided, testing the differences between AMX-Sensitive chambers vs AMX-Robust chambers at a given sampling day.

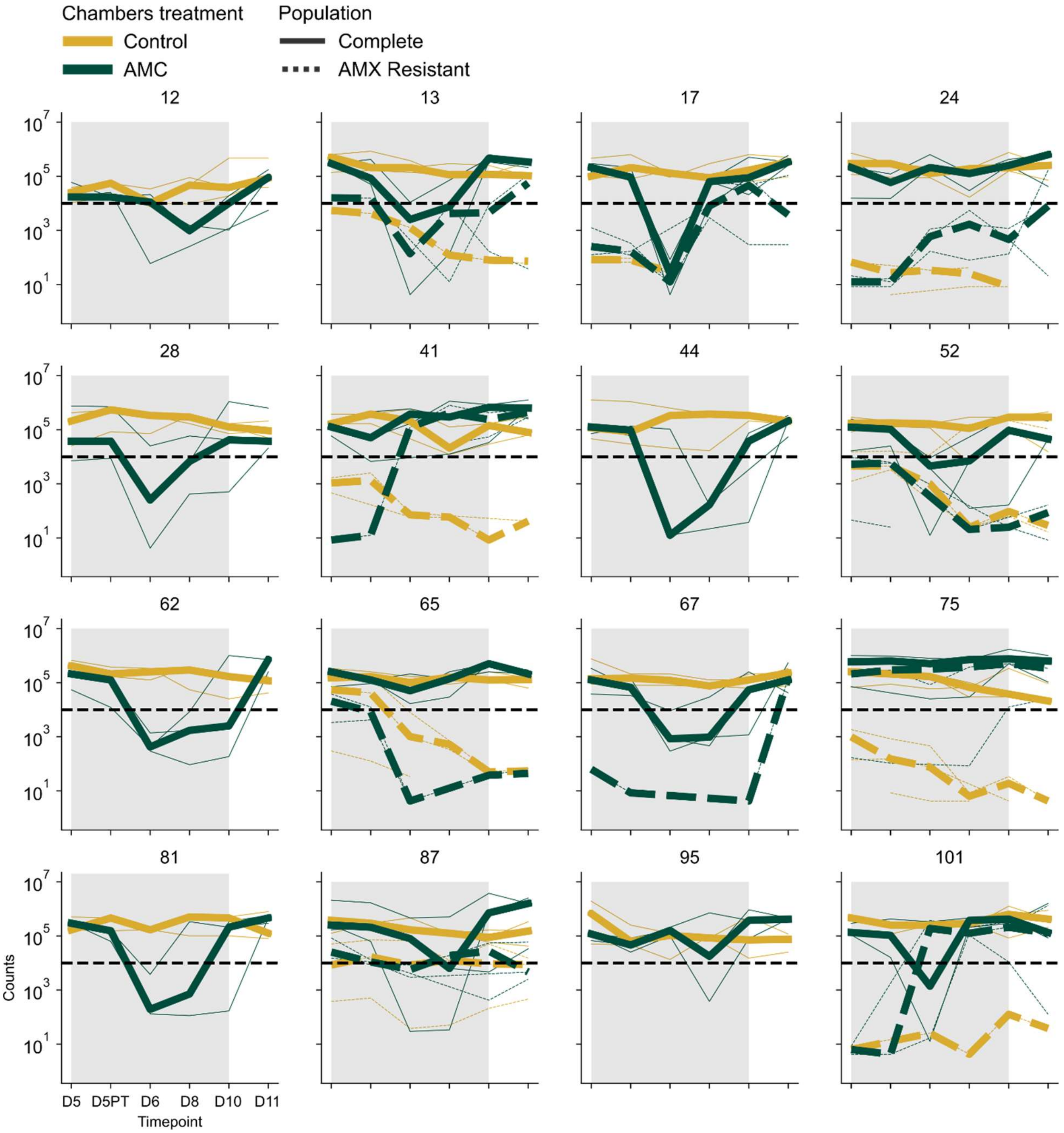


**Fig. S23**. Change in *Enterobacteriaceae* population counts over time in each MBRA chamber (AMC experiment). For each donor, populations from untreated (ochre) or AMC-treated (dark green) chambers are shown. Since populations are plated on Drigalski agar with or without AMX, we could also count the total population (full lines) or AMX-resistant population (dashed lines) in each chamber. For each triplicate, replicates are shown as thin lines and median is shown as a bold line. The detection limit of $10^4$ is shown as a dark bold dashed line.

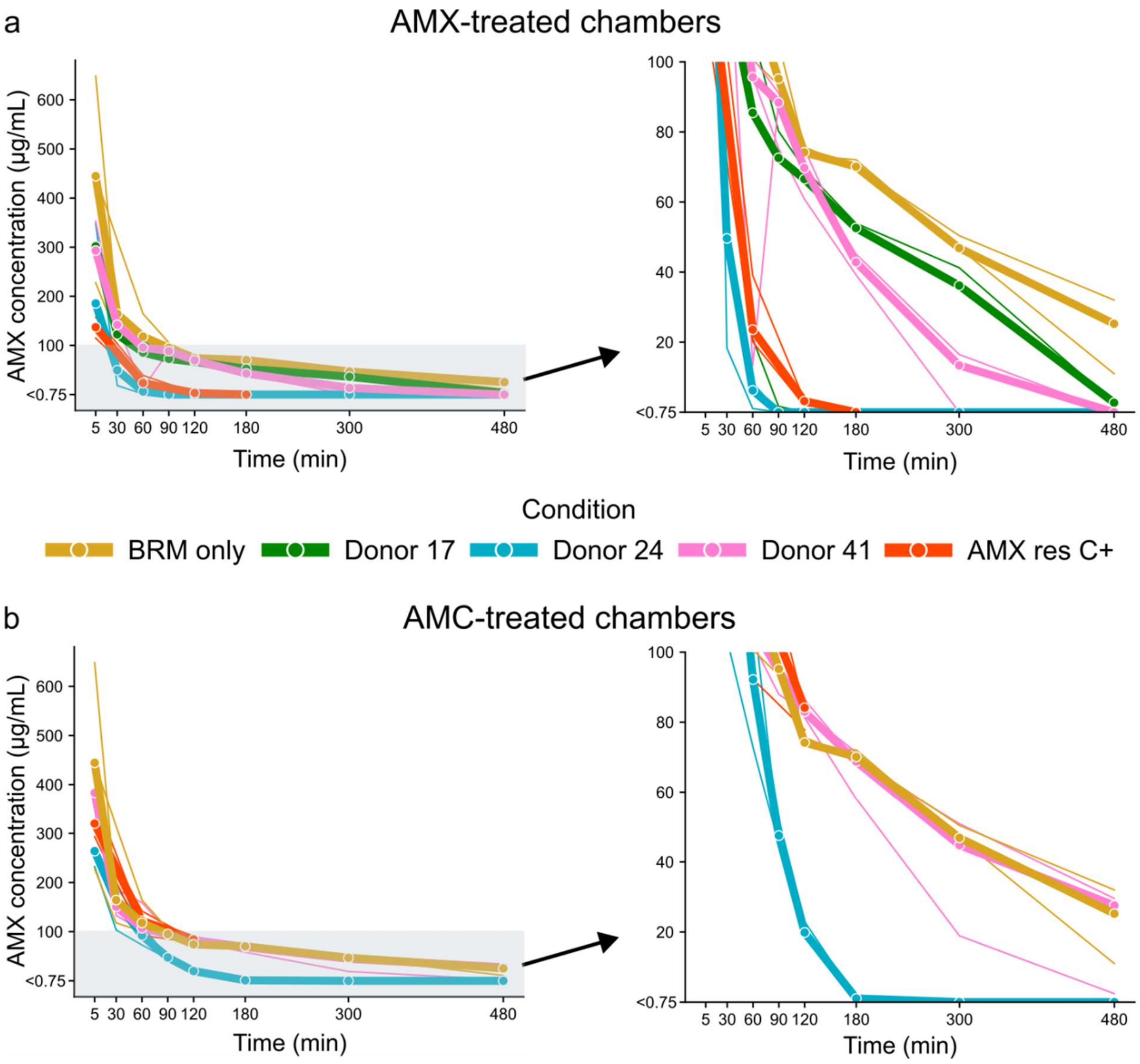


**Fig. S24**. Absolute AMX concentrations as determined by mass spectrometry, given in µg/mL. Bold lines represent the median of the triplicates which are individually shown as thin lines. On top are chambers treated with AMX (a) and at the bottom are chambers treated with AMC (b). Left graphs show the full data and right graphs are zooms of the 0.75-100 µg/mL concentrations windows for clarity purpose. The BRM only control chambers, treated solely with AMX, are shown at the bottom for comparison purposes (same data on both (a) and (b) graphs). The concentration of 0.75 µg/mL was the detection limit of AMX.

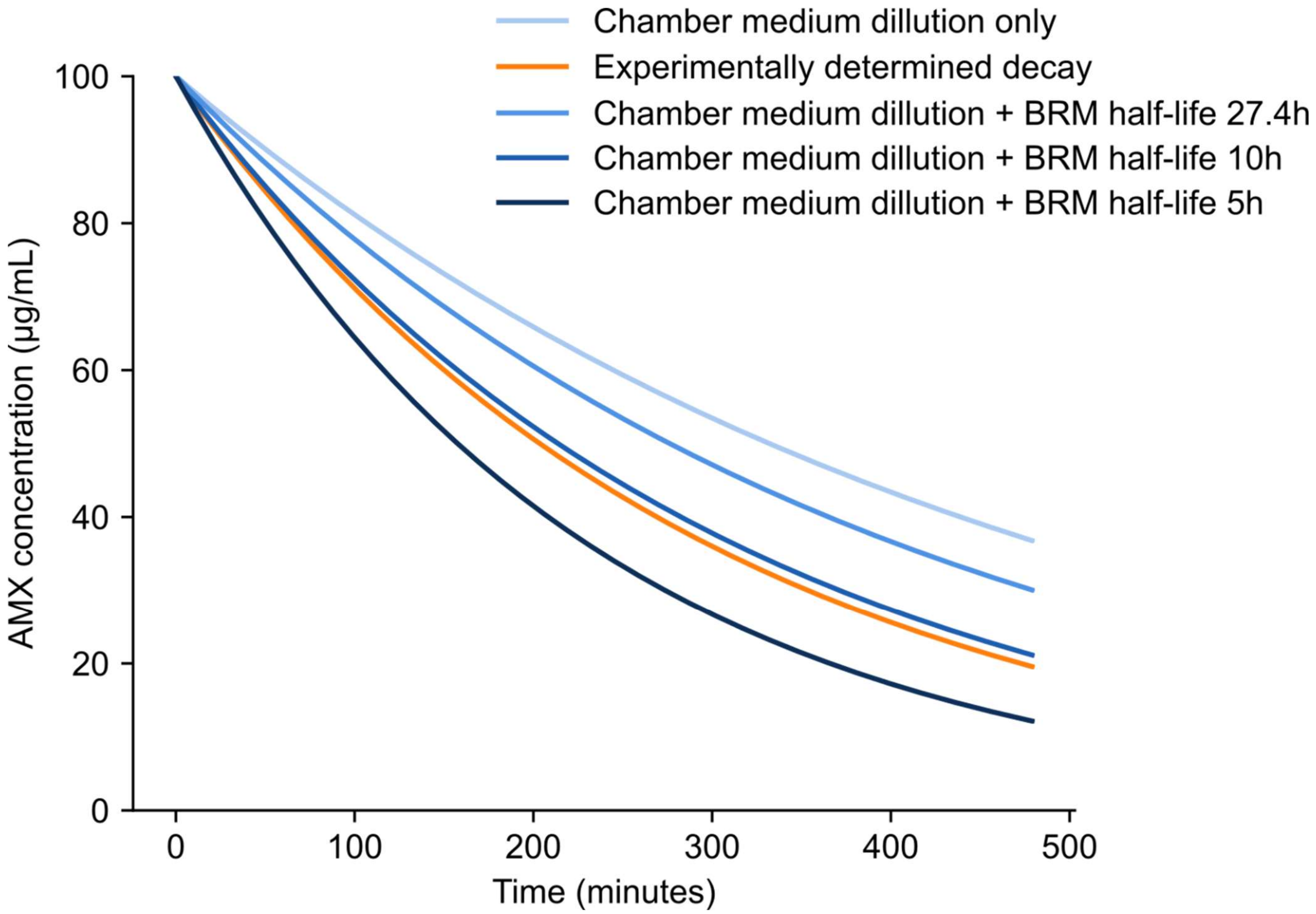


**Fig. S25**. Modelisation of AMX concentrations (µg/mL) over time in MBRA chambers, with a starting concentration of 100 µg/mL and various decay constants λ determined according to multiple AMX half-lives. Parameters taken into account are medium dilution in MBRA chambers (1.875 mL/h) as well as stability of AMX in BRM medium, with 27.4h having previously been reported as half-life of AMX in Mueller-Hinton broth (See supplementary note 6). The BRM-only decay constant was obtained from our dataset and is the same as the one being used for Fig. S26 and Fig. 5.

a

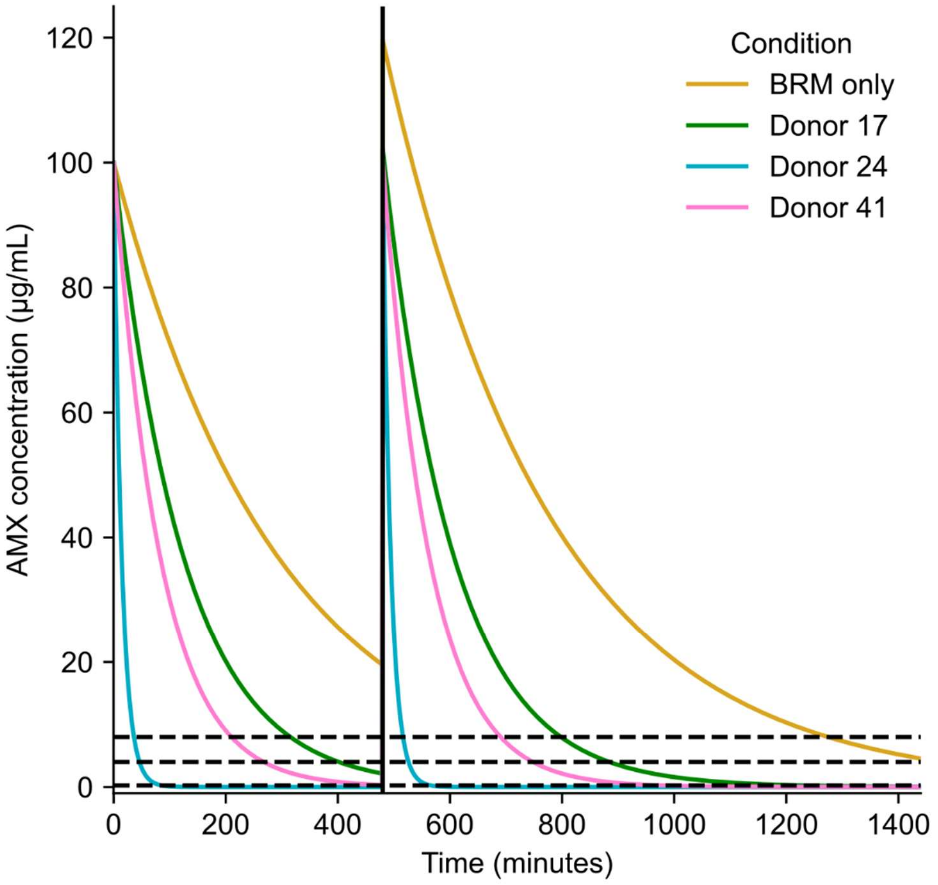

120
100
80
60
40
20
0
AMX concentration (µg/mL)
0
200
400
600
800
1000
1200
1400
Time (minutes)
Condition
BRM only
Donor 17
Donor 24
Donor 41


b

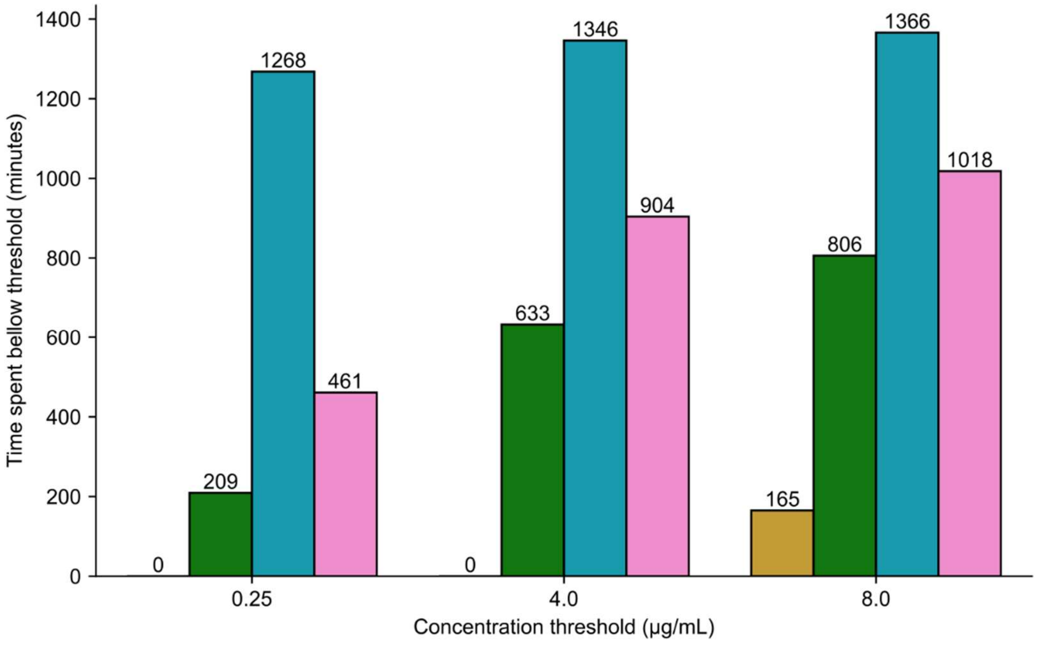

1400
1200
1000
800
600
400
200
0
Time spent bellow threshold (minutes)
0
209
1268
461
0
633
1346
904
165
806
1366
1018
0.25
4.0
8.0
Concentration threshold (µg/mL)

**Fig. S26**. Modelisation of AMX concentrations (µg/mL) in our MBRA chambers during our 8-16h treatment regimen. **a**. Concentrations of AMX modelled using the experimentally determined decay constants λ for each donor and the BRM-only condition (as in Fig. S25). A starting concentration of 100 µg/mL is used. At Time=480 min, AMX concentrations are supplemented by 100 µg/mL. Dashed lines correspond to three AMX concentrations thresholds, 8, 4, and 0.25 µg/mL. **b**.Time spent (minutes) in each chamber under various AMX concentration thresholds (µg/mL), determined using the model in **a**.

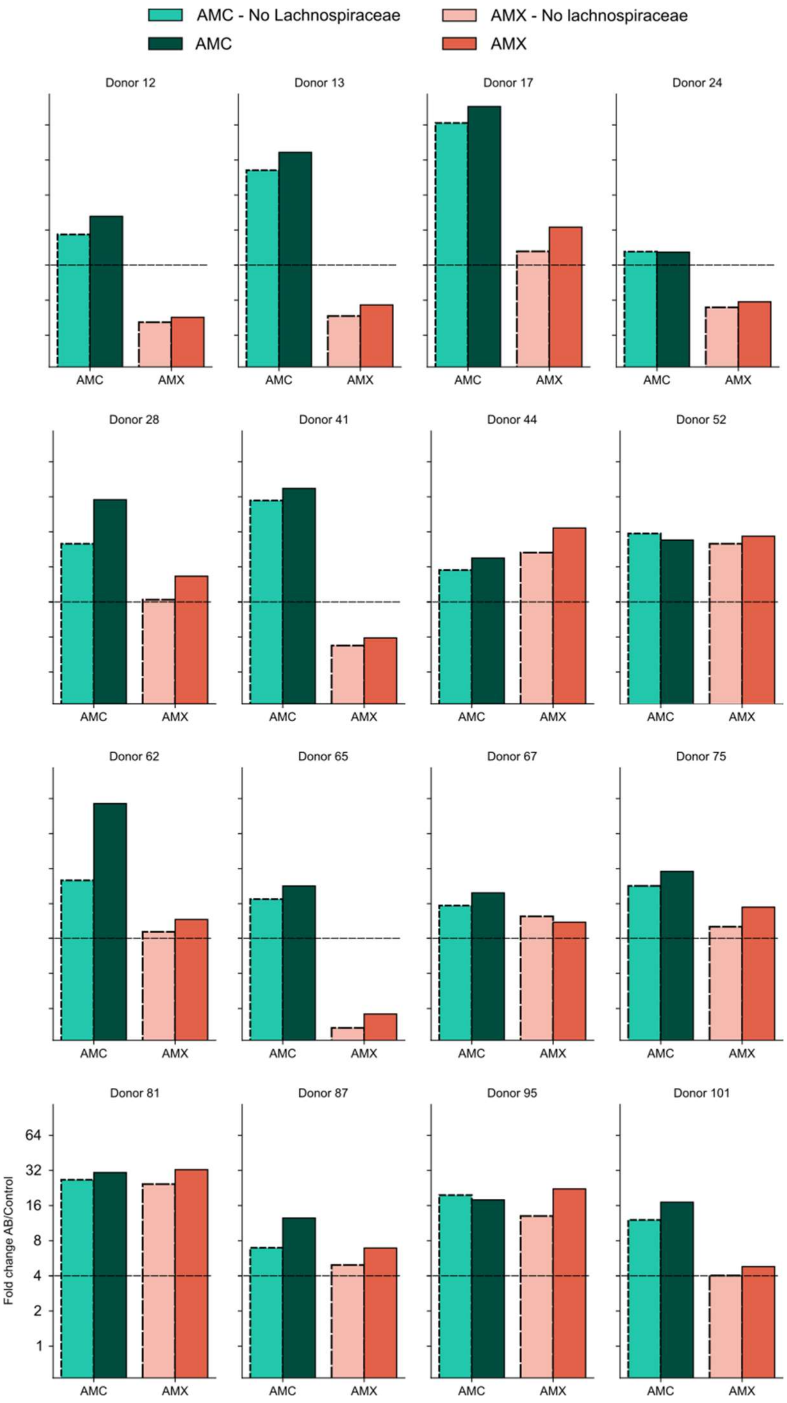

**Fig. S27.** Perturbation strength measured in the 16 donors in the Amoxicillin (AMX - red) and AMX-clavulanic acid (AMC – blue green) experiments. Shown are perturbations in microbiotas with *Lachnospiraceae* (darker full bars) or without *Lachnospiraceae* (light dashed bars). The later was obtained through the removal of *Lachnospiraceae* frequencies in the microbiotas and adjustments of frequencies of the other families accordingly.

| Donor ID | Gender | Age |
|---|---|---|
| 12 | Female | 50 |
| 13 | Female | 57 |
| 17 | Male | 78 |
| 24 | Female | 51 |
| 28 | Female | 50 |
| 41 | Female | 73 |
| 44 | Female | 59 |
| 52 | Female | 46 |
| 62 | Female | 51 |
| 65 | Female | 36 |
| 67 | Female | 55 |
| 75 | Female | 57 |
| 81 | Female | 39 |
| 87 | Female | 37 |
| 95 | Female | 52 |
| 101 | Female | 47 |

Table S1: Description of the 16 donors panel from the NutrinetSanté cohort.
